# The moulding of intra-specific trait variation by selection under ecological inheritance

**DOI:** 10.1101/2022.12.26.521924

**Authors:** Iris Prigent, Charles Mullon

## Abstract

Organisms continuously modify their environment, often impacting the fitness of future conspecifics due to ecological inheritance. When this inheritance is biased towards kin, selection favours modifications that increase the fitness of downstream individuals. How such selection shapes trait variation within populations remains poorly understood. Using mathematical modelling, we investigate the coevolution of multiple traits in a group-structured population when these traits affect the group environment, which is then bequeathed to future generations. We examine when such coevolution favours polymorphism as well as the resulting associations among traits. We find in particular that two traits become associated when one trait affects the environment while the other influences the likelihood that future kin experience this environment. To illustrate this, we model the coevolution of (a) the attack rate on a local renewable resource, which deteriorates environmental conditions, with (b) dispersal between groups, which reduces the likelihood that kin suffers from such deterioration. We show this often leads to the emergence of two highly-differentiated morphs: one that readily disperses and depletes local resources; and another that maintains these resources and tends to remain philopatric. More broadly, we suggest that ecological inheritance can contribute to phenotypic diversity and lead to complex polymorphism.

## 1 Introduction

By consuming, polluting or engineering, most if not all organisms modify and transform the environment they live in. Via such modifications, an individual impacts its own fitness as well as the fitness of conspecifics who share its environment. These fitness effects can extend to future generations when environmental modifications are transmitted to offspring under what is referred to as ecological inheritance (Odling-Smee, 1988; Laland et al., 1996; Odling-Smee et al., 1996, 2003; Bonduriansky, 2012; Bonduriansky and Day, 2020). For example, many plants continuously modify their substrate via plant-soil feedbacks that can stretch to downstream generations (Ehrenfeld et al., 2005; e.g. by producing and absorbing tannin, Kraus et al., 2003). *Pseudomonas aeruginosa* release long-lasting iron-scavenging siderophores, thus benefiting close-by conspecifics in the short and long term (Ratledge and Dover, 2000; Imperi et al., 2009), including individuals that are not living yet (Kümmerli and Brown, 2010). Ecological inheritance can of course also be harmful, for instance when individuals over-consume a slowly renewable resource or release pollutants that are difficult to degrade.

For natural selection to shape the inter-generational ecological effects of a trait, the genes underlying this trait must be statistically associated to the environment they transform (Dawkins, 1982, 2004; Brodie, 2005; Lehmann, 2007, 2008). This association entails that an environmental modification is more likely to be experienced by individuals in the future that carry the same genes as the individual who caused the initial modification. One simple and ubiquitous way for a gene-environment association to emerge is via spatial structure. When the population is subdivided and dispersal between subpopulations is limited, individuals in the same local environment are more likely to share the same genes than individuals sampled at random in the population (i.e. they are related; Hamilton, 1964; Rousset, 2004). This is true of individuals living at the same but also at different generations (Lehmann, 2007). As a result, the inter-generational ecological modification made by an individual preferentially affects the fitness of its future relatives when dispersal is limited (Lehmann, 2008).

How directional selection steers the gradual evolution of traits with inter-generational ecological effects under limited dispersal has been well-studied (Lehmann, 2007, 2008; Sozou, 2009; Mullon and Lehmann, 2018; Arnoldi et al., 2020; Mullon et al., 2021). One of the main insights from this theory is that populations in which dispersal is more limited are more likely to evolve traits that are costly to the individual but yield delayed ecological benefits, such as the preservation of a common good (Silver and Di Paolo, 2006; Lehmann, 2007, 2008; Sozou, 2009; Krakauer et al., 2009; Mullon and Lehmann, 2018; Arnoldi et al., 2020; Mullon et al., 2021; for review: Estrela et al., 2019). But while directional selection can explain trait variation between species (or completely isolated populations), it is not sufficient to investigate variation within species, which requires characterising disruptive selection (Rousset, 2004; Dercole and Rinaldi, 2008). Consequently, these previous studies focused on directional selection do not address the question of how selection can favour the emergence of polymorphism in traits with inter-generational ecological effects.

Intra-specific variation in ecologically-relevant traits is nevertheless common, with potentially significant ecological effects (Bolnick et al., 2003; Araújo et al., 2011; Bolnick et al., 2011; Violle et al., 2012; Des Roches et al., 2018). Variation in predator traits can, for example, stabilise prey-predator dynamics (Okuyama, 2008), or lead to apparent mutualism between preys (Schreiber et al., 2011). So far, mathematical models that investigate the emergence of intra-specific variation owing to disruptive selection under limited dispersal assume that traits have immediate effects, which do not carry over between generations (e.g., Day, 2001; Ajar, 2003; Wakano and Lehmann, 2014; Mullon et al., 2016; Ohtsuki et al., 2020; Schmid et al., 2022). However, simulations indicate that polymorphism in traits that have lasting ecological effects can emerge in spatially-structured population (Silver and Di Paolo, 2006; Han et al., 2006; Behar et al., 2014; Joshi et al., 2020). In such a population where harvesting of a common good evolves through social learning, agent-based simulations show for instance that different harvesting strategies can coexist when learning is fast relative to the renewal of the good, suggesting a role for ecological inheritance (Joshi et al., 2020). More broadly, these simulations offer useful but partial insights into how intra-specific variation is moulded under ecological inheritance and limited dispersal.

In this paper, we extend current theory to better understand when gradual evolution leads to polymorphism in traits that have long-lasting environmental effects through disruptive selection. We investigate mathematically the coevolution of multiple traits in a group-structured population when these traits affect the group environment, which is then passed down to future generations. We use our model to investigate the type of between-traits within-individuals correlations that are favored by selection when polymorphism emerges. Our analyses reveal in particular that two traits tend to be correlated when one modifies the environment in a long-lasting manner while the other influences the likelihood that future relatives experience this environmental modification. To illustrate this, we model the coevolution of the attack rate on a local renewable resource, which deteriorates environmental conditions, with dispersal, which reduces the likelihood that relatives suffer from such deterioration. Beyond this specific example, we discuss the other pathways revealed by our model via which selection favours trait associations under ecological inheritance, with potential implications for dispersal and behavioural syndromes, phenotypic plasticity, and niche construction.

## 2 Model and Methods

### 2.1 Life-cycle, traits and environmental dynamics

We consider a population of haploids distributed among a large number of patches. All patches carry the same number *N* of individuals and are uniformly connected by dispersal (according to the island model, Wright, 1931). Each patch is characterised by an environmental state or ecological variable *ϵ* ∈ **ℝ** (e.g. abundance of a resource, pollution level, quality of a common good), and each individual by a phenotype ***z*** = (*z*_1_, …, *z*_*n*_) ∈ **ℝ**^*n*^ made of *n* genetically determined quantitative traits, where any trait *z*_*p*_ (*p* ∈ {1,…, *n*}, referred to as “trait *p*” for short) can influence the state of the patch (e.g. attack rate on a resource, production of a pollutant, investment into a common good). We census this population at discrete time points between which the following occurs (Fig. 1A for diagram): (i) within patches, individuals interact with one another and with their environment whose state *ϵ* can change as a result (we specify such interactions and how they depend on traits below); (ii) individuals reproduce, producing a large number of clonal offspring (large enough to ignore demographic stochasticity), and then die; (iii) independently of one another, each offspring either disperses to a randomly chosen patch or remains in its natal patch; and finally (iv) offspring compete locally in each patch for *N* spots. Patch state *ϵ* and individual traits ***z*** can influence any vital rate, such as fecundity, offspring dispersal, or survival (or combination thereof). In addition, patch regulation may occur before and after dispersal, allowing for selection to be soft or hard (Wallace, 1975; Christiansen, 1975; Débarre and Gandon, 2011), as long as all patches carry *N* mature individuals by the end of the life-cycle. Generations are non-overlapping but as we detail next, individuals of different generations can interact with one another indirectly via the environment.

**Figure 1:**
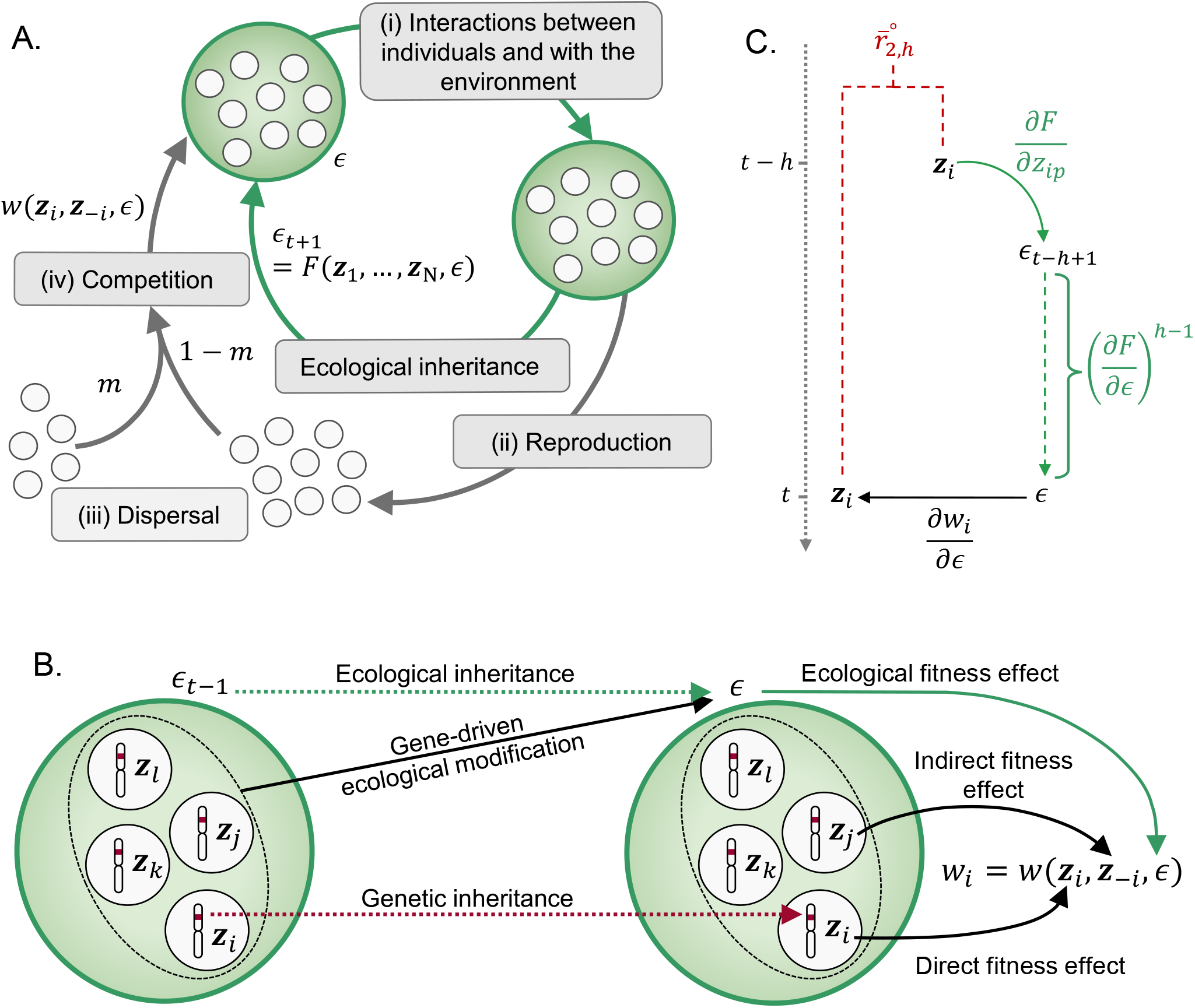
**A. Life-cycle**, section 2.1 for details. **B. Genetic and ecological inheritance**. Diagram showing the dual pathways of inheritance in the model, where individuals inherit their genes from their parent (“Genetic inheritance”) but also their local environment (“Ecological inheritance”), which has been modified by the genetically-determined traits of the parental generation (“Gene-driven ecological modification” with ***z***_*i*_ for trait of individual *i*). The fitness *w*_*i*_ of a focal individual *i* then depends on its own traits ***z***_*i*_ (“direct fitness effect”), the traits of its neighbours ***z***_−*i*_ (“indirect fitness effect”), and on the state of the environment *ϵ* (“ecological fitness effect”). **C. Diagram for the expected effect of a change in trait** *p* **on the fitness of an individual in the future via ecological inheritance**, illustrating eq. (11). A change in trait *p* in an individual at generation *t* − *h* modifies the environment of the next generation *t* − *h* + 1 by *∂F* /*∂z*_*ip*_ (solid green arrow). Due to ecological inheritance, this modification carries over downstream generations till generation *t* (according to a factor (*∂F* /*∂ϵ*)^*h*−1^, dashed green arrow). This in turn influences the fitness of a focal individual living at that time according to *∂w*_*i*_ /*∂ϵ* (solid black arrow). These two individuals separated by *h* generations belong to the same lineage (and thus express the same phenotype ***z***_*i*_) with probability 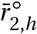 (dashed red lines).

To allow for individuals to transform their environment in a way that can be passed onto future generations, we write the state *ϵ*_*t* +1_ of a patch at a generation *t* + 1 as a function of the traits expressed by individuals in that patch at the previous generation *t*, as well as the previous state; specifically as

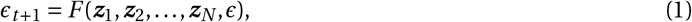

where the vector ***z***_*i*_ denotes the phenotype expressed by individual *i* ∈ {1,…, *N* } living in the patch at generation *t* and *ϵ* is the state of the patch at generation *t* (Fig. 1B for diagram). The environmental dynamics given by the map *F* (eq. 1) unfold even in the absence of genetic variation (i.e. when ***z***_*i*_ = ***z*** for all *i*). We assume that these dynamics converge to a stable equilibrium, meaning that in a population monomorphic for ***z*** (i.e. where all individuals in the population express the same phenotype ***z***), all patches are eventually characterised by the same environmental equilibrium, which we denote by 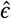 (this equilibrium will typically depend on ***z*** but we do not write this dependence explicitly for short). This equilibrium satisfies

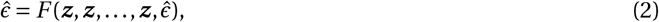

as well as the stability condition,

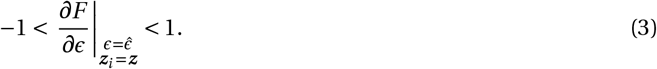

Here and hereafter, all derivatives are estimated where all individuals express the same phenotype ***z*** and all patches are characterised by the associated ecological equilibrium 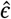. So all derivatives should be seen as functions of the evolving traits ***z***. The quantity *∂F*/*∂ϵ* gives the effect of a perturbation in the state of a patch at one generation on the state at the next generation. This *∂F* /*∂ϵ* can thus be thought of as the effect of ecological inheritance: the greater the absolute value of *∂F* /*∂ϵ* is, the more consequential an environmental modification is to future generations.

The consequences of an environmental modification also depend on how the fitness of an individual varies with its local environment. To capture this in a general way, we assume the fitness *w*_*i*_ of individual *i* ∈ {1,…, *N* } from a focal patch can be written as a function,

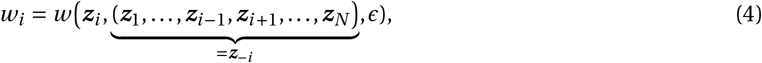

which gives the expected number of offspring produced by this individual *i* that are recruited across all patches (so counted over one full iteration of the life-cycle, from steps i-iv of section 2.1), given: (i) its phenotype, ***z***_*i*_ = (*z*_*i* 1_, …, *z*_*in*_) (with *z*_*ip*_ the value expressed for trait *p*); (ii) the phenotypes expressed by its *N* − 1 patch-neighbours, collected in ***z***_−*i*_ = (***z***_1_,…, ***z***_*i* −1_, ***z***_*i* +1_,…, ***z***_*N*_); and (iii) the environmental state of its patch, *ϵ*. This formulation allows for the fitness of an individual to depend on interactions between its own traits, the traits of its neighbours, and the environment (Fig. 1B). Since there is no class structure in our model (e.g. no age, sex or stage structure), the fitness function is invariant to permutations within ***z***_−*i*_, i.e. it does not matter which specific neighbours express which phenotype, only the collection of phenotypes matters. Fitness may however depend on the number of such neighbours and thus on patch size, *N* (eq. 33 for a specific example). Note also that even though traits ***z*** are genetically determined, the general formulation for environmental dynamics (eq. 1) and individual fitness (eq. 4) allows us to consider the evolution of plastic traits through reaction norms (as in e.g. Lande, 2009). For instance, one could assume that the expression of a plastic trait is given by *z*_1_ + *z*_2_*ϵ* and then investigate the joint evolution of *z*_1_ and *z*_2_, where *z*_2_ captures plasticity to the local environment *ϵ*.

### 2.2 Evolutionary dynamics

We are interested in the coevolution of the *n* traits, and particularly in whether such coevolution leads to polymorphism. To investigate this, we assume that traits evolve via the input of rare genetic mutations with weak phenotypic effects. Under these assumptions, evolutionary dynamics take place in two steps (Metz et al., 1995; Rousset, 2004; Dercole and Rinaldi, 2008), which can be inferred from the invasion fitness (i.e. the geometric growth rate), *W* (***ζ, z***), of a rare allele coding for a deviant phenotype ***ζ*** = (*ζ*_1_, …, *ζ*_*n*_) in a population otherwise monomorphic for ***z*** = (*z*_1_, …, *z*_*n*_) (Tuljapurkar, 1989; Ferriere and Gatto, 1995; Tuljapurkar et al., 2003; Otto and Day, 2007). Our general approach, which is based on a sensitivity analysis of *W* (***ζ, z***), is detailed in Appendix A. We summarize here how such an analysis can be used to understand how selection leads to polymorphism under ecological inheritance.

#### 2.2.1 Directional selection

First, the population evolves under directional selection whereby selected mutants rapidly sweep the popula-tion before a new mutation arises, so that the population can be thought of as “jumping” from one monomorphic state to another. Trait dynamics during this phase are characterised by the selection gradient vector,

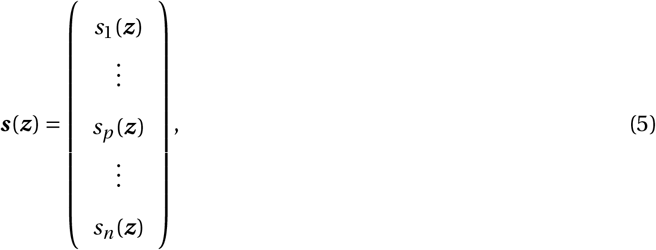

which points in the direction favoured by selection in phenotypic space (the space of all possible phenotypes, here **ℝ**^*n*^ or a subset thereof), i.e.

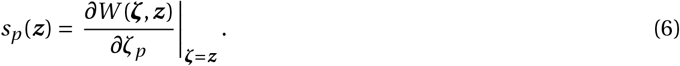

Specifically, selection favours an increase in trait *p* when *s*_*p*_ (***z***) > 0, and conversely a decrease when *s*_*p*_ (***z***) < 0. The population may thus eventually converge to a singular phenotype, 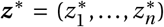, which is such that each entry of the selection gradient is zero at ***z***^∗^, i.e.

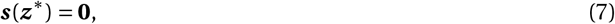

where **0** is a vector of zeroes of size *n*. For the population to converge to ***z***^∗^, it is sufficient that the Jacobian matrix,

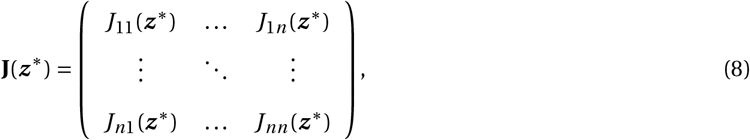

with (*p, q*) entry

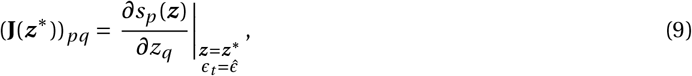

is negative-definite (i.e. the symmetric real part of **J**(***z***^∗^), [**J**(***z***^∗^) + **J**(***z***^∗^)^T^]/2, has only negative eigenvalues, Leimar, 2005, 2009; Débarre et al., 2014; Geritz et al., 2016). Such a phenotype is an attractor of evolutionary dynamics and typically referred to as (strongly) convergence stable (Leimar, 2009).

#### 2.2.2 Directional selection under ecological inheritance

The selection gradient ***s***(***z***) (eq. 6) is defined in terms of the invasion fitness *W* (***ζ, z***) of a genetic mutant, which can be seen as a measure of fitness at the level of the gene that codes for a deviant phenotype. To reveal directional selection on the inter-generational effects of traits through ecological inheritance requires expressing ***s***(***z***) in terms of *individual* (or personal) fitness, i.e. in terms of *w*_*i*_ (eq. 4, e.g. Lehmann, 2007, 2008; Mullon and Lehmann, 2018). We briefly go over these previous findings in the context of our model here.

The selection gradient on a trait *p* with inter-generational effects (given by eq. 1) can be expressed in terms of individual fitness (eq. 4) as

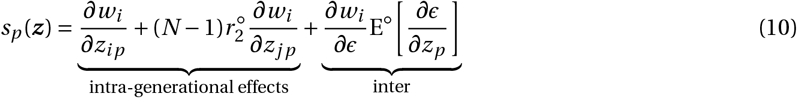

(eqs. 13-15 in Mullon and Lehmann, 2018, eq. 2 in Lehmann, 2007, eq. 4 in Lehmann, 2008 and eq. 4.11 in Sozou, 2009 – appendix B here for a re-derivation of this result). Eq. (10) consists of three weighted fitness effects. First, *∂w*_*i*_ /*∂z*_*ip*_ is the effect of a change in trait *p* in the focal individual *i* on its own fitness (direct fitness effect). Second, *∂w*_*i*_ /*∂z*_*jp*_ is the effect of a change in trait *p* in a patch-neighbour (individual *j* ≠ *i*) on focal fitness (indirect fitness effect). This is weighted by the number (*N* − 1) of such neighbours, and the pairwise relatedness coefficient under neutrality, 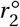, which is the probability that a randomly sampled neighbour is identical-by-descent to a focal. Throughout, quantities with a superscript ◦ are evaluated under neutrality, i.e. in a population that is monomorphic for ***z***. The quantities with a superscript ◦ may thus depend on ***z*** but we do not write such dependency to avoid notational clutter. The first two terms of eq. (10) (labelled as “intra-generational effects”) thus correspond to the standard selection gradient in group-structured population, where traits have effects within generations only. This can be read as Hamilton’s rule in gradient form, i.e. as 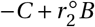, where the cost is −*C* = *∂w*_*i*_ /*∂z*_*ip*_, and the benefit is *B* = (*N* − 1)*∂w*_*i*_ /*∂z*_*jp*_ (Rousset, 2004).

Directional selection on inter-generational effects is captured by the third term of eq. (10) (labelled as “inter”). This consists of *∂w*_*i*_ /*∂ϵ*, which is the fitness effect of an environmental change, weighted by E^◦^ [*∂ϵ*/*∂z*_*p*_], which is the expected effect of a change in trait *p* in all the local ancestors (i.e. from the same patch) of a focal individual on the environment experienced by this focal. Expectation here is taken over the neutral distribution of local genealogies of the focal (i.e. the distribution of the number of local ancestors of the focal at each past generation when the population is monomorphic, hence the superscript ◦; eq. B-4 in Appendix A for definition of E^◦^[.]). What the last term of eq. (10) tells us then is that selection favours a trait change when this change in a local lineage of individuals perturbs the local environment in a way such that on average, it increases the fitness of the members of that lineage (e.g. the production of a long-lasting common good that increases the fitness of downstream relatives, Lehmann, 2007). The inter-generational nature of these effects are revealed by expanding E^◦^ [*∂ϵ*/*∂z*_*p*_] in terms of the environmental map *F* (eq. 1) as,

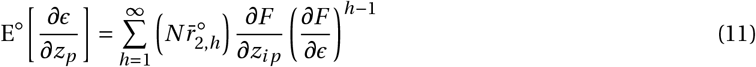

(Appendix B.3-B.4 for derivation – see also eqs. 16-20 in Mullon and Lehmann, 2018), where *∂F* /*∂z*_*ip*_ is the effect of a change in trait *p* in one focal individual (individual *i*) on its local environment over one generation (from eq. 1), and 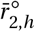 is the probability that an individual randomly sampled *h* ≥ 1 generations ago in the same patch to a focal individual is identical-by-descent to this focal (eq. B-21 in Appendix A for formal definition). Eq. (11) can be understood as follows (Fig. 1C for diagram). Consider the patch of the focal *h* generations ago. In this past generation, the focal individual has on average 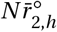 ancestors in the patch. Each of these ancestors perturbs the environment by *∂F* /*∂z*_*ip*_ (solid green arrow in Fig. 1C) due to a change in trait *p*. In turn, each perturbation persists through time via ecological inheritance, carried over each generation by a factor *∂F* /*∂ϵ*. Thus a perturbation initiated *h* generations ago has decayed by a factor (*∂F* /*∂ϵ*)^*h*−1^ by the time it reaches the focal (dashed green arrow in Fig. 1C). Summing such effects from all past ancestors (so from *h* = 1 to ∞) obtains eq. (11).

Overall, the last summand of eq. (10) may be complicated, but it captures a simple biological notion: the intergenerational effects of a trait are shaped by directional selection to benefit downstream relatives (Lehmann, 2007). In our model, these benefits are transmitted through ecological inheritance of the state of the patch (according to *∂F* /*∂ϵ*) and affect the fitness of relatives when dispersal is limited (so that inter-generational relatedness is greater than zero 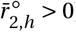).

The selection gradient ***s***(***z***) (eq. 10) characterises directional selection and thus convergence stable trait values ***z***^∗^ towards which the population evolves when traits have inter-generational ecological effects. The selection gradient, however, cannot tell us whether such traits eventually become polymorphic. This requires an understanding of disruptive selection (Dercole and Rinaldi, 2008), which we describe next.

#### 2.2.3 Disruptive, stabilising and correlational selection

Once the population expresses a convergence stable phenotype ***z***^∗^, selection is either: (i) stabilising, in which case any mutant different from ***z***^∗^ is purged so that the population remains monomorphic for ***z***^∗^; or (ii) disruptive, favoring alternative phenotypes leading to polymorphism (Metz et al., 1995; Rousset, 2004; Dercole and Rinaldi, 2008). Whether selection is stabilising or disruptive when *n* traits coevolve depends on the Hessian matrix (Leimar, 2009),

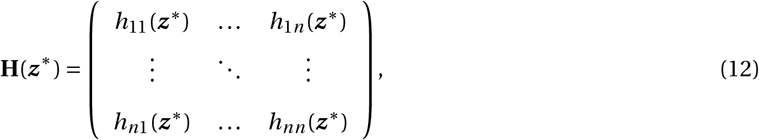

whose (*p, q*)-entry is given by

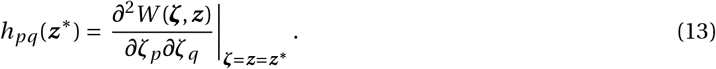

On its diagonal, **H**(***z***^∗^) indicates whether selection is disruptive on each trait (Lande and Arnold, 1983). Specifically, when *h*_*pp*_ (***z***^∗^) > 0, selection on trait *p* is disruptive when *p* evolves in isolation from the other traits (i.e. when all the other traits are fixed). Conversely, selection on trait *p* is stabilising when *h*_*pp*_ (***z***^∗^) < 0. The off-diagonal elements of **H**(***z***^∗^), meanwhile, give the strength of correlational selection, *h*_*pq*_ (***z***^∗^), on each pair of traits *p* and *q* (Lande and Arnold, 1983). These indicate the type of among-traits associations that selection favours within individuals: when *h*_*pq*_ (***z***^∗^) > 0, selection favours a positive association (or correlation) among traits *p* and *q*, and a negative association when *h*_*pq*_ (***z***^∗^) < 0. With *n* traits coevolving, selection in a population expressing a convergence stable phenotype ***z***^∗^ is stabilising when the leading eigenvalue of **H**(***z***^∗^) is negative, and disruptive when the eigenvalue is positive (Leimar, 2009; Débarre et al., 2014; Geritz et al., 2016).

The Hessian **H**(***z***^∗^) in eq. (13) is defined in terms of invasion fitness *W* (***ζ, z***). Our aim in this paper is to characterise the Hessian matrix, and thus correlational and disruptive selection, in terms of individual fitness *w*_*i*_ (eq. 4) and inter-generational effects through the environmental map *F* (eq. 1), similarly to the selection gradient shown in section 2.2.2. Our derivations can be found in Appendix C. We summarise our results in the next section.

## 3 Correlational selection under ecological inheritance

We first show in Appendix C.1 that correlational selection on traits *p* and *q* (or disruptive selection on trait *p* when *p* = *q*) can be decomposed as the sum of two terms,

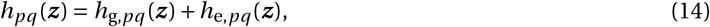

which we detail in sections 3.1 and 3.2, respectively.

### 3.1 Intra-generational fitness effects

The first term of eq. (14), *h*_g,*pq*_ (***z***), corresponds to correlational selection due to the intra-generational effects of traits on fitness (so ignoring inter-generational ecological effects on fitness and focusing on genetic effects on fitness only, hence the g in the subscript of *h*_g,*pq*_ (***z***)). We show in Appendix C.2 that it can be expressed as

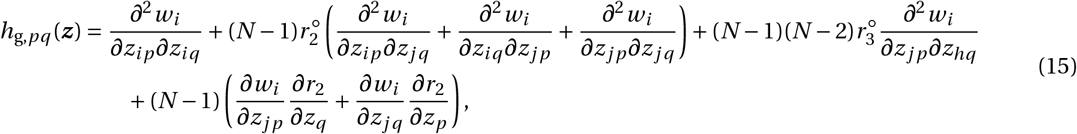

which is equivalent to the coefficient of correlational selection derived in previous papers where traits have intra-generational effects only (eqs. 13a-b-c in Mullon et al. 2016, eqs. 7a-b-c in Mullon and Lehmann 2019; see also Ajar, 2003; Wakano and Lehmann, 2014 for the case where *p* = *q*). We refer interested readers to these papers for a detailed interpretation of eq. (15), but briefly this equation can be read as the sum of three terms. The first, *∂*^2^ *w*_*i*_ /(*∂z*_*ip*_*∂z*_*iq*_), is the effect of joint changes in traits *p* and *q* of the focal individual on its own fitness. This cross derivative quantifies the synergistic (or “multiplicative” or “interaction”) effects of traits on fitness: when positive, it tells us that fitness increases more when both traits *p* and *q* change in a similar way (i.e. both increase or both decrease, so that *p* and *q* have complementary effects on fitness); conversely when negative, fitness increases more when both traits *p* and *q* change in opposite ways (i.e. one increases and the other decreases, so that *p* and *q* have antagonistic effects on fitness). In a well mixed population, correlational selection depends only on such “direct” synergy (i.e. synergistic effects of focal traits on focal fitness, so that *h*_*pq*_ (***z***) = *∂*^2^ *w*_*i*_ /(*∂z*_*ip*_*∂z*_*iq*_), Lande, 1979; Phillips and Arnold, 1989; Leimar, 2009; Débarre et al., 2014).

The rest of eq. (15) is due to limited dispersal and the interactions among contemporary relatives (i.e. living at the same generation) that result from such limitation. The remainder of the first of line of eq. (15) consists of what can be referred to as “indirect” synergistic effects (Mullon and Lehmann, 2019). These are effects of joint changes in traits *p* and *q* on focal fitness, where at least one of these changes occurs in a neighbour to the focal, weighted by relevant relatedness coefficients. Specifically, *∂*^2^ *w*_*i*_ /(*∂z*_*ip*_*∂z*_*jq*_) is the effect on focal fitness of joint changes in trait *p* in the focal and in trait *q* in a neighbour (indexed *j*), and *∂*^2^ *w*_*i*_ /(*∂z*_*jp*_*∂z*_*jq*_), of changes in traits *p* and *q* in the same neighbour *j*. Both are weighted by 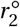. The first line of eq. (15) also features *∂*^2^ *w*_*i*_ /(*∂z*_*jp*_*∂z*_*hq*_), which is the effect on focal fitness of joint changes in different neighbours (indexed *j* and *h* with *j* ≠ *h*). This is weighted by 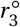, which is the coefficient of threeway relatedness, i.e. the probability that three individuals randomly sampled in a patch under neutrality are identical-by-descent. Finally, the second line of eq. (15) consists of the product between the indirect fitness effect of one trait (*∂w*_*i*_ /*∂z*_*jp*_), and the effect of the other on pairwise relatedness (*∂r*_2_/*∂z*_*q*_), which quantifies the effect of a trait change on the probability that a rare mutant individual expressing this change interacts with another mutant in the same patch (eq. C-15 for formal definition). This reveals in particular that selection favours an association among two traits when one trait improves the fitness of contemporary neighbours, and the other trait increases the probability that these contemporary neighbours are relatives (Mullon et al., 2016, 2018; Mullon and Lehmann, 2019).

### 3.2 Inter-generational effects: three pathways for correlational selection via ecological inheritance

The second term of eq. (14), *h*_e,*pq*_ (***z***), is correlational selection due to the inter-generational ecological effects of traits on fitness (hence the e of *h*_e,*pq*_ (***z***)) and thus constitutes the more novel part of our results. We find that this coefficient can be decomposed as three terms,

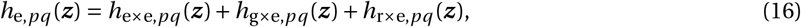

corresponding to three pathways through which correlational selection can act owing to ecological inheritance (eq. C-18 in Appendix for decomposition).

#### 3.2.1 Environmentally-mediated synergy

The first pathway,

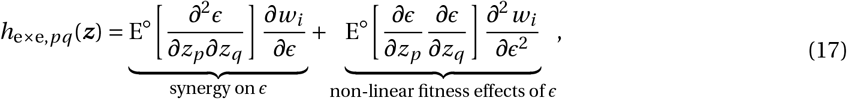

can be thought of as the synergistic effects of traits on fitness via the environment (hence the e×e subscript). Each of the two terms of eq. (17) reveals one way such synergy can come about. The first, labelled “synergy on *ϵ*”, is the most intuitive. It consists of the product between: (i) *∂w*_*i*_ /*∂ϵ*, which is the fitness effect of an environmental change; and (ii) E^◦^ [*∂*^2^*ϵ*/(*∂z*_*p*_*∂z*_*q*_)], which is the expected effect of a change in both traits *p* and *q* in all the local ancestors of a focal individual on the environment experienced by this focal (where recall from section 2.2.2 that with E^◦^[.], expectation is taken over the neutral distribution of local genealogies of the focal, eq. B-4 in Appendix A for definition of E^◦^[.]). This expectation quantifies the inter-generational environmental modifications made by a lineage of individuals that express a joint change in traits *p* and *q*. The first term of eq. (17) says that selection will associate two traits *p* and *q* when these have synergistic effects on the environment (according to E^◦^ [*∂*^2^*ϵ*/(*∂z*_*p*_*∂z*_*q*_)]) that in turn affects fitness (i.e. *∂w*_*i*_ /*∂ϵ* ≠ 0). As an example, consider a scenario where *ϵ* is the amount of a common good that can be transferred between generations (e.g. pyoverdine in siderophore-producing bacteria), so that fitness increases with such an amount (*∂w*_*i*_ /*∂ϵ* > 0). Let trait *p* be the production of this common good and *q* its protection against degradation or expropriation (so that *ϵ* depends on the product between both traits). Traits *p* and *q* would have complementary effects on *ϵ* in this example (E^◦^ [*∂*^2^*ϵ*/(*∂z*_*p*_*∂z*_*q*_)]> 0). The first term of eq. (17) tells us that in this case, selection favours a positive association between both traits, i.e. that individuals who tend to participate more in the common good also tend to protect it more.

The second term of eq. (17), labelled “non-linear fitness effects of *ϵ*”, indicates that correlational selection can also associate traits that influence the environment independently of one another, according to E^◦^[*∂ϵ*/*∂z*_*p*_ × *∂ϵ*/*∂z*_*q*_], which is the expectation of the product between the effect of a change in trait *p* in all the local ancestors of a focal individual on the environment experienced by this focal, and such an effect of a change in trait *q*. These independent ecological effect lead to correlational selection when the environment has non-linear effects on fitness (so *∂*^2^*w*_*i*_ /*∂ϵ*^2^ ≠ 0). In the common good example introduced in the previous paragraph for instance, selection for a positive association among production and protection would be strengthened where fitness accelerates with the amount of common good (*∂*^2^*w*_*i*_ /*∂ϵ*^2^ > 0) and weakened where it decelerates (*∂*^2^*w*_*i*_ /*∂ϵ*^2^ < 0). This is because such non-linearity in fitness creates synergy among traits via their ecological effects.

Correlational selection thus emerges when traits have synergistic effects on the environment, or when traits in-fluence the environment which in turn has non-linear effects on fitness. The strengths of these two effects depend on the inheritance of ecological effects from local ancestors, as quantified in eq. (17) by E^◦^ [*∂*^2^*ϵ*/(*∂z*_*p*_*∂z*_*q*_)] and E^◦^ [*∂ϵ*/*∂z*_*p*_ × *∂ϵ*/*∂z*_*q*_], respectively. We expand those in terms of the environmental *F* (eq. (1)) in Appendix C.3.1. We show in particular that,

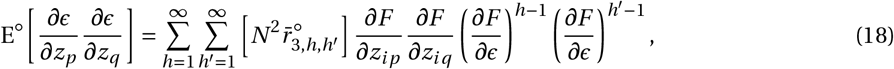

in which 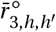 is the inter-generational threeway coefficient of relatedness: the probability that a focal individual and two randomly sampled individuals from that same patch *h* > 0 and *h*^′^ > 0 generations before the focal are all identical-by-descent under neutrality (*h* here is a dummy variable, not to be mixed up with correlational selection *h*_*pq*_ (*z*), which we always write as a function of *z* to avoid confusion; see eq. C-23 in Appendix for formal definition of 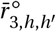). We explain eq. (18) graphically in Supplementary Fig. S1. The expression for the synergistic effects of traits on the environment, E^◦^ [*∂*^2^*ϵ*/(*∂z*_*p*_*∂z*_*q*_)], is more complicated and more difficult to parse and we have therefore left it in Appendix C.3.1 (eq. C-34).

#### 3.2.2 Genes-environment interactions

The second pathway through which correlational selection can act owing to ecological inheritance (second term of eq. 16) is given by

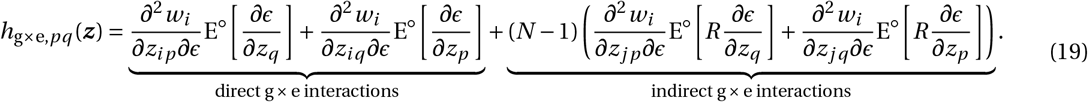

This pathway emerges when fitness is influenced by the interaction between the environment and traits (or the genes coding for these traits, hence the g×e subscript). Specifically, the first term of eq. (19), labelled “direct g × e interactions”, consists of the interaction between the environment and the expression of one trait by the focal (*∂*^2^ *w*_*i*_ /(*∂z*_*ip*_*∂ϵ*)), multiplied to the inter-generational ecological effects of the other trait (E^◦^ *∂ϵ*/*∂z*_*q*_, which is the expected effect of a change in trait *q* in all the local ancestors of a focal individual on the environment experienced by this focal and is given by eq. 11). To understand the implications of this, consider again a scenario where *ϵ* is some inter-generational common good and one trait, say *q*, is the production or maintenance of this common good (so that E^◦^ [*∂ϵ*/*∂z*_*q*_]> 0). The other trait *p*, however, now is some costly competitive trait whose cost depends on environmental conditions (e.g. horn length in beetles; Emlen, 1994), so that individuals living in better patches (i.e. with greater *ϵ*) pay a lower expression cost (leading to *∂*^2^*w*_*i*_ /(*∂z*_*ip*_*∂ϵ*)) > 0). The first term of eq. (19) in this example would be positive, indicating that it favours a positive association between traits *p* and *q*, i.e. individuals who participate more to the common good also express larger competitive traits. This is because individuals who contribute more to the common good also tend to live in better habitats (owing to limited dispersal and past relatives contributing to the environment, see eq. 11). They can thus also afford to express larger competitive traits.

The second term of eq. (19), labelled “indirect g × e interactions”, consists of the interaction between the environment and the expression of one trait by a neighbour of the focal (*∂*^2^*w*_*i*_ /(*∂z*_*jp*_*∂ϵ*)), multiplied to E^◦^[*R* × *∂ϵ*/*∂z*_*q*_], which is the expected product between: (i) the ecological effects of the other trait (*∂ϵ*/*∂z*_*q*_); and (ii) the frequency of relatives among the neighbours of the focal, *R*, which here should be seen as a random variable (with expectation E^◦^[.] taken as before over the distribution of local genealogies under neutrality so that 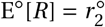). This expected product E^◦^[*R* × *∂ϵ*/*∂z*_*q*_] is indicative of the covariance between the genetic and ecological environments of the focal. When E^◦^[*R* × *∂ϵ*/*∂z*_*q*_] is large, environmental transformations driven by trait *q* tend to have a large effect not only the focal individual but also on its contemporary patch relatives. If in turn, trait *p* of these relatives interacts with the environment to increase the fitness of the focal individual (according to *∂*^2^*w*_*i*_ /(*∂z*_*jp*_*∂ϵ*)), eq. (19) reveals that correlational selection will associate these two traits. To see what indirect gene-environment interactions might entail, consider a situation where *ϵ* is the state of the patch, trait *q* is some investment to maintain this state for future generations, and trait *p* is a trait that increases the fitness of neighbours living in the current generation such as helping. Assume further that the benefits of helping decrease with the patch state *ϵ*, for instance because helping is mostly relevant when the environment is of low quality. Mathematically, this translates into negative indirect gene-environment interactions: *∂*^2^*w*_*i*_ /(*∂z*_*jp*_*∂ϵ*) < 0. According to eq. (19), this favours a negative correlation between investing into current members through helping (via *p*) and investing into future patch members through patch maintenance (via *q*), i.e. individuals that invest more into future relatives invest less in present relatives and vice-versa.

The relevance of indirect gene-environment interactions for correlational selection depends on E^◦^[*R* ×*∂ϵ*/*∂z*_*q*_], which we show is in terms of the environmental map *F* (eq. 1) given by

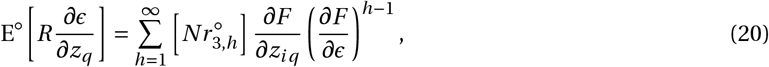

where 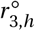 is the probability that two individuals living in the same generation, plus a third individual *h* generations ago, all randomly sampled from the same patch are identical-by-descent (eq. C-39 for definition; Appendix C.3.2 for derivation eq. 20). This probability indicates the likelihood for an individual to influence the environment of at least two downstream relatives living *h* generations away in the same patch and thus directly interacting with one another. Accordingly, the greater 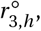, the more influence a modification to the patch environment can have on social interactions, and thus the more relevant indirect gene-environment effects are to selection (Supplementary Fig. S2 for a graphical interpretation of eq. 20).

#### 3.2.3 Biased ecological inheritance

The third and final pathway for correlational selection (*h*_r×e,*pq*_ (***z***) in eq. 16) can be expressed as

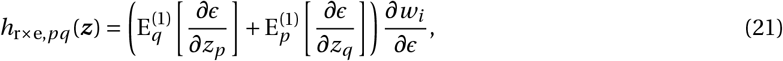

where 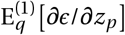 is the expected effect of a change in trait *p* in all the local ancestors of a focal individual on the environment experienced by this focal, where expectation is taken over the perturbation of the distribution of local genealogies owing to a change in trait *q* (hence the superscript (1) and subscript *q* in 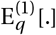 to contrast with E^◦^[.], which is expectation over the neutral distribution; eq. C-6 for a formal definition of 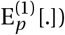. More intuitively perhaps, 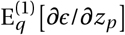 quantifies how trait *q* influences the way an environmental modification driven by a change in trait *p* is inherited. This can be seen more explicitly when we unroll 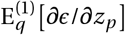 over its inter-generational effects:

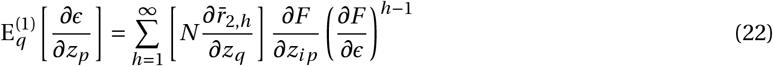

(Appendix C.3.3 for derivation). Here, 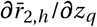 is the effect of a change in trait *q* on the relatedness between individuals living in the same patch separated by *h* generations. To understand this effect better, consider a focal individual who expresses a change in trait *q* relative to a resident. This 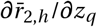 quantifies how such a trait change influences the probability that an individual randomly sampled in that same patch *h* generations ago is identical-by-descent to the focal, relative to the probability that two resident individuals separated by *h* generations are identical-by-descent under neutrality (i.e. relative to 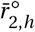, Appendix. E.2 for details). So when for instance 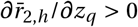, individuals that express greater values of trait *q* are more likely to transmit environmental modifications to their kin. What eq. (22) substituted into (21) in turn says is that correlational selection will associate this trait *q* with another trait *p* when trait *p* leads to inter-generational environmental modifications that increase fitness (i.e. so that *∂F* /*∂z*_*ip*_ > 0 in eq. 22 and *∂w*_*i*_ /*∂ϵ* > 0 in eq. 21; Supplementary Fig. S3 for diagram). We explore the potential implications of such correlational selection in section 4 with a specific example.

### 3.3 Summary

To summarize our findings so far, we have identified three main pathways via which traits can be linked by correlational selection under ecological inheritance (eq. 16). Each of these pathways can be expressed in terms of how a trait change in an individual causes an environmental modification that in turn influences the fitness of future relatives (eq. 11, eq. 18, eq. 20, eq. 22, and eq. C-34 in Appendix C.3.1). This perspective not only offers a clear view on correlational selection on inter-generational effects, it also allows us to efficiently compute the Hessian matrix and thus investigate the conditions that lead to polymorphism in traits that influence the environment in the long term. In fact, what remains to be characterized for such computation are the various relevant relatedness coefficients (e.g. 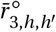) or their perturbation due to selection (e.g. 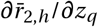). We do so in Appendices D and E. Substituting for these into eq. (11), eq. (18), eq. (20), eq. (22), and eq. (C-34) in Appendix C.3.1, we obtain the expressions found Table 1. This Table 1, together with eq. (10), eq. (14), eq. (15), eq. (16), eq. (17), eq. (19), and eq. (21), gives all that is necessary to investigate directional, correlational and disruptive selection under the general model described in section 2.1. We illustrate such an approach in the next section.

**Table 1:** Weights on fitness effects relevant to directional, disruptive and correlational selection under eco-logical inheritance. Recall that *N* is the (fixed) number of individuals per patch, and *m* is backward dispersal, i.e. the probability that an individual randomly sampled in a patch under neutrality is an immigrant. This probability may be a fixed parameter or an evolving variable, in which case *m* depends on the resident trait ***z*** (as e.g. when traits that influence gene flow evolve, like in our example section 4). The quantity *w*_p,*i*_ is the philopatric component of individual fitness, i.e. the expected number of offspring of individual *i* that remain in their natal patch, which depends on the same parameter as total individual fitness eq. (4) (eq. 33 for an example). ^†^ Relevant equation in Appendix where derivation can be found.

| Symbol | Value | Eq.<sup>†</sup> |
| --- | --- | --- |
| $r_2^\circ$ | $= \frac{(1-m)^2}{N - (N-1)(1-m)^2}$ | (D-2) |
| $\bar{r}_2^\circ$ | $= \frac{1}{N} + \frac{N-1}{N} r_2^\circ$ | (D-3) |
| $r_3^\circ$ | $= \frac{(1-m)^3 (1 + 3(N-1)r_2^\circ)}{N^2 - (N-1)(N-2)(1-m)^3}$ | (D-5) |
| $\bar{r}_3^\circ$ | $= \frac{1}{N^2} + \frac{3(N-1)}{N^2} r_2^\circ + \frac{(N-1)(N-2)}{N^2} r_3^\circ$ | (D-6) |
| $E^\circ \left[ \frac{\partial \epsilon}{\partial z_p} \right]$ | $= \frac{\partial F}{\partial z_{ip}} \frac{N(1-m)}{1 - \frac{\partial F}{\partial \epsilon}(1-m)} \bar{r}_2^\circ$ | (B-25) |
| $E^\circ \left[ \frac{\partial \epsilon}{\partial z_p} \frac{\partial \epsilon}{\partial z_q} \right]$ | $= \frac{\partial F}{\partial z_{ip}} \frac{\partial F}{\partial z_{iq}} \frac{N^2(1-m)}{1 - \left(\frac{\partial F}{\partial \epsilon}\right)^2 (1-m)} \left[ \bar{r}_3^\circ + 2 \frac{\partial F}{\partial \epsilon} C_\epsilon \right]$ | (C-33) |
| $E^\circ \left[ \frac{\partial^2 \epsilon}{\partial z_p \partial z_q} \right]$ | $= \frac{N(1-m)}{1 - \frac{\partial F}{\partial \epsilon}(1-m)} \left( \frac{\partial^2 F}{\partial z_{ip} \partial z_{iq}} \bar{r}_2^\circ + (N-1) \frac{\partial^2 F}{\partial z_{ip} \partial z_{jq}} \left[ \frac{2}{N} r_2^\circ + \frac{N-2}{N} r_3^\circ \right] \right)$<br>$+ \frac{\partial F}{\partial z_{ip}} \frac{\partial F}{\partial z_{iq}} \frac{\partial^2 F}{\partial \epsilon^2} \frac{N^2(1-m)^2}{\left(1 - \frac{\partial F}{\partial \epsilon}(1-m)\right) \left(1 - \left(\frac{\partial F}{\partial \epsilon}\right)^2 (1-m)\right)} \left[ \bar{r}_3^\circ + 2 \frac{\partial F}{\partial \epsilon} C_\epsilon \right]$<br>$+ \frac{N^2(1-m)}{1 - \frac{\partial F}{\partial \epsilon}(1-m)} \left( \frac{\partial F}{\partial z_{ip}} \frac{\partial^2 F}{\partial z_{iq} \partial \epsilon} + \frac{\partial F}{\partial z_{iq}} \frac{\partial^2 F}{\partial z_{ip} \partial \epsilon} \right) C_\epsilon$ | (C-35) |
| $E^\circ \left[ R \frac{\partial \epsilon}{\partial z_p} \right]$ | $= \frac{\partial F}{\partial z_{ip}} \frac{N(1-m)^2}{N - \frac{\partial F}{\partial \epsilon}(1-m)^2(N-1)} \left( \frac{\frac{\partial F}{\partial \epsilon}(1-m)}{1 - \frac{\partial F}{\partial \epsilon}(1-m)} \bar{r}_2^\circ + N \bar{r}_3^\circ \right)$ | (C-42) |
| $\frac{\partial r_2}{\partial z_p}$ | $= \frac{2r_2^\circ}{1-m} \left[ \frac{\partial w_{p,i}}{\partial z_{ip}} [1 + (N-1)r_2^\circ] \right.$<br>$\left. + (N-1) \frac{\partial w_{p,i}}{\partial z_{jp}} [2r_2^\circ + (N-2)r_3^\circ] + N^2 \frac{\partial w_{p,i}}{\partial \epsilon} \frac{\partial F}{\partial z_{ip}} C_\epsilon \right]$ | (E-22) |
| $E_p^{(1)} \left[ \frac{\partial \epsilon_t}{\partial z_q} \right]$ | $= \frac{\partial r_2}{\partial z_p} \frac{\partial F}{\partial z_{iq}} \frac{N(1-m)}{1 - \frac{\partial F}{\partial \epsilon}(1-m)} \left( \frac{N-1}{N} + \frac{1}{2N r_2^\circ} \right)$<br>$+ \frac{\frac{\partial F}{\partial \epsilon}}{1 - \frac{\partial F}{\partial \epsilon}(1-m)} \left( \frac{\partial w_{p,i}}{\partial z_{ip}} E^\circ \left[ \frac{\partial \epsilon}{\partial z_q} \right] + (N-1) \frac{\partial w_{p,i}}{\partial z_{jp}} E^\circ \left[ R \frac{\partial \epsilon}{\partial z_q} \right] \right)$<br>$+ \frac{\partial w_{p,i}}{\partial \epsilon} \frac{\partial F}{\partial z_{ip}} \frac{\partial F}{\partial z_{iq}} \frac{\frac{\partial F}{\partial \epsilon} N^2(1-m)}{\left(1 - \frac{\partial F}{\partial \epsilon}(1-m)\right) \left(1 - \left(\frac{\partial F}{\partial \epsilon}\right)^2 (1-m)\right)} \left[ \bar{r}_3^\circ + 2 \frac{\partial F}{\partial \epsilon} C_\epsilon \right]$ | (E-48) |
| $C_\epsilon$ | $= \frac{(1-m)}{N - \frac{\partial F}{\partial \epsilon}(1-m)^2(N-1)} \left( (N-1)(1-m) \bar{r}_3^\circ + \frac{\bar{r}_2^\circ}{1 - \frac{\partial F}{\partial \epsilon}(1-m)} \right)$ | |

## 4 Joint evolution of dispersal with the attack rate on a local renewable resource

We now go over a specific model that looks at the joint evolution of two ecologically-relevant traits: (i) the rate of attack or consumption of a resource within patches; and (ii) dispersal between patches. The evolution of both traits have been studied in isolation in multiple studies (for dispersal: Hamilton and May (1977); Taylor and Frank (1996); Gandon and Michalakis (1999); Gandon and Rousset (1999); Ajar (2003); for resource exploitation: Van Baalen and Sabelis (1995); Pels et al. (2002); Rauch et al. (2002); Lehmann (2008); Kylafis and Loreau (2008); Messinger and Ostling (2013)) but none of those let both traits coevolve. Further, studies so far have focused on the effects of directional selection, which is not sufficient to determine whether polymorphism emerges. Here we show that it readily does.

### 4.1 A resource-consumer model in a patch-structured population

We first specify a resource-consumer scenario and lay the building blocks of our analysis.

#### 4.1.1 Traits

Each individual is characterised by two traits: (i) the rate *z*_1_ of attack on a local resource in a patch (during step (i) of the life-cycle; see section 2.1); and (ii) the probability *z*_2_ of juvenile dispersal, which we assume is costly with a probability *c*_d_ of dying during dispersal (step (iii) of the life-cycle; see section 2.1).

#### 4.1.2 Environment and its dynamics

The environmental state *ϵ* of a patch at a generation *t* is the abundance of the resource in that patch before consumption at the generation (so before step (i) of the life-cycle). We derive the inter-generational dynamics of resource abundance (i.e. of *ϵ* from *t* to *t* + 1) from a model of consumer-resource dynamics that occur in continuous time within generations (Schmid et al., 2022). To specify these dynamics, let us denote intra-generational time by *τ* which runs from 0 to *T*, where *T* is the length in continuous time of a generation (i.e. a whole iteration of the life-cycle). We let *ϵ*_*t, τ*_ be the resource abundance at time *τ* within generation *t* in a focal patch. With this notation, the abundance *ϵ*_*t*, 0_ of the resource in that patch before consumption at generation *t* is given by *ϵ* = *ϵ*_*t*, 0_ = *ϵ*_*t* −1,*T*_. To obtain *ϵ*_*t* +1_, we track *ϵ*_*t, τ*_ from *τ* = 0 to *T*. We assume that resource dynamics during that time period are decomposed into two phases: a consumption phase (for 0 ≤ *τ* ≤ *τ*_1_) followed by a renewal phase (for *τ*_1_ < *τ* ≤ *T*) so that *τ*_1_ is the amount of time the resource is being consumed (and *T* − *τ*_1_ the amount of time it renews itself).

##### Consumption phase

First, each individual *i* ∈ {1,…, *N* } in the patch consumes the resource at a rate give by its trait *z*_*i* 1_. Specifically, the rate of change in the amount *ρ*_*i, τ*_ of resources collected by individual *i* = 1,…, *N* at time *τ* is

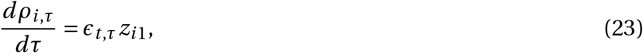

while the rate of change in resource abundance in the patch is

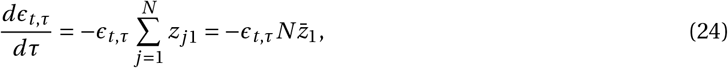

where 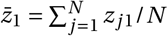 is the average attack rate in the patch. Solving eqs. (23) and (24) with initial conditions *ρ*_*i*, 0_ = 0 for each *i* ∈ {1,…, *N* }, we obtain that by the end of consumption, the resource abundance in the patch is

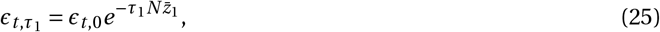

and that the amount of resources consumed by a focal individual *i* is,

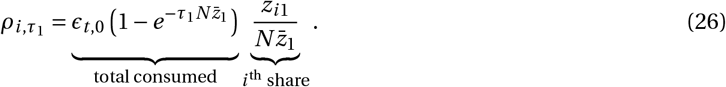

As highlighted with the underbraces in eq. (26), this amount can be separated between the total amount of resources consumed in the patch, and the share obtained by individual *i*. Eq. (26) can thus be seen as a contest success function of the ratio type, which is commonly used to model competition for resources (Hirshleifer, 1989).

##### Renewal phase

After consumption, we assume the resource renews itself growing logistically, according to

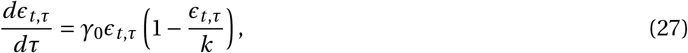

where *γ*_0_ is the per capita growth rate of the resource when at low abundance, and *k* the carrying capacity of a patch for the resource, which we set to *k* = 1 for simplicity. Solving eq. (27) for *τ*_1_ < *τ* ≤ *T* with initial condition given by eq. (25), we obtain that by the end of generation *t* (so at the beginning of generation of *t* + 1), the resource abundance is

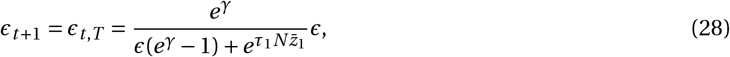

where *γ* = *γ*_0_(*T* − *τ*_1_) is per-generation renewal rate (letting *γ* → ∞ then recovers the model of Schmid et al., 2022, in which resource abundance is fixed between generations). Using the notation of eq. (1), the environmental map of our model then is,

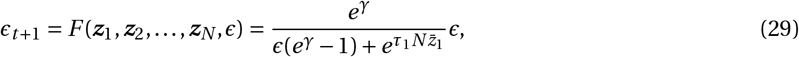

where *ϵ* = *ϵ*_*t*, 0_ is the resource abundance at the beginning of generation *t*. The environmental map eq. (29) is independent of dispersal (trait *z*_2_) as dispersal does not influence the way individuals consume resources.

##### Resource equilibrium

Solving for equilibrium 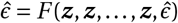 (eq. 2) and checking its stability (using eq. 3), we find that in a monomorphic population where each individual express the same ***z*** = (*z*_1_, *z*_2_), the resource abundance stabilises for

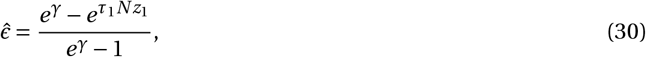

which is positive as long as the renewal rate *γ* is large enough compared to consumption (specifically when *γ* > *Nz*_1_*τ*_1_). Otherwise, the resource goes extinct. We focus our attention on the case where the resource is maintained (so where *γ* > *Nz*_1_*τ*_1_).

#### 4.1.3 Fitness

An individual uses the resources it has collected to produce offspring, favoring increased attack rate. Increasing one’s attack rate may however also be costly, for instance due to lost opportunities or increased risk. To reflect these benefits and costs, we assume that the fecundity of a focal individual *i* with attack rate *z*_*i* 1_ in a patch where its neighbours express rates 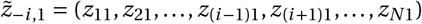 and the resource abundance is *ϵ*, is given by

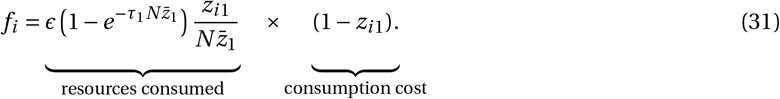

This equation consists of the product between the amount of resources consumed by the focal individual (eq. 26 with *ϵ*_*t*, 0_ = *ϵ*) and the individual cost of consumption. From eq. (31), fecundity in a population monomorphic for ***z*** is

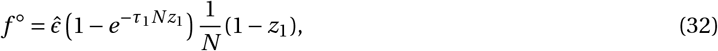

where the equilibrium amount of resources 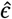 is given by eq. (30). Under the island model of dispersal (e.g. eq. 6.5 in Rousset, 2004), the fitness of a focal individual can then be written as

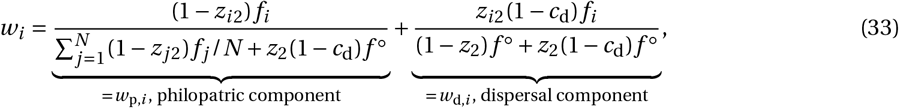

where the first term, *w*_p,*i*_, is the philopatric component of fitness, i.e. the expected number of offspring that establish in their natal patch (consisting of the ratio of offspring of the focal that remain in their natal patch to the total number of offspring that enter competition in that patch); and the second, *w*_d,*i*_, is the dispersal component, i.e. the expected number of offspring that establish in non-natal patches (consisting of the ratio of offspring of the focal that leave their natal patch to the total number of offspring that enter competition in another patch).

#### 4.1.4 Relatedness

In addition to the environmental map (eq. 29) and fitness (eq. 33), the analysis described in section 3 relies on several relatedness coefficients under neutrality and selection (Table 1). Those in turn depend on the backward probability of dispersal, *m* (i.e. the probability that in a monomorphic population, a randomly sampled individual is an immigrant). In this model where dispersal *z*_2_ is evolving, such probability is given by

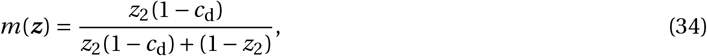

which consists of the ratio of the number of individuals dispersing into a patch to the total number of individuals that compete for breeding spots.

Equations (29)-(34) (together with Table 1) are all the ingredients necessary to perform the analysis of disruptive selection laid out in section 3 (and section 2 for directional selection). Details on our analysis can be found in Appendix F whose main results we summarize below, paying special attention on whether polymorphism emerges. All our derivations can also be followed from the accompanying Mathematica Notebook (Supplementary Files).

### 4.2 Directional selection: convergence to intermediate dispersal and attack traits

Substituting eqs. (29)-(33) into the selection gradient eq. (10) and analysing this gradient according to the approach described in section 2.2.1, we find that the population first converges to a singular strategy 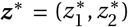 for both traits. In line with previous results (e.g., Hamilton and May, 1977; Taylor, 1988; Ajar, 2003), the singular value 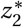 for dispersal reads as,

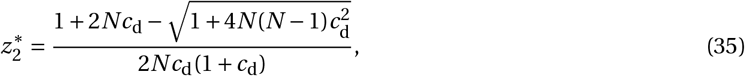

which decreases with the cost *c*_d_ of dispersal and with the number *N* of individuals per patch (Fig. 2Ai and Supplementary Fig. S4Ai, Appendix F.2.2 for details). The dispersal singular value 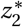 is independent from the attack rate. This is because the attack rate does not influence how fitness varies due to a marginal change in dispersal (as *f*_*i*_ = *f* ^◦^ for all *i* in eq. 33 when the population is monomorphic for *z*_1_).

**Figure 2:**
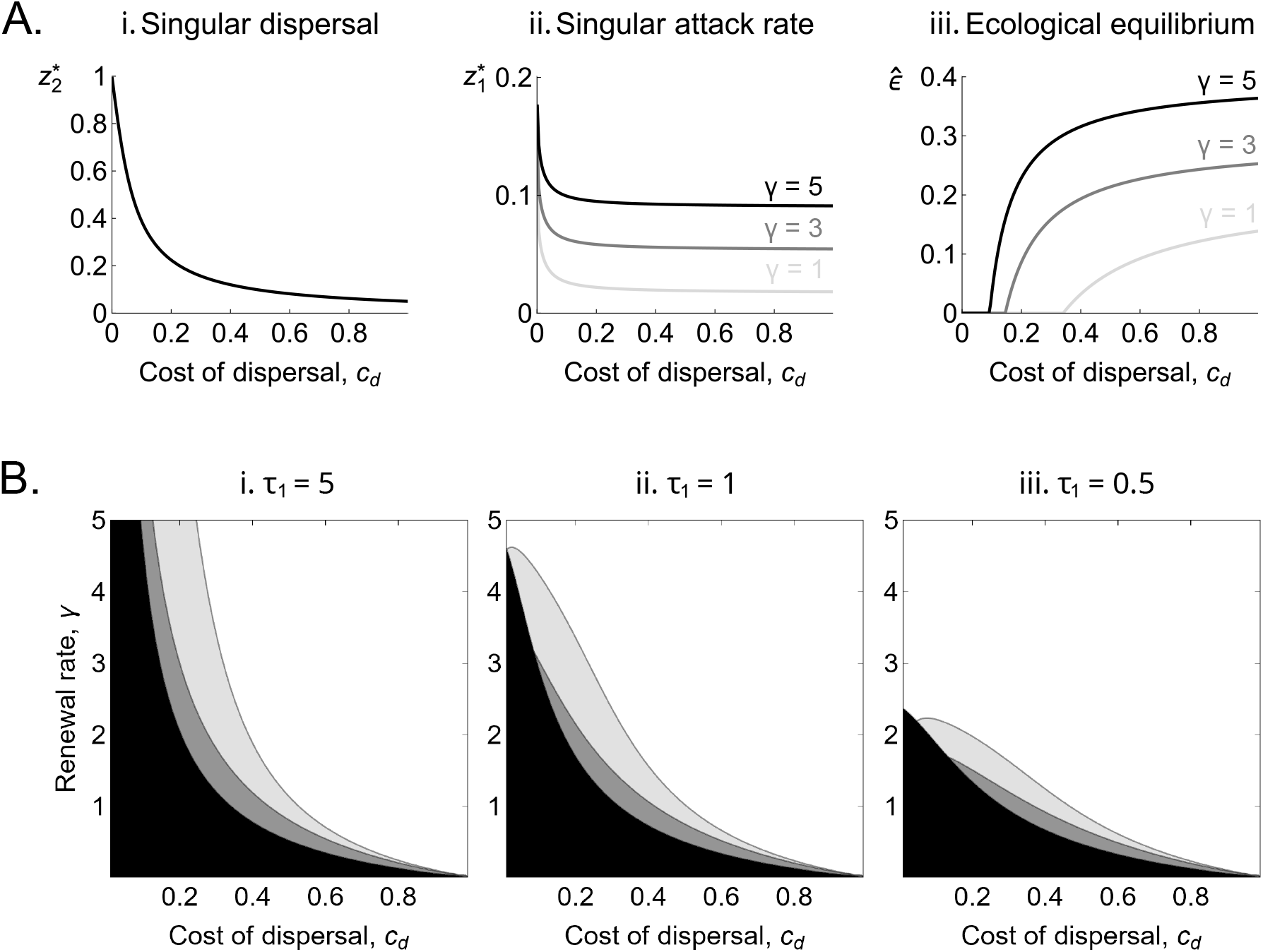
A. Convergence stable (i) dispersal and (ii) attack rate, and (iii) associated resource abundance. Obtained from: (i) eq. (35); (ii) solving eq. (36) numerically for 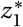; and (iii) from eq. (30) with 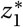 (black: *γ* = 5; dark gray: *γ* = 3; light gray: *γ* = 1). Note from Aiii that not all convergence stable attack rates plotted in Aii lead to the maintenance of the resource (i.e. lead to 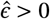). When renewal rate is low for instance (*γ* = 1), the cost of dispersal *c*_d_ must be greater than 0.35 for resource maintenance. Other parameters: *τ*_1_ = 5, *N* = 10. See Supplementary Fig. S4A for the effect of *N*. **B. Ecological and evolutionary outcomes**. These graphs show the parameter regions where the joint evolution of attack rate and dispersal leads to: resource extinction (in black); disruptive selection (in gray; dark gray for when the attack rate *z*_1_ evolves alone and dispersal is fixed for 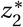); and stabilising selection (in white). In (i) *τ*_1_ = 0.5; (ii) *τ*_1_ = 1; (iii) *τ*_1_ = 5; other parameters: *N* = 10, Supplementary Fig. S4B for the effects of *N*. Appendix F and Mathematica Notebook (Supplementary Files) for analysis.

The singular value 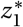 for the attack rate, meanwhile, satisfies the following equality,

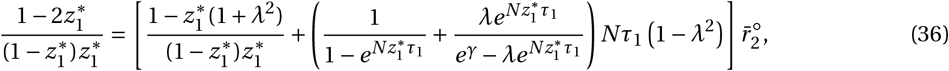

where *λ* = 1 − *m*(***z***^∗^) is the probability that an individual is philopatric in a population monomorphic for ***z***^∗^, which thus depends on the evolved dispersal strategy 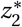 (found by substituting eq. 35 into eq. 34). Solving eq. (36) for 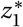 numerically, we find that when dispersal is costly (i.e. *c*_d_ is large) the population evolves lower attack rate (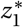 is small, Fig. 2Aii). This is because when *c*_d_ is large, dispersal evolves to be limited (eq. 35). As a result, inter-generational relatedness becomes large (i.e. 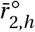 in eq. (11) becomes large), which in turn favors the evolution of restraint so that individuals leave more resources to their downstream relatives (Lehmann, 2008). Conversely, when dispersal cost *c*_d_ is low, dispersal evolves to be large, which causes inter-generational relatedness to drop, and thus the evolution of high attack rates as consumption no longer affects relatives. In contrast, the attack rate decreases with patch size *N* (Supplementary Fig. S4Aii). This is due to competition for resources being within patches such that the amount of resources collected by one individual is inversely proportional to *N* (eq. 26). Consequently, the benefit from increased attack rate is smaller when patches are larger, favouring the evolution of lower attack rates. Another relevant parameter to the evolution of the attack rate is the rate *γ* of resource renewal. In particular, high renewal rate *γ* leads to more exploitative strategies (i.e. greater 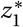, Fig. 2.A.ii). This is because when *γ* is large, ecological inheritance is weak as the resource renews itself quickly after consumption. Individuals can thus consume more resources while incurring little cost to their descendants and the descendants of their relatives (i.e. *∂F* /*∂ϵ* is small in eq. 11).

Plugging the singular strategy for attack rate 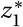 (given by eqs. 35-36) into the equilibrium resource abundance (eq. 30) allows us to understand the effect of evolutionary dynamics on the resource. As expected from the previous paragraph, high dispersal cost *c*_d_ leads to higher resource abundance as consumption evolves to be more restrained (Fig. 2.A.iii). Conversely, low dispersal cost reduces resource abundance at evolutionary equilibrium. If in addition to low dispersal cost, the rate *γ* of renewal is also low, then the resource may in fact go extinct as it is unable to renew itself fast enough in the face of increased consumption (Fig. 2.A.iii). We also find that the resource abundance at equilibrium decreases with patch size *N* (Supplementary Fig. S4Aiii).

### 4.3 Disruptive and correlational selection: the emergence of dispersive overconsumers and sessile scrimpers

To determine whether the population becomes polymorphic once it has converged to the singular phenotype 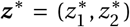 (given by eqs. 35-36), we substitute eqs. (29)-(33) into eqs. (14)-(21) and perform the analysis described in section 2.2.3 (Appendix F for details). As previously found (Ajar, 2003), selection on dispersal is always stabilising when dispersal evolves alone (eq. F-12 here for our derivation). When the attack rate evolves but dispersal is fixed at its singular strategy, selection on the attack rate is typically stabilising, except where dispersal cost *c*_d_ and renewal rate *γ* are low, close to their threshold values for resource maintenance (dark gray region in Fig 2B, Appendix F.3.2 for details). When both traits coevolve, the range of values for *c*_d_ and *γ* for which selection is disruptive is wider (dark and light gray regions in Fig. 2B). When *c*_d_ and *γ* are high, selection is always stabilising (white region in Fig. 2B). Unlike *c*_d_ and *γ*, patch size *N* has little effect on whether selection is stabilising or disruptive (Supplementary Fig. S4B).

In sum, the dynamics in our model lead to three outcomes depending mostly on the dispersal cost *c*_d_ and resource renewal rate *γ*: (i) when *c*_d_ and *γ* are both low, consumer evolution leads to resource extinction (so that if the consumer relies entirely on this resource, it would also go extinct); (ii) when *c*_d_ and *γ* are high, the consumer remains monomorphic for dispersal and attack rate such that the resource is maintained; (iii) when *c*_d_ and *γ* are intermediate, the consumer becomes polymorphic.

A closer look at correlational selection on dispersal and attack rate, *h*_12_(***z***^∗^), reveals two things about the nature of the polymorphism. The first is that since *h*_12_(***z***^∗^) is always positive (Appendix F.2.3), the polymorphism should be characterised by a positive association between the two traits. We thus expect two types to emerge: (i) one that consumes and disperses more (“dispersive overconsumers”); and (ii) another that consumes and disperses less (“sessile scrimpers”). The second relevant aspect of this polymorphism that our analysis shows is that the term that mainly contributes to correlational selection is the one capturing biased ecological inheritance, *h*_r×e,12_(***z***^∗^) (eq. 21-22, Appendix F.2.3 for details). More specifically, it is the combination of negative effects of dispersal on inter-generational relatedness, 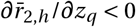, and of consumption on the environment, *∂F* /*∂z*_*i* 1_ < 0, that leads correlational selection to be positive (owing to eq. 22). This indicates that polymorphism in our model is due to a positive association between dispersal and attack rate, leading scrimpers to preferentially inherit the patch they maintain from relatives, and overconsumers to preferentially inherit the patch they deplete from non-relatives.

### 4.4 The rise and fall of overconsumption

To check our mathematical analyses, we ran individual-based stochastic simulations under conditions that lead to stabilising and disruptive selection (Appendix F.4 for simulation procedure and Supplementary Files for code). As predicted, the population gradually converges to the singular strategy for dispersal and consumption in both cases (Fig. 3AB). Concomitantly, the resource abundance goes to its equilibrium 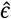 given by eq. (30) (Fig. 3CD). Where selection is stabilising, the population remains monomorphic for both traits (i.e. unimodally distributed around this strategy, Fig. 3A), and the resource abundance within patches remains distributed around the ecological equilibrium 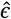 (Fig. 3C). In contrast, two morphs that correspond to dispersive overconsumers and sessile scrimpers emerge and become increasingly differentiated where selection is disruptive (Fig. 3B). In this case, the distribution of resource densities becomes bimodal so that the population consists of patches of either low (i.e. with small *ϵ*) or high quality (i.e. with larger *ϵ*, Fig. 3D). This is due to variation in morph composition among patches such that patches with a greater frequency of overconsumers are typically of low quality (Supplementary Fig. S5).

**Figure 3:**
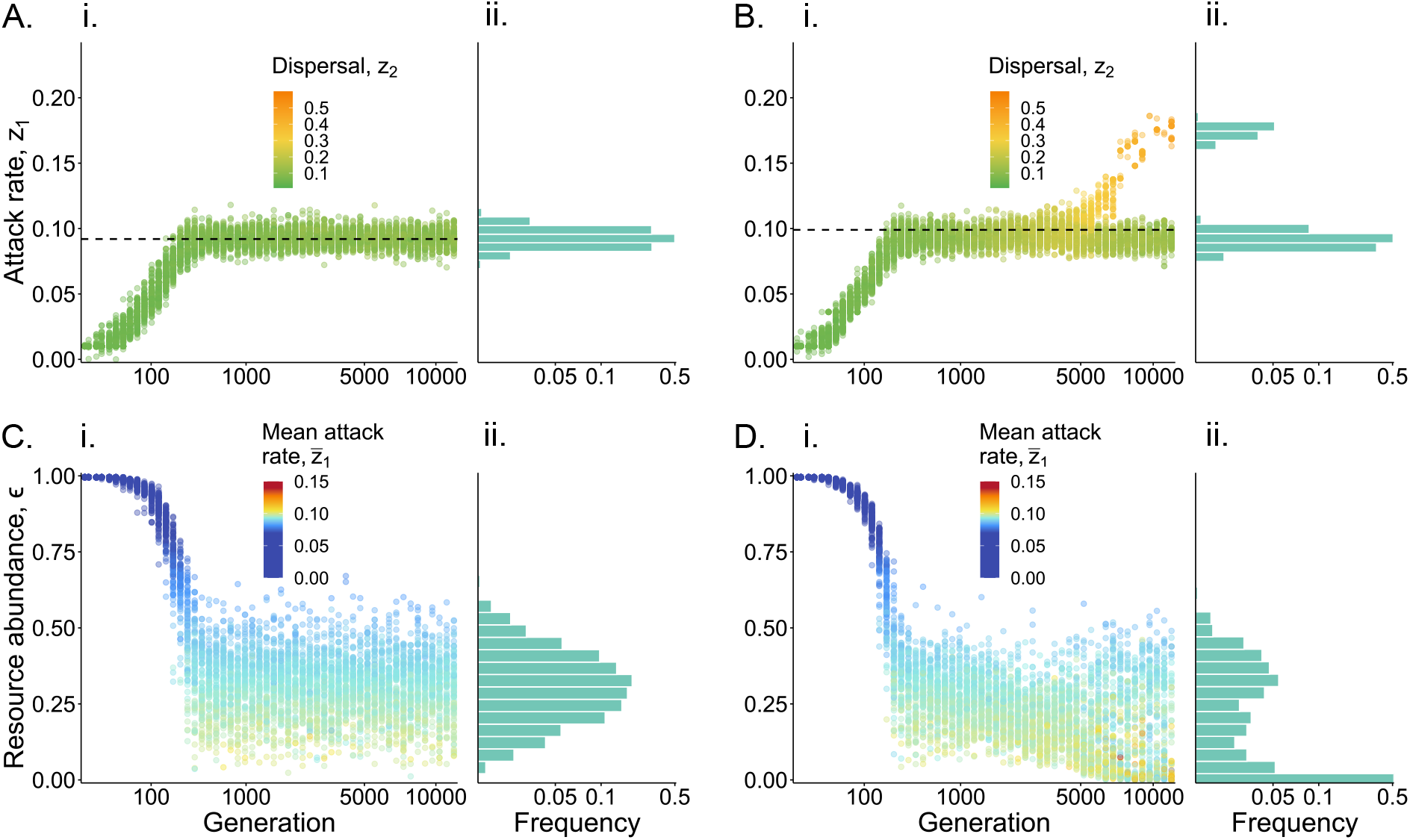
Co-evolutionary dynamics of dispersal and attack rate, with concomitant resource abundance distribution. These graph show the results of individual-based stochastic simulations for a population subdivided among *N*_p_ = 1000 patches (Appendix F.4 for details), under stabilising (A and C: *c*_d_ = 0.5) and disruptive selection (B and D: *c*_d_ = 0.1). Other parameters: *γ* = 5, *τ*_1_ = 5, *N* = 10. We chose *N* = 10 for computational expediency. Similar dynamics are observed with larger patch sizes (Supplementary Fig. S4C). **A-B: Attack rate and dispersal evolution**. In (i) each point represents an individual, coloured according to its dispersal probability (see figure for legends). (ii) histogram of equilibrium of attack rates in the population. Dashed black line shows the convergence stable strategy (from eq. 36). As expected, the population remains monomorphic under stabilising selection in A, and becomes polymorphic under disruptive selection in B, where the polymorphism is characterised by a positive association between attack and dispersal. **C-D: Resource abundance distribution and association with attack rate**. In (i) each point represents the resource abundance in that patch, coloured according to the mean attack rate in the patch (see figure for legends). Patches where individuals on average express higher attack rates (warmer colours) tend to carry fewer resources. (ii) shows histogram of resource abundance across patches.

When two morphs coexist, our simulations reveal ecological and evolutionary cycles whereby the population alternates between generations during which scrimpers are common and resources are plentiful, and generations during which overconsumers are more abundant and resources are scarce (Fig. 4A). With evolution favouring increasingly differentiated morphs, overconsumers have an increasingly detrimental effect on their patch so that the amplitude of these cycles increases (compare Fig. 4A with B; and red with blue in Fig. 4C). In fact, there comes a time when overconsumers are so rare in periods of low abundance that they may stochastically go extinct (i.e. they become so rare in our simulations that by chance, none reproduce). The population is then monomorphic for the scrimper morph, which in the absence of the overconsumer morph is counter-selected. The population thus converges once again to the singular strategy (given by eqs. 35-36), whereupon polymorphism emerges and collapses again and again (Fig. 4D).

**Figure 4:**
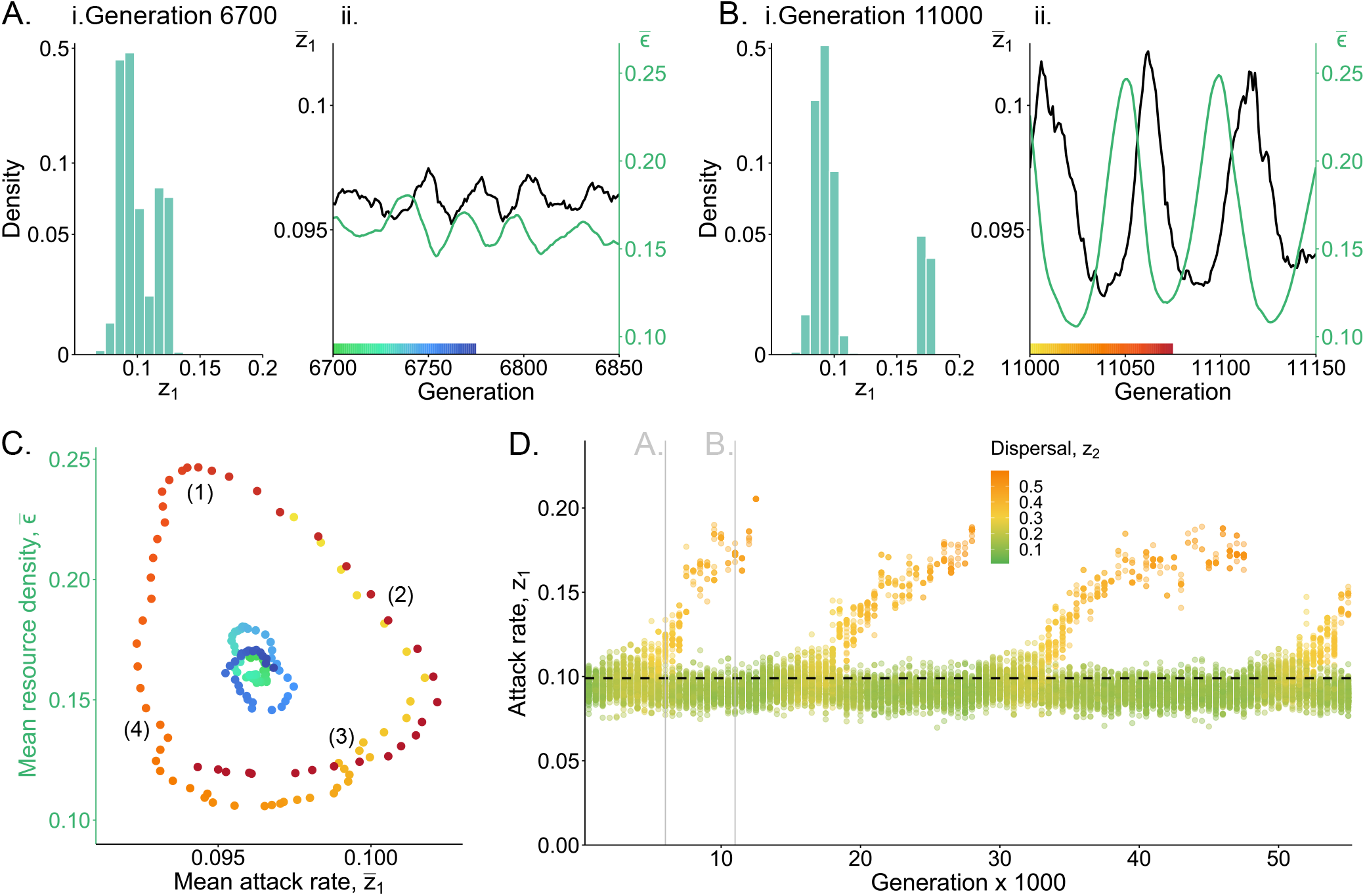
Ecological and evolutionary cycles when the attack rate coevolves with dispersal. Our simulations under disruptive selection are characterised by cycles that occur at different time scales (all plots here are with the same parameters as in Fig. 3B). **A-C:** fluctuations on short ecological time scales (in the order of tens of generations); and **D:** on long evolutionary times scales (in the order of thousands of generations). At the onset of polymorphism when both morphs are weakly diverged (e.g. at generation 6’700, Ai for trait distribution in the population), the population fluctuates between periods with more and fewer overconsumers (in black in Aii), and of lower and higher abundance (in green in Aii). When later on the divergence among morphs is greater (e.g. at generation 11’000, Bi for trait distribution in the population), the amplitude of these fluctuations also increase (Bii). The joint fluctuations in population average attack rate and resource abundance shown in A and B can be mapped onto a phase plot, which is shown in C (blue tones for fluctuations seen Aii, and red tones for fluctuations seen Bii; see x-axes of Aii and Bii for colour legend of time points). This phase plot C shows four phases: (1) when the resource is abundant and consumption is low, favouring an increase in overconsumers; (2) as overconsumers become more frequent, resource abundance falls; (3) when the resource is scarce, overconsumers are counter-selected; (4) once overconsumers are rare, resource abundance increases. As the morphs becomes increasingly differentiated due to disruptive selection, the over-consumer morph becomes increasingly rare, leading to its stochastic extinction in times of low abundance, until it reappears through disruptive selection whereupon the evolutionary cycles starts again (these evolutionary cycles are shown in D, where each point represents an individual, coloured according to its dispersal probability – see figure for legends).

## 5 Discussion

Here, we have extended current theory on the gradual evolution of traits under ecological inheritance to understand how phenotypic variation within populations is moulded by disruptive and correlational selection. Our analyses indicate that ecological inheritance opens three pathways for correlational selection to shape polymorphism and create associations among traits (eq. 16).

The first of these pathways associates traits that have synergistic effects on fitness via the environment (*h*_e×e,*pq*_ (***z***), eq. 17). This is relevant to situations where several traits jointly contribute to the local environment which can be inherited by future generations of relatives. In a naked mole-rat colony for instance, a well-maintained burrow rests on multiple tasks, such as gnawing at the tunnel walls, digging, sweeping substrate, bringing in material to build the nest. According to our model, the tendencies to perform these different tasks may become linked within individuals in two cases: (i) when traits have multiplicative effects on the environment (first term of eq. 17); or (ii) when the environment has non-linear effects on fitness (second term of eq. 17). In the absence of interference among characters, we may expect different traits with beneficial consequences for the environment, such as those contributing to a burrow, to have positive multiplicative effects. In this case, correlational selection favours a positive association between traits and within individuals (under case (i) above). Interestingly, such a positive association has been reported in the naked mole-rat among nest building and burrowing (Siegmann et al., 2021), so that rather than specialising in either of these two different tasks, individuals contribute either more or less to both within colonies. In fact, task specialization and labour division appears to be absent from many animal societies (controlling for sex- or condition-specific effects, Kitchen and Packer, 1999; e.g. in meerkat, Clutton-Brock et al., 2003, wild banded mongoose, Sanderson et al., 2015, and purple-crowned fairy-wren, Teunissen et al., 2020). According to eq. 17 (second term), one situation under which division of labour towards a heritable environmental factor may evolve (i.e. such that correlational selection is negative) is where traits have positive independent effects on the environment, but an improved environment results in diminishing returns on fitness (i.e. such that *∂*^2^*w*_*i*_ /*∂ϵ*^2^ < 0). This would happen for instance when fitness saturates or plateaus with environmental quality, which is a natural assumption for most models (Foster, 2004 for a review on the diminishing returns of helping).

The two other pathways for correlational selection, *h*_r×e,*pq*_ (***z***) and *h*_g×e,*pq*_ (***z***), favour the association between traits that lastingly modify the environment with traits that belong to two broad classes, respectively. One class consists of characters that influence the likelihood that environmental modifications are experienced by downstream relatives (*h*_r×e,*pq*_ (***z***), eq. 21). This should be especially relevant for traits that underlie gene flow as gene flow is the main driver of relatedness. We showed for instance in section 4 that selection readily links dispersal with the attack rate on a local resource, leading to the coexistence of two morphs: a dispersive morph that consumes more and depletes the local resource, and a sessile morph that consumes less and maintains the resource. This positive association between dispersal and attack rate allows overconsumers to preferentially bequeath the patch they deplete to non-relatives, and more frugal individuals to preferentially leave the patch they maintain to relatives. The polymorphism here differs from the social polymorphism described in Mullon et al. (2018) where the social trait has only immediate intra-generational effects, and not inter-generational effects like here. In fact, no polymorphism emerges in our illustrative example when there is no carry-over effects between generations (i.e. when the resource renewal rate *γ* is large such that resource abundance is the same at each generation). This highlights how the emergence of intra-specific trait variation in attack rate is driven by ecological inheritance rather than direct social interactions in our example.

Other than the attack rate, variation in handling time or feeding efficiency can also affect resource abundance (Holling, 1959; Rueffler et al., 2006) and may thus also become linked with dispersal. The expectation from our model is that individuals that have a more negative impact on resource abundance, such as with shorter handling time or greater feeding efficiency, evolve to disperse more readily. In addition to resources, individuals can modify many other environmental factors that are relevant to fitness (Estrela et al., 2019 for review). For example, Drosophila larvae metabolically release nitrogenous waste thereby deteriorating the rearing environment of future larvae (Borash et al., 1998). Microbes can modify the pH of their environment which feeds back on their growth and survival (Ratzke and Gore, 2018). Others can reduce the concentration of toxic metals or antibiotics, improving their substrate (O’Brien et al., 2014; Yurtsev et al., 2016; Frost et al., 2018). The traits that underlie such modifications may thus also become linked to dispersal, leading to kin-biased ecological inheritance. A broad-brush conclusion from our model is therefore that under ecological inheritance, correlational selection may associate dispersal with multiple traits that have environmental effects, leading to the emergence of dispersal syndromes (Ronce and Clobert, 2012). Such syndromes, which have been observed across a wide range of taxa (fish, Cote et al., 2010b; Fraser et al., 2001; mammals, Haughland and Larsen, 2004; lizards, Cote and Clobert, 2007; for reviews: Cote et al., 2010a; Spiegel et al., 2017), are ecologically and evolutionarily significant as they influence the demographic and genetic consequences of movement (Ronce and Clobert, 2012; Edelaar and Bolnick, 2012; Raffard et al., 2021). In the model presented in section 4 for instance, the association between dispersal and attack rate on a resource led to complex dynamics, with cycles occurring both on ecological and evolutionary timescales (Fig. 4).

The second class of traits that selection associates with characters that modify the environment consists of traits whose effects on fitness depend on that environment, i.e. due to “gene-environment” interactions (*h*_g×e,*pq*_ (***z***), eq. 19). Such context- or environment-dependent effects are not uncommon. Traits that are useful during competitive interactions, like conspicuous traits to attract mates (Mappes et al., 1996; Woods Jr et al., 2007; Dougherty, 2021) or fighting appendages like antlers or horns (Emlen, 2008; Miller, 2013), are costly to produce but expression costs likely depend on the environment, at least partly. Indeed, individuals that grow in better conditions or are better provisioned often show more extravagant traits without suffering a greater cost of expression (Mappes et al., 1996; Vehrencamp et al., 1989). The suggestion from our analysis is that context-dependent traits of the sort should become linked to characters that improve the environment when this environment is bequeathed to relatives. This is because such combination of linkage and ecological inheritance allows genes that are good in certain environments to be expressed more often in those environments. This reasoning extends to traits that have indirect context-dependent fitness effects (“indirect g × e interactions” term in eq. 19), for instance favouring the association of helping with traits that deteriorate the environment when the fitness effects of helping increase as the environment decreases in quality (as observed e.g. for cooperative breeding in birds Emlen, 1982). The above considerations should be especially relevant to plastic phenotypes through reaction norms (West-Eberhard, 1989; Stearns, 1989), where one of the evolving trait is the response to the environment and another is modification to this environment. Our model can in fact readily be used to investigate how correlational selection associates environmental response with environmental modification within individuals, and thus help understand the maintenance of variation in plasticity and reaction norms (Pigliucci, 2005).

The gene-environment interactions of our model can also be connected to the so-called process of niche construction, which is “the process whereby organisms actively modify their own and each other’s evolutionary niches” (Laland et al., 2016). This definition is made more explicit by considering the formal models developed by the authors that use it. The typical set-up is a population genetics model with two loci, **E** and **A**, at each of which two alleles segregate (Laland et al., 1996, 1999; Odling-Smee et al., 2003; Silver and Di Paolo, 2006). The current and past allele frequency in the population at locus **E** determines an environmental variable, which in turn determines whether carrying allele *a* or *A* at locus **A** is more beneficial. Such fitness epistasis via the environment is precisely captured by the direct gene-environment interactions in eq. (19) (specifically by the first term where trait *p* is allelic frequency within individuals at locus **A** and trait *q* at locus **E**). Our approach thus encompasses these models. But whereas population genetics models focus on short term evolution through changes in frequency of alleles with potentially large effects, the approach we take here investigates phenotypic evolution in the long term under the constant input of mutation with weak effects. One of our main contributions to this approach is having provided to a way to determine whether gradual evolution in a dispersal-limited population leads to polymorphism in “niche construction” traits owing to disruptive selection and frequency-dependent interactions. The polymorphism here contrasts in two ways with the genetic polymorphism reported in the population genetics models of niche construction (Laland et al., 1996, 1999; Odling-Smee et al., 2003). First, the genetic polymorphism in these previous models is due to specific assumptions about fitness and genetic constraints that create overdominance rather than because of disruptive selection on traits like in our model. Second, while we allow for limited dispersal and local environmental effects, these population genetics models assume that the population is well-mixed and that the same environment is experienced by all individuals in the population (though see Silver and Di Paolo, 2006 for simulations). This entails that a trait cannot be statistically associated to its environmental effect and as a result, there cannot be any selection on a trait’s inter-generational effects (to see this, one can put all relatedness coefficients and their perturbations to zero in our equations; Dawkins, 1982, 2004; Brodie, 2005; Lehmann, 2007, 2008). The inter-generational environmental effects of traits that evolve in those population genetics models are thus a complete by-product of evolution rather than an adaptation.

Like all formal models, ours relies on many assumptions that are relaxed in nature. One is that individuals haploid and reproduce asexually. Provided genes have additive effects on traits, diploidy and sexual reproduction do not influence evolutionary dynamics under directional selection (Geritz and Kisdi, 2000; Rousset, 2004). The emergence of polymorphism due to correlational selection may however depend on the genetic architecture of traits. Where different traits are encoded by separate loci, meiotic recombination breaks the positive genetic linkage favoured by correlational selection. But if the genetic architecture is allowed to evolve (through e.g. recombination modifiers or pleiotropic loci), then correlational selection favours an architecture that allows associations among traits to be heritable (Sinervo and Svensson, 2002), which in turn leads to polymorphism (Mullon et al., 2018). Another useful simplifying assumption we have made is that individuals disperse uniformly among patches so that there is no isolation-by-distance. While isolation-by-distance does not lead to fundamental changes in how selection shapes traits with lasting ecological effects (as shown by analyses of directional selection, Lehmann, 2008), it introduces interesting effects whereby selection depends not only on temporal but also on spatial environmental effects of traits. Finally, we have assumed that patches are of constant size and that traits influence a single environmental variable. We extended our example (section 4) using individual based simulations to consider changing local population size in response to trait evolution (Supplementary Fig. S6 and Supplementary Files for code). Our simulations show similar evolutionary dynamics as those found in the baseline model (Supplementary Fig. S6), which suggests that incorporating explicit demography does not necessarily affect disruptive selection and the nature of polymorphism in this example. But as analyses of directional selection have demonstrated (Lehmann and Rousset, 2010; Lion, 2016, for reviews), demographic structure can in some cases influence qualitatively the evolution of social traits in spatially structured populations. It may thus be interesting, albeit challenging, to extend our analysis of disruptive selection to include demographic fluctuations owing to trait evolution and multiple environmental variables (extending e.g. Ohtsuki et al., 2020 to consider inter-generational effects on patch state). This would be useful to investigate whether the evolution of environmental degradation, such as through resource depletion or release of pollutants, can lead to population extinction (Matsuda and Abrams, 1994; Gyllenberg and Parvinen, 2001; Ferriere and Legendre, 2013).

To sum up, we have investigated the coevolution of multiple traits in a group-structured population when these traits affect the group environment, which is then bequeathed to future generations. We found that such bequeathal provides ground for different types of traits to become linked by selection, with implications for a wide range of traits involved in niche construction, division of labor, dispersal syndromes, condition-dependence and phenotypic plasticity. Our results broadly suggest that ecological inheritance can contribute to phenotypic diversity and potentially lead to complex polymorphism involving multiple traits with long-lasting effects on the environment.

## Supplementary Figures

**Figure S1:**
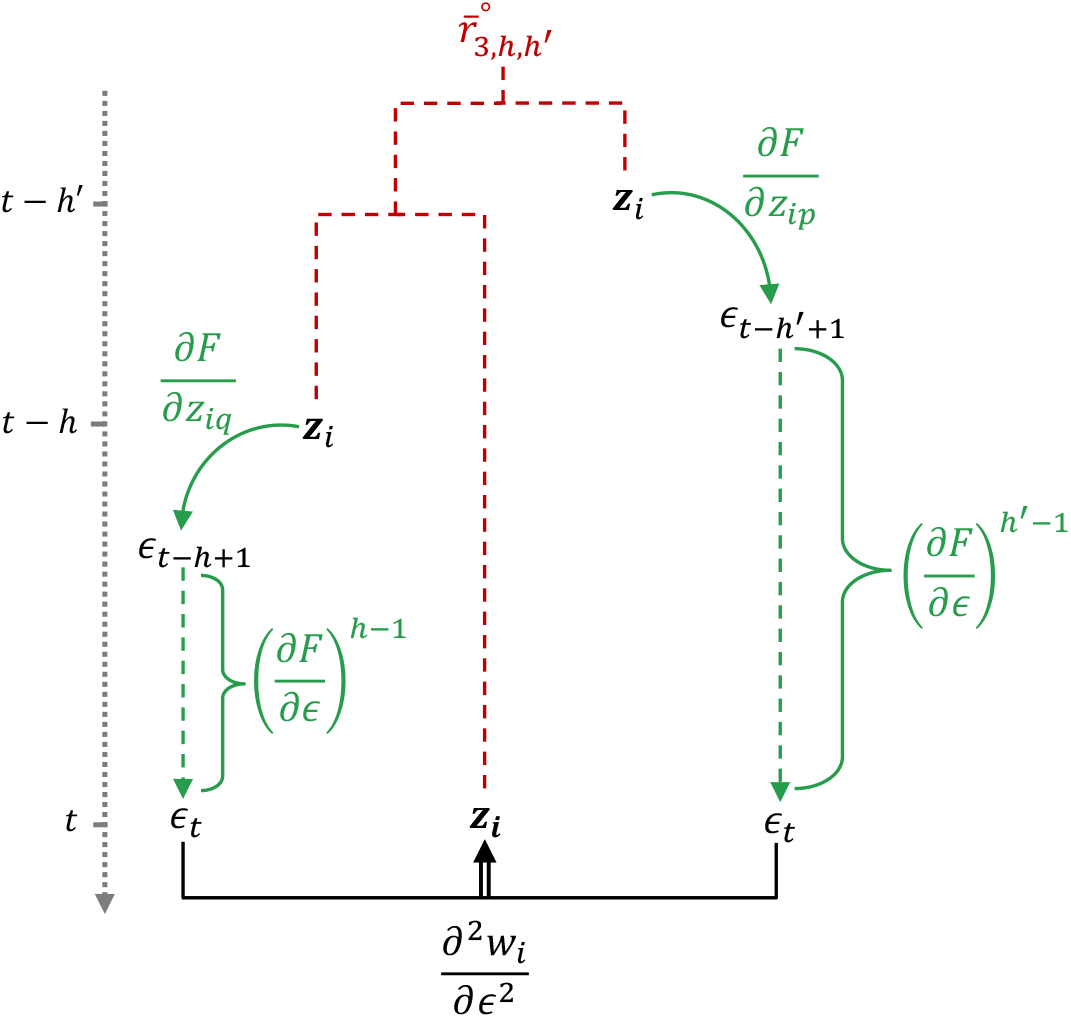
Diagram for synergy due to non-linear fitness effects of the environment, eq. (18) substituted into eq. (17) (second term). To be read similarly to Fig. 1C of main text.

**Figure S2:**
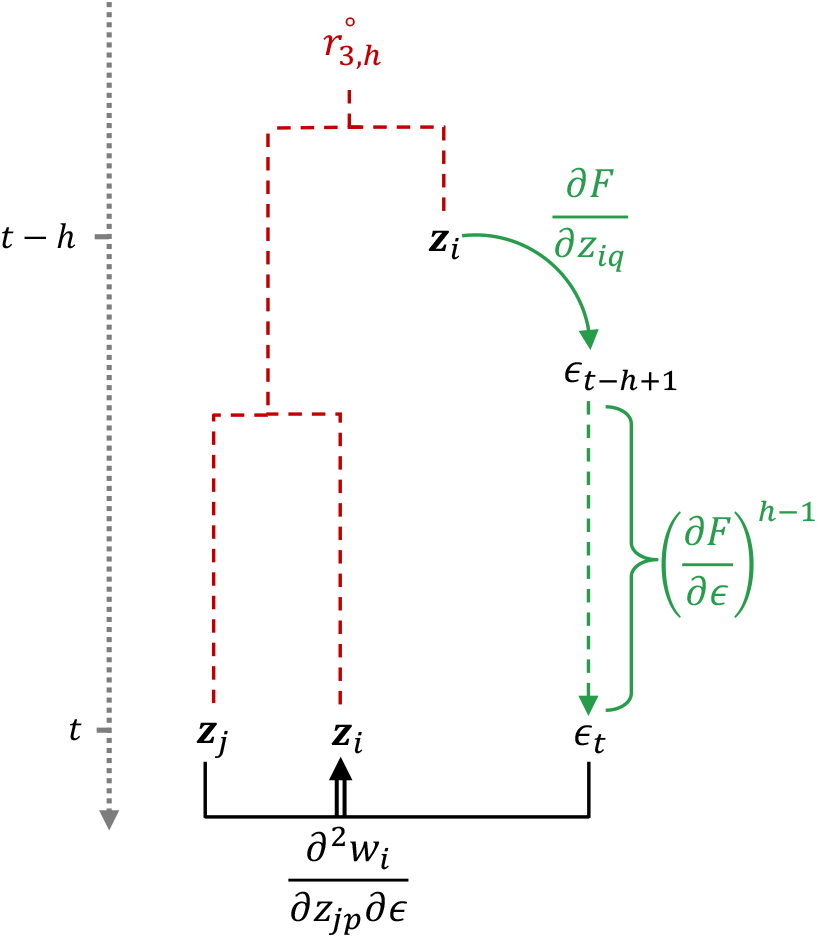
Diagram for indirect gene-environment interactions, eq. (20) substituted into eq. (19) (first term within brackets). To be read similarly to Fig. 1C of main text.

**Figure S3:**
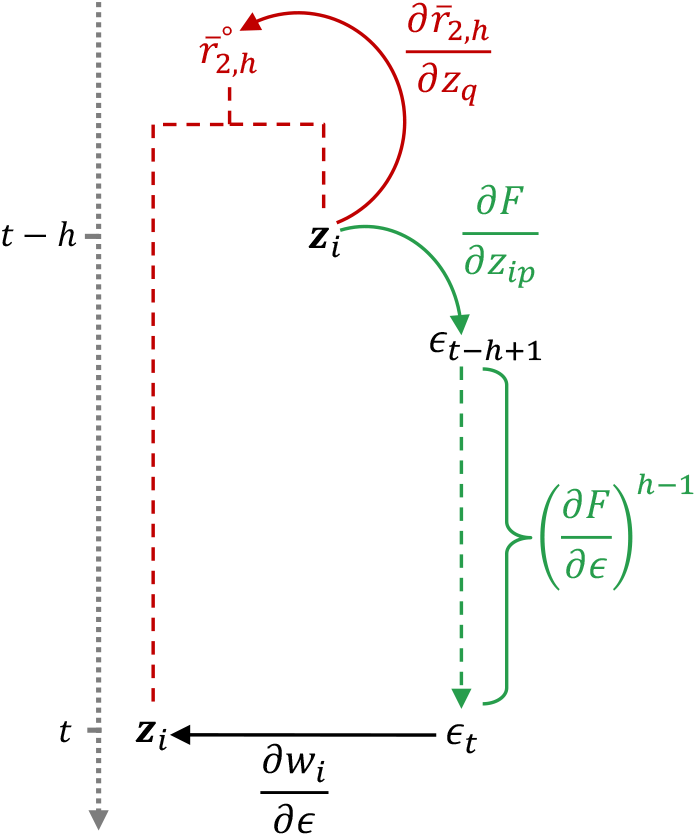
Diagram for biased ecological inheritance, eq. (22) substituted into eq. (21) (first term). To be read similarly to Fig. 1C of main text.

**Figure S4:**
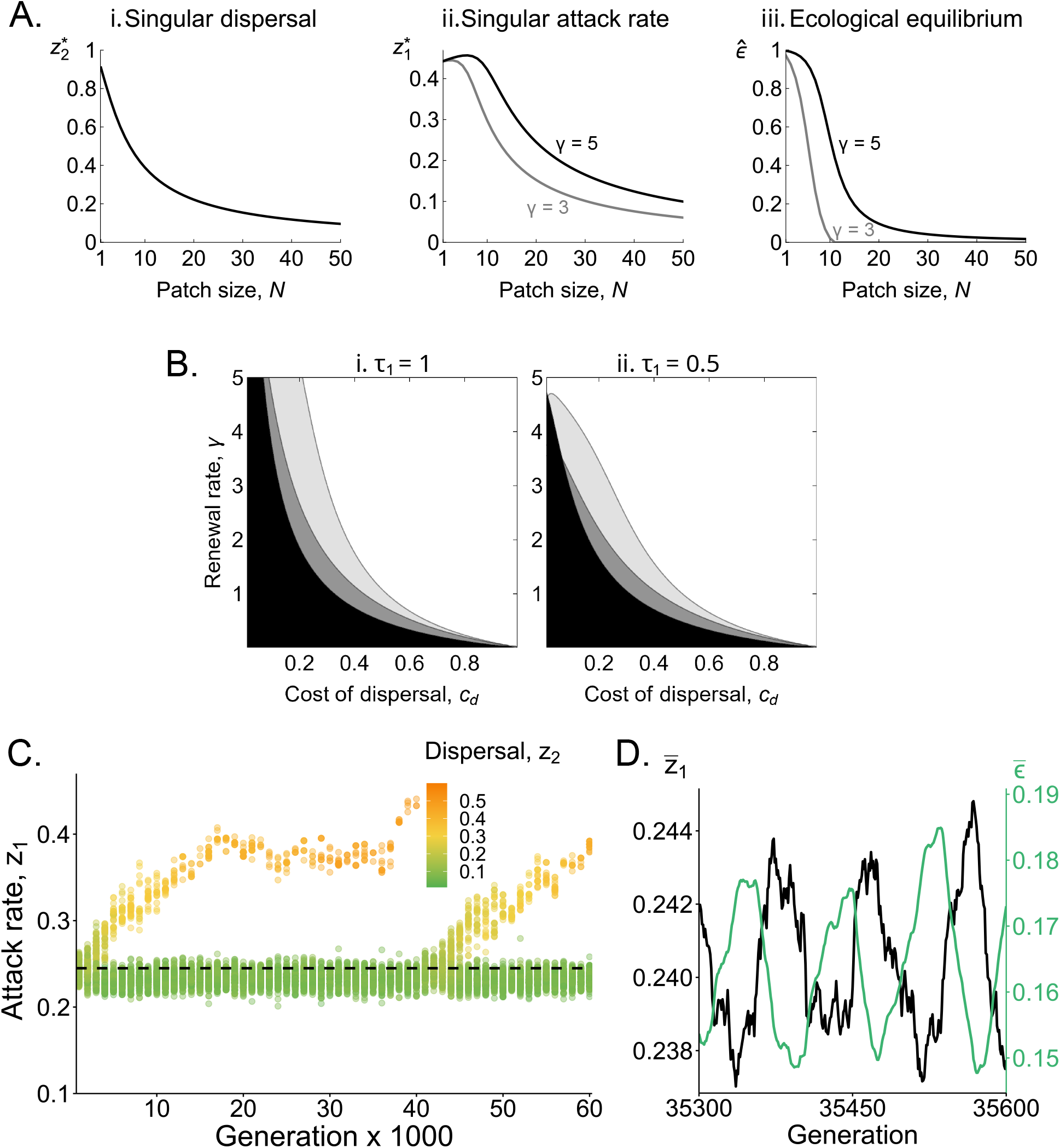
Effect of patch size on the evolution of the attack rate and dispersal. A. Convergence stable dispersal and attack rate, and associated resource level. Obtained from (i) eq. (35); (ii) solving eq. (36) numer-ically for 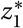; and (iii) from eq. (30) with 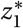 (black: *γ* = 5; gray: *γ* = 3, Other parameter: *τ*_1_ = 5 and *c*_d_ = 0.1).**B. Ecological and evolutionary outcome**. These graphs show the parameter regions where the joint evolution of attack rate and dispersal leads to: resource extinction (in black); disruptive selection (in gray; dark gray for when the attack rate *z*_1_ evolves alone and dispersal is fixed for 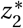); and stabilising selection (in white). Patch size *N* = 20, with (i) *τ*_1_ = 1 and (ii) *τ*_1_ = 0.5. **C. Co-evolutionary dynamics of dispersal and attack, from individual-based simulations**. Each point represents an individual, coloured according to its dispersal probability (see figure for legends). Parameters: *N* = 20, *τ*_1_ = 1, *c*_d_ = 0.1, *γ* = 5. This shows similar albeit slower dynamics as to when *N* = 10 (Fig. 4D). **D. Fluctuations in attack rate and resource abundance on ecological time scale**. Average attack rate (black) and resource abundance (green) in the population over a period of 300 generations, showing similar fluctuations as to when *N* = 10 (Fig. 4Aii and Bii). Same simulation as in C.

**Figure S5:**
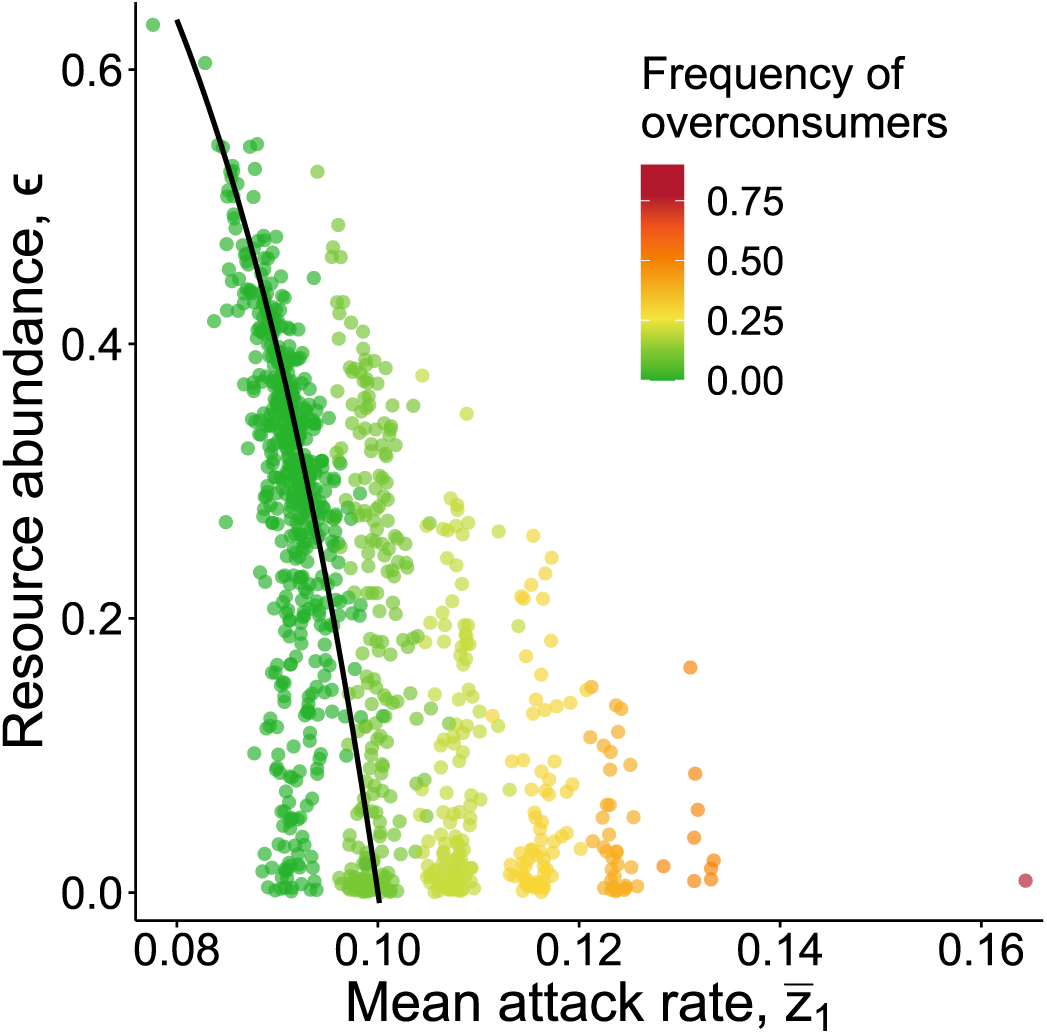
Negative correlation between resource abundance and mean attack rate within patches. From individual-based simulations, same as Fig. 3B at generation 11’000. Each patch is represented as a point according to the abundance of resource it carries and the mean attack rate of the individuals that inhabit it (colour gives the frequency of overconsumers in the patch, see Figure for legend). As expected, this shows that patches with more overconsumers tend to carry fewer resources. In black: expected equilibrium in resource abundance 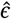 as a function of mean attack rate 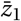 (from eq. 30). For these parameter values, the resource goes to extinction when 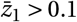 (i.e. at equilibrium, 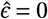 when 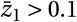).

**Figure S6:**
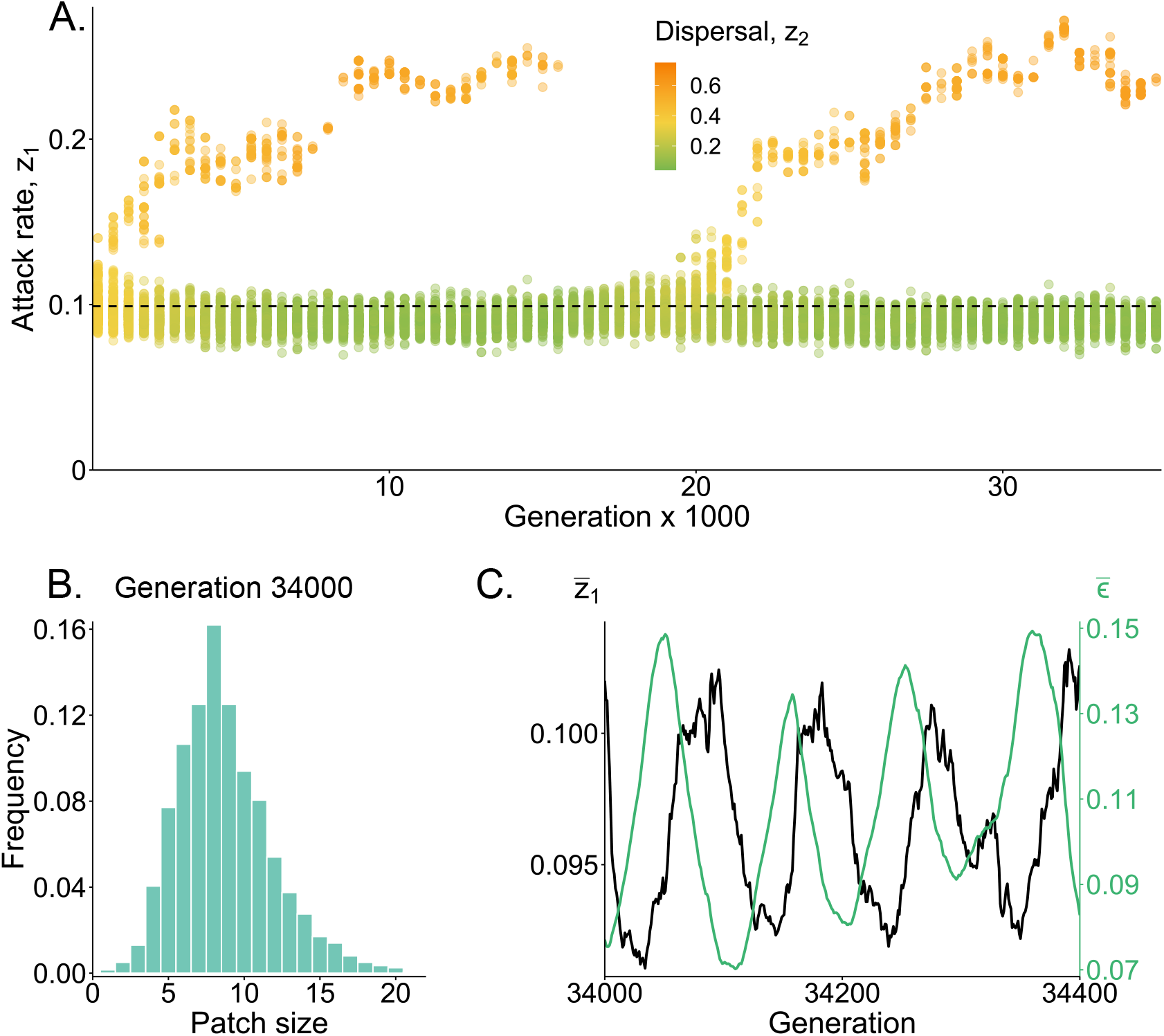
Evolution of dispersal and the attack rate under demographic stochasticity. These graphs show the results of stochastic individual-based simulations where we no longer impose a fixed patch size (i.e. relax the assumption that individuals produce a very large, effectively infinite number of offspring). The algorithm is same as the one described in Appendix F.4, except that instead of performing a fecundity- and dispersal-weighted sampling of *N* individuals from the parental population to form the next generation in each patch (i.e. instead of step ii in Appendix F.4), each parent *i* in each patch *j* now produces *b*_*i j*_ offspring, where *b*_*i j*_ is sampled from a Poisson distribution with mean E[*b*_*i j*_] = *b*_0_ + *f*_*i j*_ /*σ*^2^ (where *b*_0_ = 2 is baseline fecundity; *f*_*i j*_, which is given by eq. F-15 in Appendix F.4, is the effect of traits and consumption on reproduction; and *σ* = 0.03 is inversely proportional to the strength of selection; n.b. these parameters lead to mean fecundity E[*b*_*i j*_] to vary between 2 and approximately 2.1 among individuals in these simulations). Then, each offspring disperses independently with its genetically determined probability *z*_2_ (modelled as a Bernoulli trial with probability *z*_2_ of success). When dispersing, an offspring has a probability 1 − *c*_d_ of surviving dispersal, and if it does, it then has an equal probability of immigrating into any patch other than its original patch. After dispersal, the parental population dies and density-dependent competition occurs in each patch, with each offspring in a patch with *N*_off_ offspring surviving to adulthood with probability 1/(1 + *χN*_off_), where *χ* = 0.008 captures the strength of density dependence (chosen such that the expected patch size in a population monomorphic for the singular strategy is 10). Other parameters: *c*_d_ = 0.1, *γ* = 5, *τ*_1_ = 5, *N*_p_ = 2000. **A. The emergence of sessile scrimpers and dispersive overconsumers**. Each point represents an individual, coloured according to its dispersal probability (see figure for legends). Like where patch size is fixed, we see that as overconsumers become increasingly differentiated, they become rarer until they stochastically get extinct. The population then evolves toward the singular strategy for dispersal and attack rate, and the evolutionary cycles starts again. **B. Distribution of patch sizes at a given generation**. This shows the variation in patch size due to demographic stochasticity. **C. Fluctuations in attack rate and resource abundance on ecological time scale**. Average attack rate (black) and resource abundance (green) showing similar fluctuations as to patch size is fixed (Fig. 4Aii and Bii).

## Appendix

### A Invasion analysis

Our investigations are based on an invasion analysis of rare mutants. To infer on invasion, we use the basic reproductive number as a proxy of invasion fitness (i.e. a quantity that is sign equivalent around one to invasion fitness so that it equally tells about invasion). We detail this basic reproductive number here. For ease of presentation, we consider the case where the phenotype consists of a single trait. The extension to multiple traits is straightforward.

We denote the basic reproductive number of a rare mutant coding for trait *ζ* ∈ **ℝ** in a resident population monomorphic for *z* ∈ **ℝ** by *R*_0_(*ζ, z*). In the island model of dispersal, the basic reproductive number *R*_0_(*ζ, z*) is given by the expected number of successful offspring produced by an individual that is randomly sampled from a local lineage of rare mutants with trait *ζ* in a resident population with trait *z* (where a local lineage consists of all the individuals carrying the mutant who reside in a focal patch; Mullon et al., 2016; Lehmann et al., 2016). We will sometimes refer to such randomly sampled individual from a local lineage of rare mutants as a “representative” mutant for short.

Suppose the mutation appeared as a single copy at generation *t* = 0 in a focal patch. For our model (see section “life-cycle” in the main text), we need to take into account that the fitness of an individual at an arbitrary generation *t* > 0 depends on the environmental state of the patch and that in turn, this state depends on the genetic history of the patch (i.e. the sequence of mutants and residents there have been in the patch since generation *t* = 0). Because the population consists of patches carrying a finite number of individuals, the genetic history of a focal patch is stochastic due to local sampling effects. To capture this history and its stochastic nature, we define *M*_*t*_ ∈ {1, …, *N* } as a random variable for the number of mutants at generation *t* in the focal patch. With the mutation arising at generation *t* = 0 as a single copy, we have *M*_0_ = 1. We write *H*_*t*_ = {*M*_*i*_ }_0≤*i* ≤*t*_ ∈ ℋ_*t*_ for a random collection of the number of mutants in the focal patch from the time the mutation appeared until generation *t* (where ℋ_*t*_ is the countable set of all possible *H*_*t*_) and let Pr(*H*_*t*_) be the probability that history *H*_*t*_ is realized.

Using the above notation and following Mullon et al. (2021) (see their appendix A), the probability *ψ*_*t*_ (*H*_*t*_) that an individual, randomly sampled from the local mutant lineage, lives in the patch at time *t* and that this patch has history *H*_*t*_ can be written as,

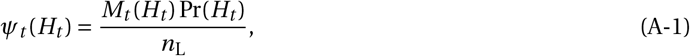

where

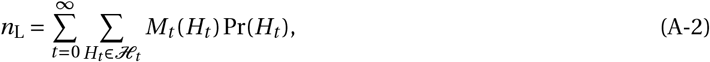

is the expected total size of the local mutant lineage over its lifetime, where we have deliberately stressed the dependence of *M*_*t*_ on *H*_*t*_ by writing it as *M*_*t*_ (*H*_*t*_). Because the population is otherwise monomorphic for the resident *z* and patches are not completely isolated from one another, there is constant immigration of *z*-individuals into the focal patch. As a result, a local mutant lineage eventually vanishes with probability one (i.e. lim_*t* →∞_ Pr(*M*_*t*_ = 0) = 1). In other words, a local mutant lineage has a finite lifetime and so its total size *n*_L_ < ∞ is bounded. For our analyses, it will sometimes be convenient to rewrite (A-1) as

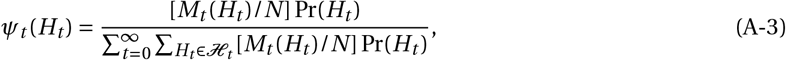

where *M*_*t*_ (*H*_*t*_)/*N* is the mutant frequency in the focal patch at generation *t*.

The probability density function *ψ*_*t*_ (*H*_*t*_) characterises the genetic history preceding a representative mutant. To calculate *R*_0_(*ζ, z*), we further need to specify the individual fitness of such a mutant, which is defined as the expected number of successful offspring of this individual. To do so, let us first define *z*_−*i, t*_ as a vector with *N* − 1 entries collecting the phenotypes of the patch-neighbours of a focal mutant, so that *z*_−*i, t*_ consists of *M*_*t*_ − 1 entries *ζ* and *N* − *M*_*t*_ entries *z*, i.e.,

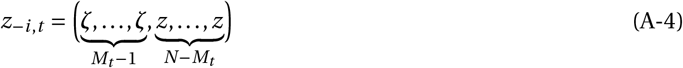

To highlight the dependence of *z*_−*i, t*_ on *M*_*t*_, we will sometimes denote this vector as *z*_−*i*_ (*H*_*t*_). The fitness of an individual also depends on the environmental state of the patch it resides in, which in turn depends on the genetic history of the patch. To capture this, we write the environmental state of the focal patch at generation *t* as *ϵ*(*H*_*t*_). Using the notation introduced here and eq. (4) of the main text, the fitness of a mutant living at time *t* in the focal patch with history *H*_*t*_ can be written as

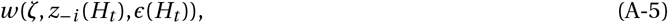

where *ζ* is the (mutant) phenotype of the individual whose fitness is being considered, *z*_−*i*_ (*H*_*t*_) is the vector of phenotypes of its neighbours, and *ϵ*(*H*_*t*_) is the environmental state of its patch.

Finally, we can use the above to write *R*_0_(*ζ, z*), i.e. the fitness of a representative mutant, as

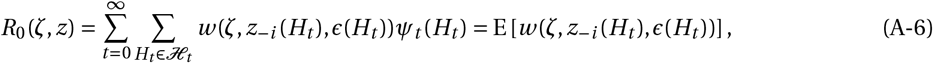

where we have defined the conditional expectation operator,

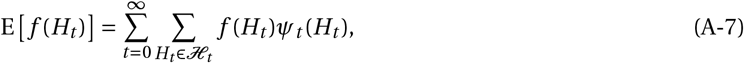

for any function *f* (note that although the expectation does not depend on time as we sum over all *t*, we have kept in our notation the dependence on *t* for *H*_*t*_ within the expectation to highlight how the genetic history of a patch does depend on time). From the theory of multi-type branching processes, it has been shown that if *R*_0_(*ζ, z*) ≤ 1, then the mutant will vanish with probability one (Lehmann et al., 2016). Otherwise, there is a non-zero probability that the mutant invades and replaces the resident. *R*_0_(*ζ, z*) thus constitutes a proxy for invasion fitness *W* (*ζ, z*) of the mutant, equally telling about the nature of selection.

### B Directional selection

Here, we use *R*_0_(*ζ, z*) to re-derive the selection gradient (eq. 10 of the main text) that has been described in previous papers (Lehmann, 2007, 2008; Mullon and Lehmann, 2018). Readers familiar with these aforementioned papers may skip this section. First, we compute the directional selection gradient *s*(*z*) acting on a single trait, which we then generalize to multiple traits.

#### B.1 The two components of the selection gradient

The selection gradient is given by the marginal effect of the trait on the reproductive number, i.e. by

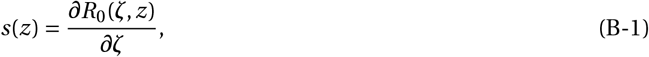

where here and thereafter all derivatives are evaluated under neutrality, i.e., where the population is monomorphic for the resident, *ζ* = *z*, and at environmental equilibrium, 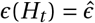, for short. Strictly speaking, this selection gradient defined from *R*_0_(*ζ, z*) is proportional to the one defined from invasion fitness (eq. 6) but we do not distinguish between these gradients as they give the same information about the direction of selection (same for the Hessian). Substituting eq. (A-6) into eq. (B-1) and using the chain rule, we obtain

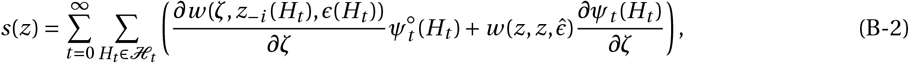

where 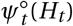 denotes the probability density function *ψ*_*t*_ (*H*_*t*_) under neutrality and thus characterises patch dynamics for a neutral mutant.

To simplify eq. (B-2), we first note that 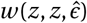 is the fitness of a resident individual in the resident population. Since the population size is constant, this is one, 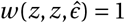. Second, because *ψ*_*t*_ (*H*_*t*_) is a probability density function, it sums to one over its support, i.e. 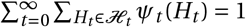. The derivative of this sum with respect to *ζ* therefore vanishes, 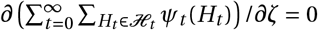. From these observations, the selection gradient simplifies to,

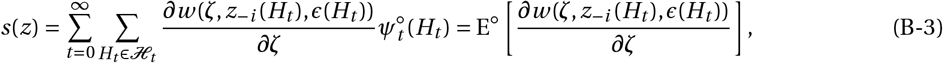

where we use

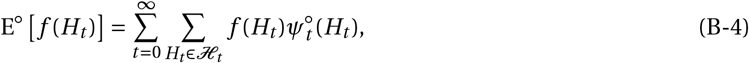

to denote expectation of a function *f* over 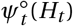.

Next, we re-write the fitness of a focal individual as eq. (4) in the main text,

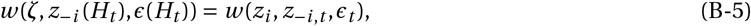

where *z*_*i*_ = *ζ* is the trait of the focal individual, *z*_−*i, t*_ = *z*_−*i*_ (*H*_*t*_) is the vector collecting the traits of the patch neighbours to the focal (eq. A-4), and *ϵ*_*t*_ = *ϵ*(*H*_*t*_) is the environmental state of the focal patch (we drop the time indices in the main text for simplicity). Using the chain rule, we can decompose the derivative of individual fitness as

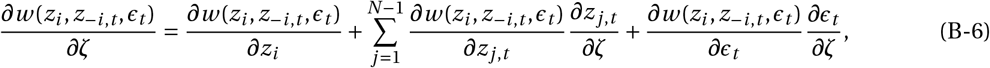

where *z*_*j, t*_ is entry *j* of the vector *z*_−*i, t*_. Since the marginal fitness effect of a trait change in every neighbour *j* ∈ {1, 2,…, *N* − 1} is the same, we can take the derivative out of its sum over *j* in eq. (B-6), and using eq. (A-4) obtain

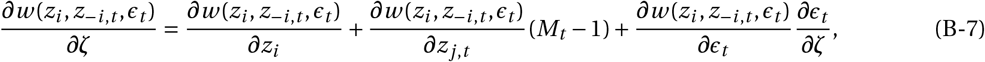

where

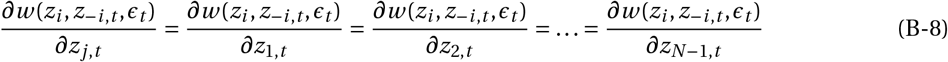

is the indirect fitness effect.

Substituting eq. (B-7) into eq. (B-3), we find that the selection gradient can be decomposed into two components,

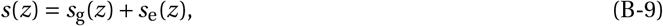

where

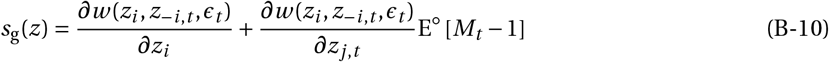

captures selection due to genetic effects of the trait on fitness (hence the subscript g), and

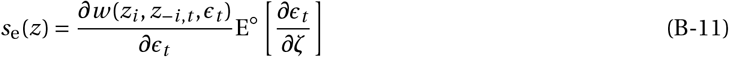

captures selection due to the ecological effects of the trait (hence the subscript e). We detail these components in sections B.2 and B.3 respectively.

#### B.2 Directional selection due to genetic effects

Following straightforward re-arrangements, *s*_g_(*z*) (eq. B-10) turns out to be the standard selection gradient on traits with fitness effects only (Frank, 1998; Rousset, 2004, for textbook treatments),

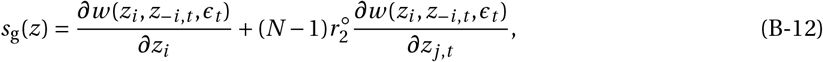

where

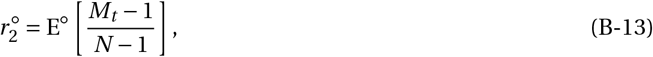

corresponds to the standard coefficient of pairwise relatedness, i.e., the probability that under neutrality, a random individual sampled among the *N* −1 neighbours of a representative mutant is also a mutant (or equivalently, the probability that two individuals sampled without replacement from the same patch at the same generation are identical-by-descent, IBD; eq. D-2 for derivation).

#### B.3 Directional selection due to ecological effects

##### B.3.1 Conditional mutant effect on the environment

To specify *s*_e_(*z*) (eq. B-11), we first derive with respect to *ζ* both sides of

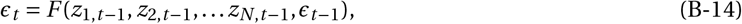

where *F* gives the dynamics of the local environment over one generation (eq. 1 of the main text). We thus obtain

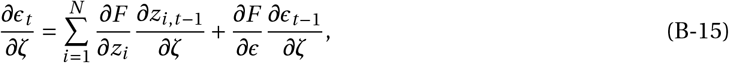

where we have used

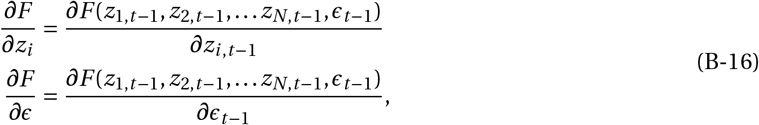

for short. Since there are *M*_*t* −1_ mutants in the patch at generation *t* − 1, eq. (B-15) becomes

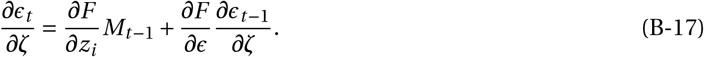

Eq. (B-17) can be viewed as a linear (non-homogeneous) recurrence in *∂ϵ*_*t*_ /*∂ζ*. Since the first mutant appeared at *t* = 0, this recurrence has initial condition *∂ϵ*_0_/*∂ζ* = 0. Solving eq. (B-17) using standard methods, we obtain

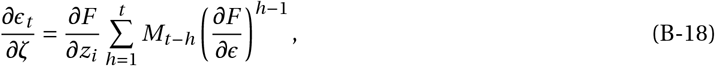

where *M*_*t* −*h*_ is the random variable for the number of mutants in the focal patch, *h* generations before *t*.

##### B.3.2 Unconditional mutant effect on the environment

Eq. (B-18) formalises the notion that the effect of the mutation on the environment at generation *t* depends on the whole (random) sequence of mutants in the focal patch up to *t* (i.e. on *H*_*t*_). As per eq. (B-11), we further need to take the expectation of this effect over the distribution 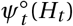 of *H*_*t*_ under neutrality,

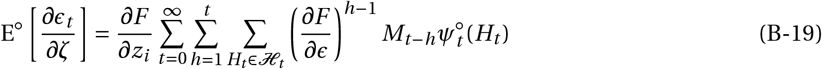

(using eq. B-4). A change of dummy variables in the sums of the right-hand side of this equation allows us to rewrite it as

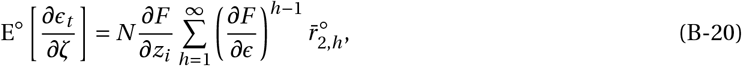

where

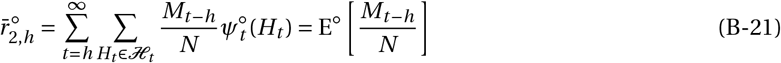

corresponds to the probability that under neutrality, an individual that is randomly sampled from the focal patch *h* ≥ 1 generations before a representative mutant is also a mutant. Alternatively, 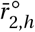 can also be understood as the probability that under neutrality, two individuals sampled *h* generations apart from the same patch are IBD. We compute 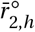 in Appendix D.2.1, obtaining eq. (D-7). Substituting eq. (D-7) into eq. (B-20), we obtain

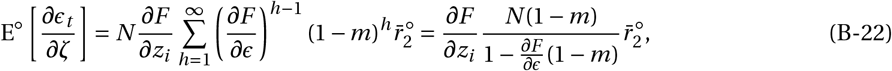

where

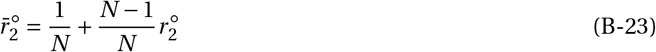

is the probability that two individuals sampled with replacement from the same patch at the same generation are IBD (under neutrality).

#### B.4 Selection gradient vector

Extending the selection gradient to multiple traits is straightforward (Leimar, 2009; Débarre et al., 2014; Geritz et al., 2016). One needs to consider that each trait *p* of the phenotype vector ***z*** = (*z*_1_, …, *z*_*n*_) can influence individual fitness (eq. 4) and the environmental state of the patch (eq. 1). From these considerations and using the same arguments as the ones used for eqs. (B-1)-(B-12), we get

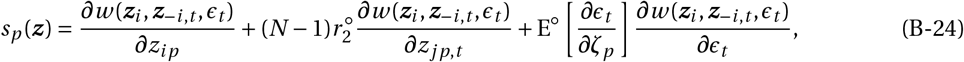

for the selection gradient on trait *p*. Eq. (B-24) corresponds to eq. (10) in the main text, where we drop time-indices and write *z*_*jp,t*_ = *z*_*jp*_ for the trait *p* of a patch-neighbour, *ϵ*_*t*_ = *ϵ* for the state of the patch, *w* (***z***_*i*_, ***z***_−*i, t*_, *ϵ*_*t*_) = *w*_*i*_ for focal fitness, and E^◦^ [*∂ϵ*_*t*_ /*∂ζ*_*p*_] = E^◦^ [*∂ϵ*/*∂z*_*p*_] for the expected effect of a change in trait *p* on the environment experienced by a carrier of this change. Following similar arguments as to those in eqs. (B-15) to (B-22), we find that this latter effect is

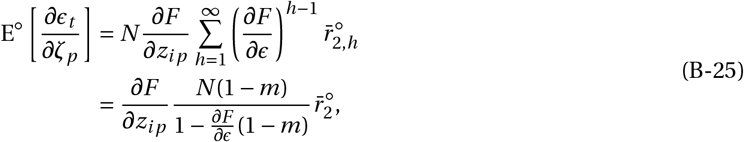

where the first line can be rearranged to read as eq. (11) of the main text, and the second line to row 5 of Table 1.

### C Disruptive selection

In this appendix, we derive the results presented in section 3 of the main text, which shows disruptive and correlational selection. As done for the selection gradient, we first compute the second-order effects of selection,

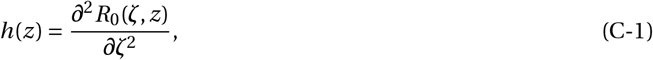

when a single trait is evolving which we then generalize to multiple traits (Leimar, 2009; Débarre et al., 2014; Geritz et al., 2016). With a single evolving trait, *h*(*z*) corresponds to the coefficient of disruptive selection on *z*.

#### C.1 Decomposing the disruptive selection coefficient

##### C.1.1 Neutral and mutant genetic structure

First, we substitute for the reproductive number *R*_0_(*ζ, z*) (eq. A-6) into eq. (C-1) and using the chain rule, we obtain

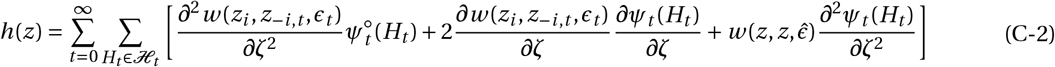

(with eq. B-5). We again use the fact that the sum of *ψ*_*t*_ (*H*_*t*_) over its support is always equal to one (i.e. constant), so that the last term of eq. (C-2) vanishes. This leaves us with two components for disruptive selection,

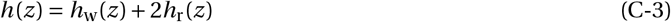

where

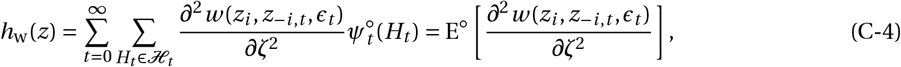

captures the second-order fitness effects of a mutation on fitness averaged over neutral mutant patch dynamics (i.e. over the neutral distribution 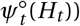, and

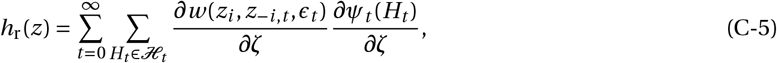

depends on the effect of the trait on mutant patch dynamics (via *∂ψ*_*t*_ (*H*_*t*_)/*∂ζ*). For short hand, we use

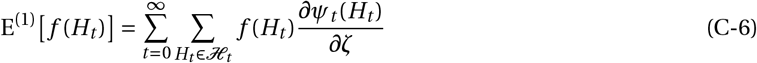

to denote expectation of a function *f* over the perturbation of the probability density distribution *ψ*_*t*_ (*H*_*t*_). With this, eq. (C-5) becomes

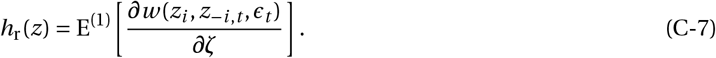

To characterise *h*_w_(*z*) (eq. C-4) further, we first expand the second-order derivatives using the chain rule:

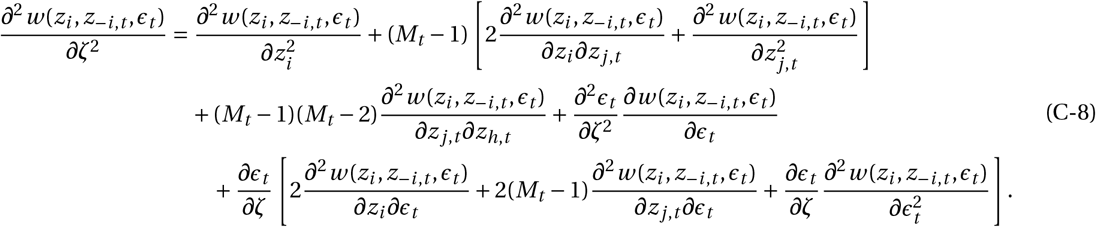

Substituting eq. (C-8) into eq. (C-4), we obtain after some straightforward re-arrangements,

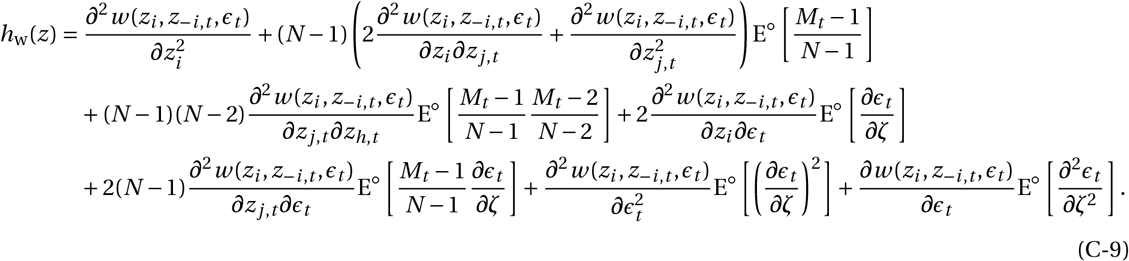

The other term participating to disruptive selection, *h*_r_(*z*) (eq. C-7), meanwhile, can be specified by substituting eq. (B-7) into it, yielding

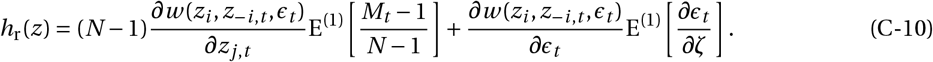

##### C.1.2 Intra- and inter-generational effects

To better contrast our model with previous results that ignore inter-generational ecological effects (Ajar, 2003; Wakano and Lehmann, 2014; Mullon et al., 2016; Mullon and Lehmann, 2019), we re-arrange the coefficient of disruptive selection *h*(*z*) (eqs. C-3, C-9–C-10) as,

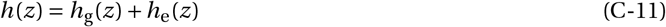

where

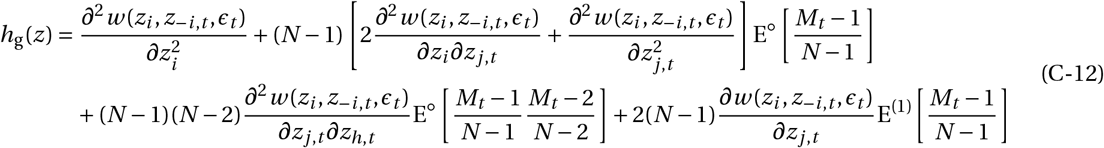

is due to intra-generational effects of traits on fitness, while

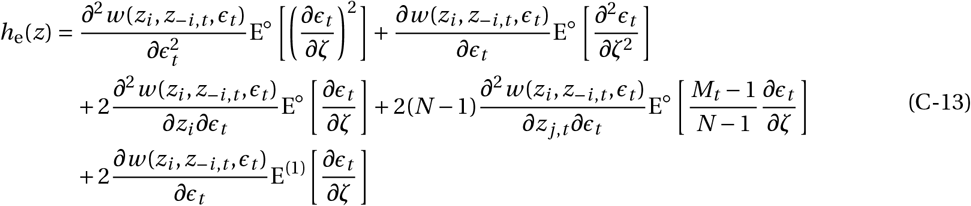

is due to inter-generational effects of traits on fitness. This is the decomposition presented in the main text eq. (14) in the form of correlational selection for the multi-trait case.

#### C.2 Intra-generational fitness effects

Here, we derive eq. (15) of the main text from eq. (C-12).

##### C.2.1 Disruptive selection due to intra-generational fitness effects

From eq. (B-13), we first note that 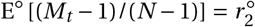 is neutral pairwise relatedness in eq. (C-12). Second,

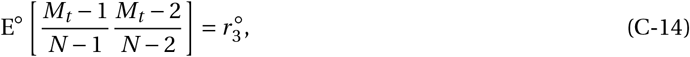

corresponds to the three-way relatedness coefficient under neutrality, i.e. the probability that three individuals sampled without replacement from the same patch are IBD in a monomorphic population. Finally, using eq. (C-6), we have

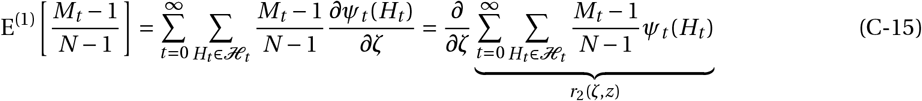

where *r*_2_(*ζ, z*) is the probability that in a mutant patch, a randomly sampled neighbour to a mutant is also mutant. This *r*_2_(*ζ, z*) can thus be thought of as “mutant intra-generational relatedness” and eq. (C-15) as the effect of the trait on such relatedness (see Ajar, 2003; Wakano and Lehmann, 2014; Mullon et al., 2016; Mullon and Lehmann, 2019 for further considerations).

Substituting eqs. (B-13), (C-14) and (C-15) into eq. (C-12), we obtain

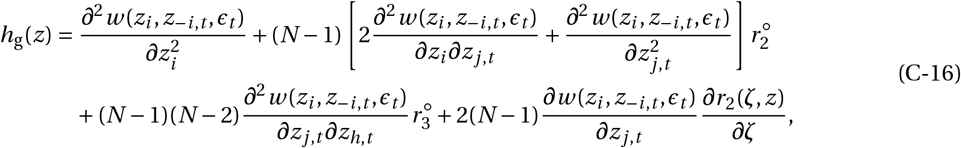

which corresponds to eq. (8) in Ajar (2003) and eqs. (26-27-29) in Wakano and Lehmann (2014).

##### C.2.2 Correlational selection due to intra-generational fitness effects

Eq. (C-16) can be straightforwardly extended to consider the case where multiple traits coevolve. Following previous papers connecting disruptive selection when a single and when multiple traits evolve (Leimar, 2009; Débarre et al., 2014; Mullon et al., 2016; Geritz et al., 2016), we obtain from eq. (C-16) that correlational selec-tion acting on traits *p* and *q* due to intra-generational fitness effect is

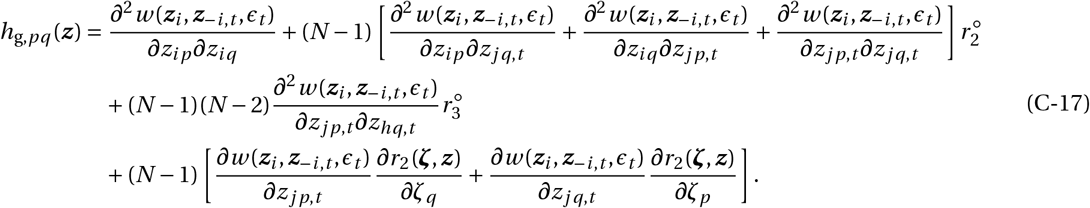

This corresponds to eq. (15) of the main text, where we denote *∂r*_2_/*∂z*_*p*_ = *∂r*_2_(***ζ, z***)/*∂ζ*_*p*_ for the effect of a change in trait *p* on the probability of sampling a mutant among the *N* − 1 neighbours of a focal mutant. We calculate this effect on relatedness in Appendix E.1.

#### C.3 Inter-generational ecological fitness effects

We decompose *h*_e_(*z*) (eq. C-13) into three biologically relevant terms,

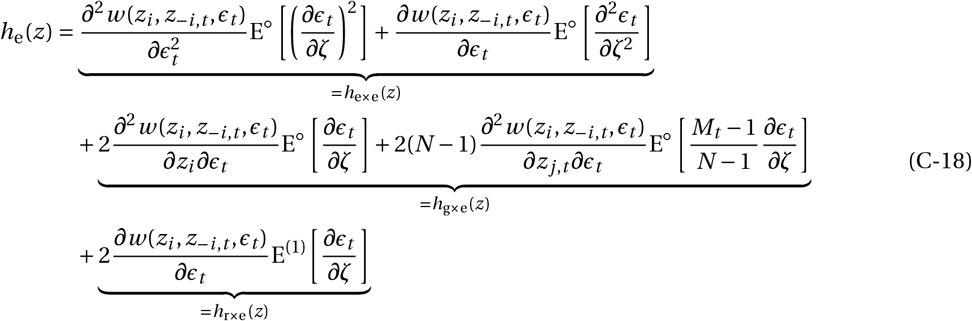

which correspond to three pathways by which the inter-generational effects of traits on fitness may lead to polymorphism. We develop each of these terms below and connect them to the main text equations of section 3.2, which are for the multi-traits case (with eq. C-18 equivalent to eq. 16).

##### C.3.1 Non-linear ecological effects

The first term of eq. (C-18) collects the non-linear ecological effects,

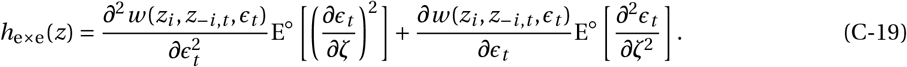

The multi-trait equivalent to eq. (C-19) is eq. (17) of the main text. Below, we unfold both E^◦^ [(*∂ϵ*_*t*_ /*∂ζ*)^2^] and E^◦^ [*∂*^2^*ϵ*_*t*_ /*∂ζ*^2^] to reveal their inter-generational nature.

###### Squared mutant effect on the environment

From eq. (B-18), we have

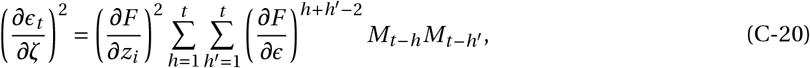

which we substitute into eq. (B-4) to obtain

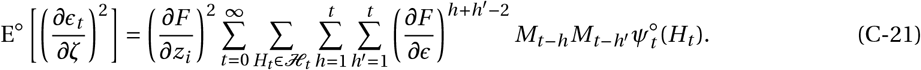

We can change the order of summations in eq. (C-21) to obtain,

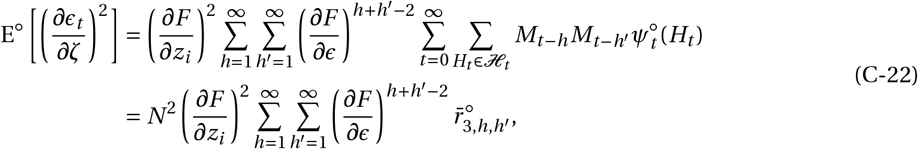

where we defined

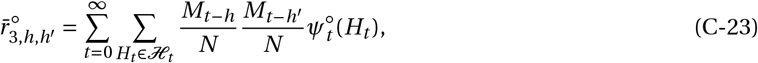

as the probability that under neutrality, two individuals sampled respectively *h* and *h*^′^ generations before a representative mutant are also mutants (all from the same patch).

###### Second order mutant effect on the environment

To specify E^◦^ [*∂*^2^*ϵ*_*t*_ /*∂ζ*^2^] of eq. (C-19), we first derive both sides of eq. (B-14) twice with respect to *ζ*. Using similar argument as for the first order ecological effects (eq. B-14 to eq. B-17), we obtain

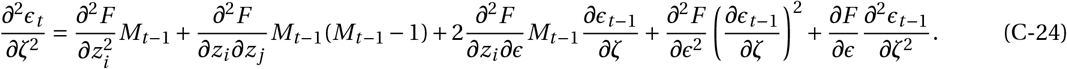

Equation (C-24) can be seen as a linear (non-homogeneous) recurrence for *∂*^2^*ϵ*_*t*_ /*∂ζ*^2^ in *t* with initial conditions *∂*^2^*ϵ*_0_/*∂ζ*^2^ = 0 and *∂ϵ*_0_/*∂ζ* = 0 (since the first mutant appeared at time *t* = 0). Solving eq. (C-24) using standard methods from dynamical systems, we find

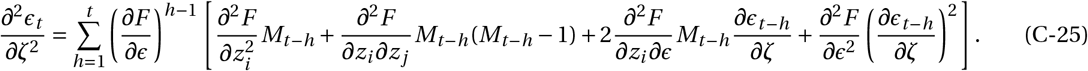

Substituting for *∂ϵ*_*t* −*h*_ /*∂ζ* using eq. (B-18) into eq. (C-25) then yields,

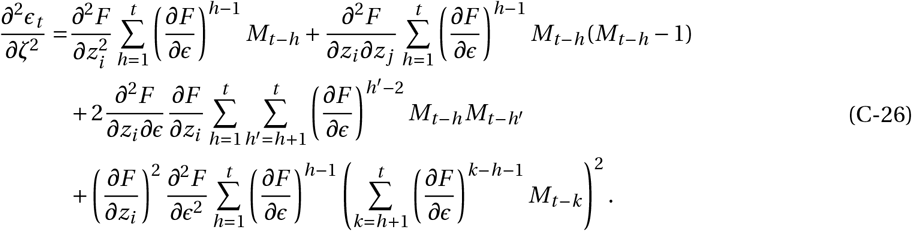

To average these effects and obtain E^◦^ [*∂*^2^*ϵ*_*t*_ /*∂ζ*^2^], we substitute eq. (C-26) into eq. (B-4), which we can rearrange into

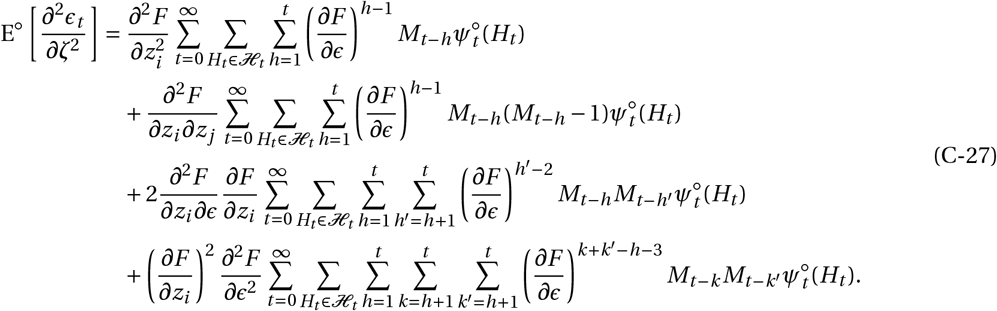

Finally, we can change the summation order in each term of eq. (C-27) to get,

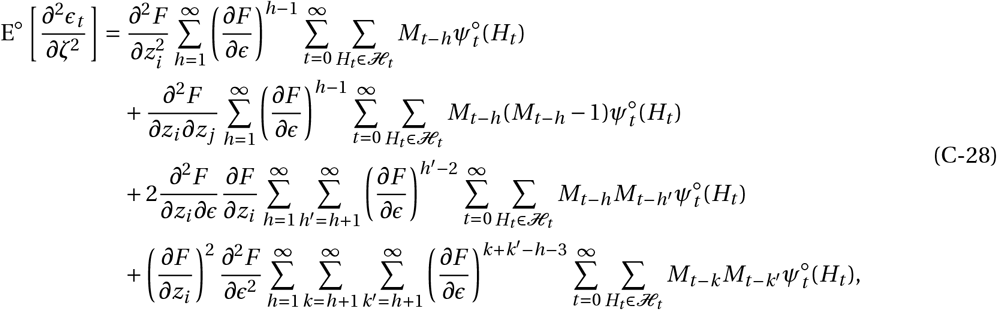

which can be written as

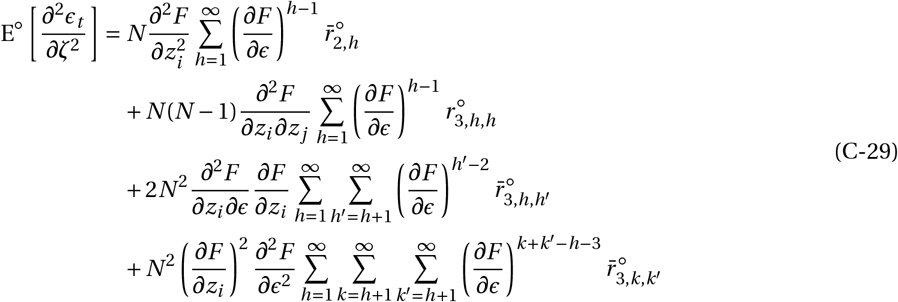

where 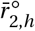 and 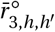 are defined in eq. (B-21) and eq. (C-23), respectively, and were we have defined

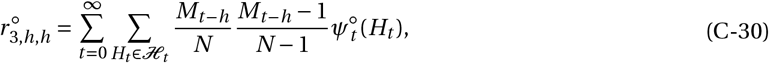

as the probability that under neutrality, two distinct individuals sampled *h* generation before a representative mutant are also mutants.

###### Correlational selection due to non-linear ecological effects

Eq. (C-19) can be extended to obtain the component *h*_e×e,*pq*_ (***z***) of correlational selection on traits *p* and *q* that is due to non-linear ecological effects:

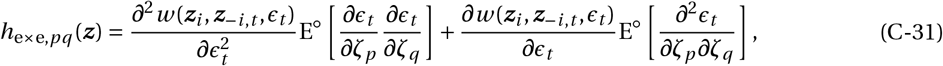

which corresponds to eq. (17) of the main text. From eq. (C-22), meanwhile, we obtain the expected effect of the product of those two traits *p* and *q* on the environment,

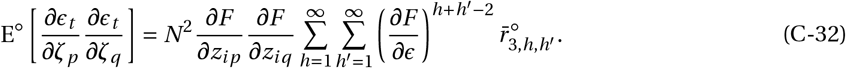

After straightforward re-arrangements, eq. (C-32) corresponds to eq. (18) of the main text. For computational purposes, it will turn out to be convenient to decompose eq. (C-32) as

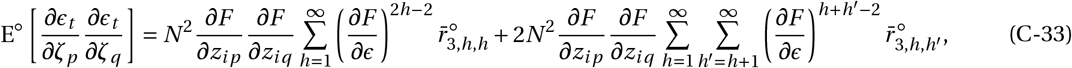

where 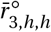 is the probability that under neutrality, two individuals sampled with replacement *h* generations before a representative mutant are also mutants (i.e. defined by eq. C-23, where *h* = *h*^′^).

Similarly, the multi-trait version of eq. (C-29) reads as,

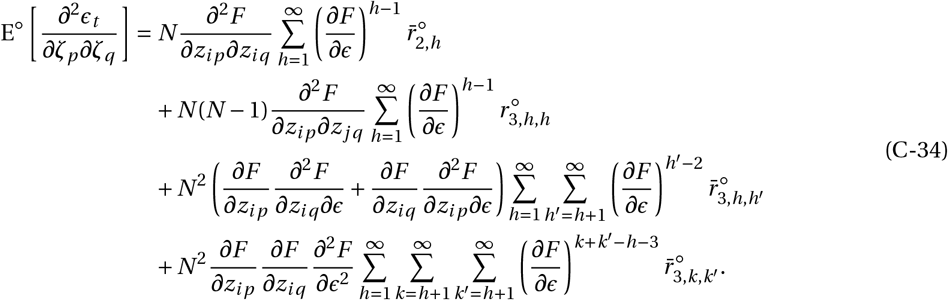

We do not show eq. (C-34) in the main text but see Fig. C-1 for a graphical interpretation. It is convenient to write eq. (C-34) as

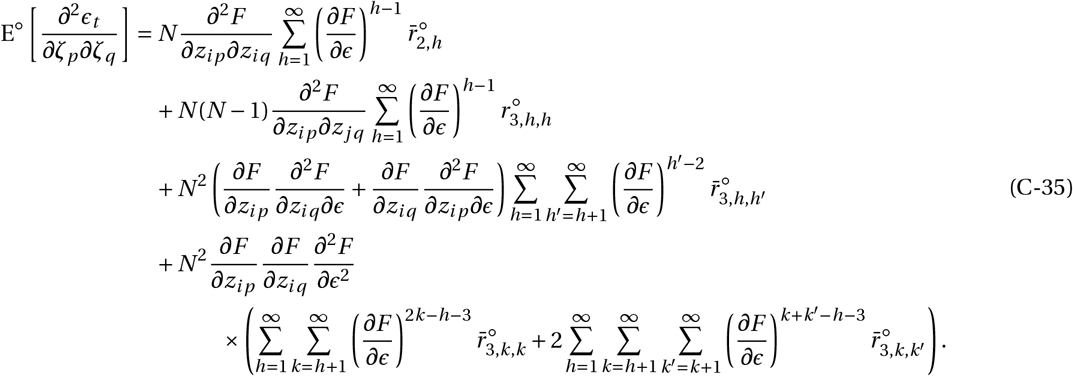

The relatedness coefficients, 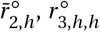, and 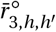, are calculated in Appendix D (eqs. (D-7), (D-9), and (D-20) respectively). We substitute for those into eq. (C-33) and eq. (C-35), yielding rows 6 and 7 in Table 1, respectively.

##### C.3.2 Genes-environment interactions

The second term of *h*_e_(*z*) (eq. C-18) collects the effects that are caused by gene-environment interactions:

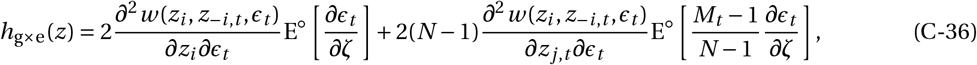

where E^◦^ [*∂ϵ*_*t*_ /*∂ζ*] is given in eq. (B-20) and E^◦^ [(*M*_*t*_ − 1)/(*N* − 1) × *∂ϵ*_*t*_ /*∂ζ*] is computed below.

###### Product between mutant frequency and mutant effect

Using eqs. (B-4) and (B-18) obtains

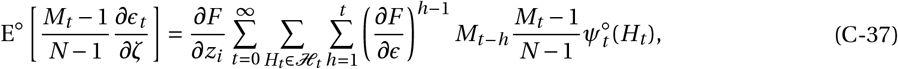

where changing the summation order yields

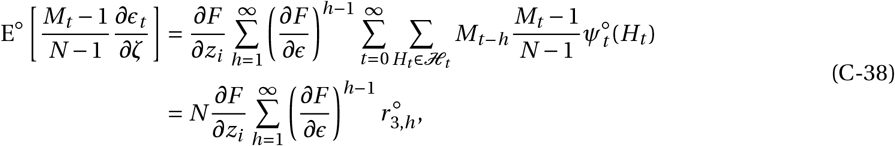

where we defined

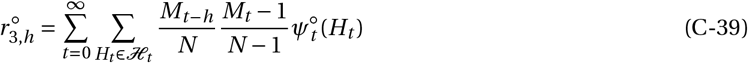

as the probability that under neutrality, two individuals, one sampled among the neighbours of a representative mutant and another sampled *h* generations ago, are also mutants.

**Figure C-1:**
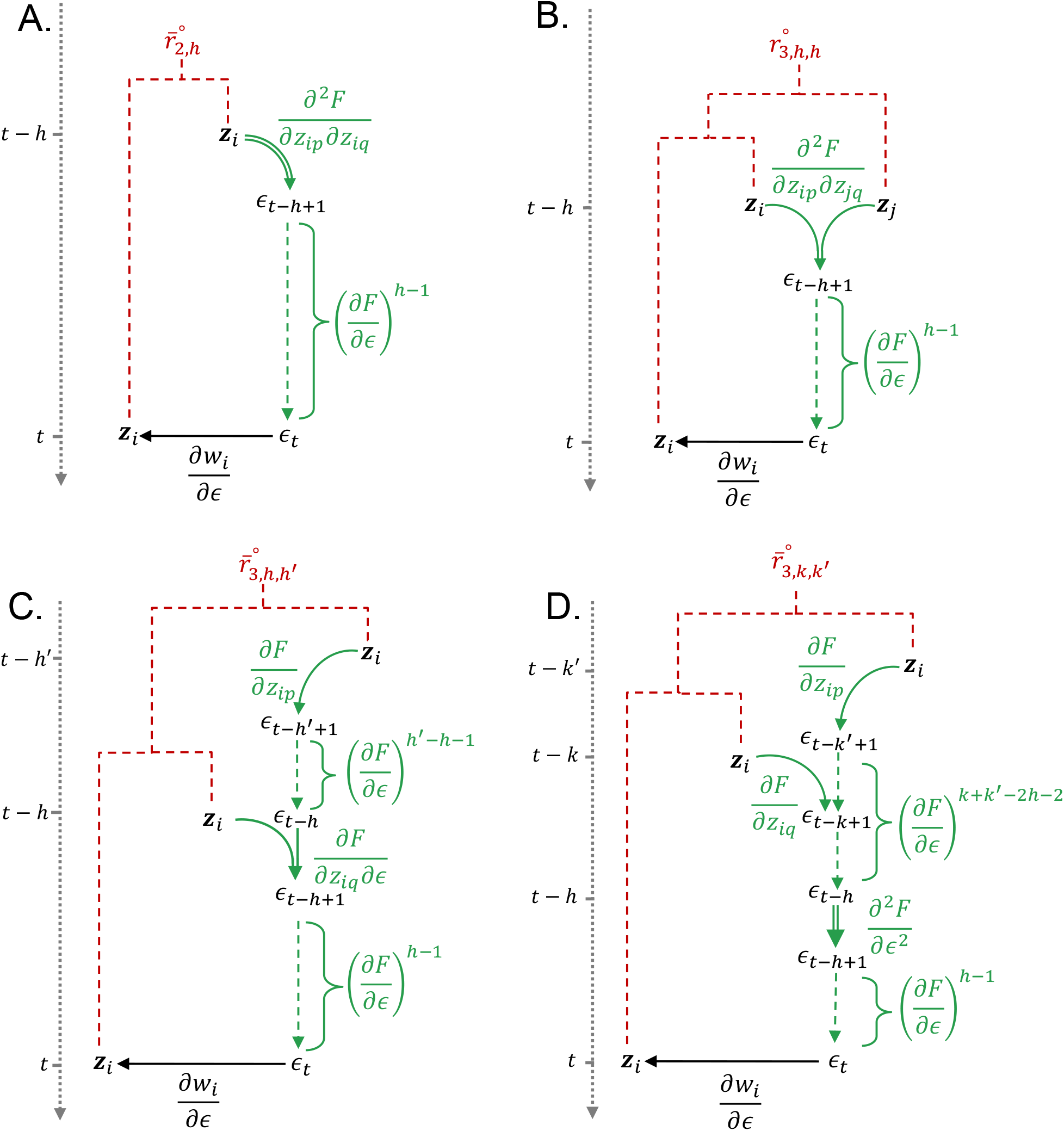
Diagram for expected synergistic effect of traits *p* and *q* on the environment, eq. (C-34) plugged into eq. (C-31). To be read similarly to Fig. 1C of main text. A. corresponds to the first line of eq. (C-34), B to the second line of eq. (C-34), C to the third line of eq. (C-34) and D to the fourth line of eq. (C-34).

###### Correlational selection due to genes-environment interactions

Extending eq. (C-36) to the multi-trait case gives *h*_g×e,*pq*_ (***z***), the correlational selection on traits *p* and *q* that is due to genes-environment interactions:

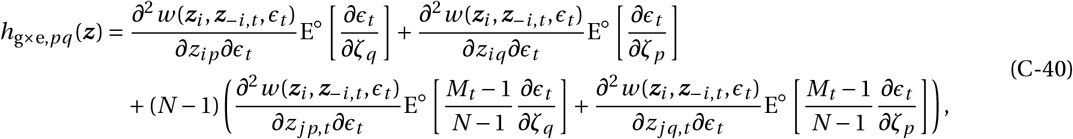

which corresponds to eq. (19) of the main text, where we denote

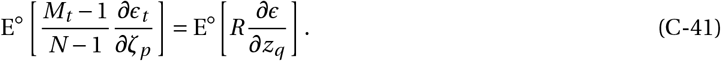

From eq. (C-38), this expectation is

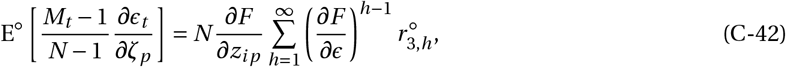

which gives us eq. (20) of the main text. The relatedness coefficient, 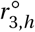, is computed in Appendix D (eq. D-15), which substituted into eq. (C-42) gives row 8 in Table 1.

##### C.3.3 Biased ecological inheritance

The last term of *h*_e_(*z*) (eq. C-18),

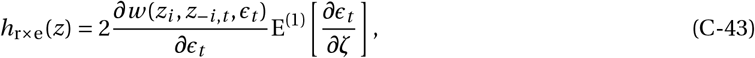

consists of the effects that are caused by biased ecological inheritance, whose multi-trait equivalent is eq. (21) of the main text. Eq. (C-43) is the product of two terms: (i) the fitness effect of a change in the environment; and (ii) the first order effect of the trait on the environment experienced by a mutant. To compute this latter effect, we substitute eq. (B-18) into eq. (C-6) obtaining

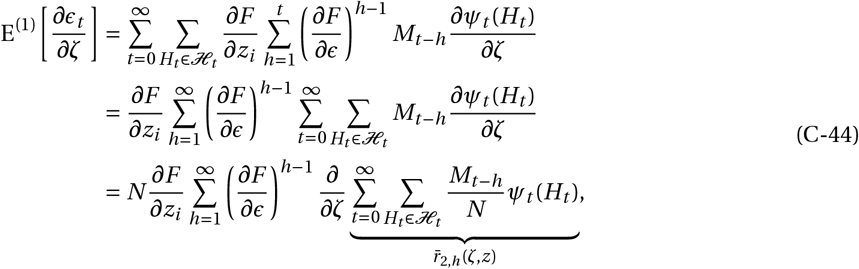

where 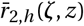 is the probability that given a mutant individual in a mutant patch, a randomly sampled individual from that same patch *h* generations ago is also mutant. Under neutrality, this reduces to 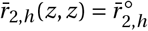.

The derivative of 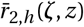 with respect to *ζ* can thus be interpreted on the effect of such trait on intergenerational relatedness.

###### Correlational selection due biased ecological inheritance

Eq. (C-43) can be readily extended to obtain *h*_r×e,*pq*_ (***z***), that is the correlational selection on two traits *p* and *q* that is due to biased ecological inheritance,

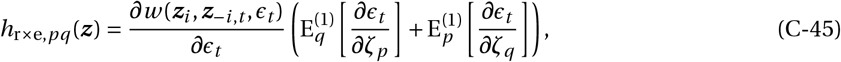

which corresponds to eq. (21) of the main text, where we denote

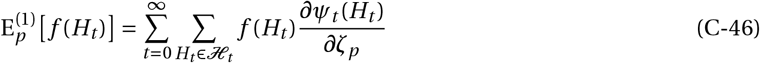

as the expectation of a function *f* over the perturbation of the probability density function *ψ*_*t*_ (*H*_*t*_) due to a change in trait *p* (i.e. the multi-trait version of eq. C-6). Substituting eq. (B-18) into eq. (C-46), we find

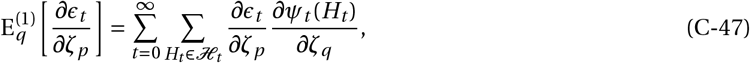

the effect of trait *p* on the environment experienced by a mutant averaged over patch dynamics perturbed by a change in trait *q* (and similarly for 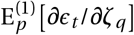. From eq. (C-44), this is

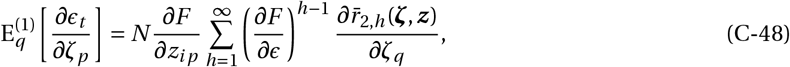

giving us eq. (22) of the main text, where for simplicity we write

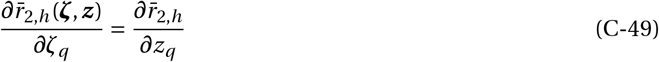

for the effect of a change in trait *q* on inter-generational relatedness. We calculate this effect on relatedness in Appendix E.2 (eq. E-48), which substituted into eq. (C-48) lead to row 10 in Table 1.

### D Relatedness under neutrality

Here, we calculate the neutral relatedness coefficients that are necessary to compute directional, disruptive and correlational selection (and necessary for the rows 1-8 of Table 1 of the main text), using coalescent arguments (e.g., Karlin, 1968; Rousset, 2004).

#### D.1 Intra-generational relatedness

The relevant intra-generational relatedness coefficients for the island model under a Wright-Fisher life cycle are well known (e.g. Ajar, 2003; Ohtsuki, 2010). We rederive them here for the sake of completeness and to illustrate the type of arguments used.

##### D.1.1 Pairwise relatedness

The probability 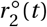 that two individuals living at time *t* in a focal patch are IBD satisfies the recurrence,

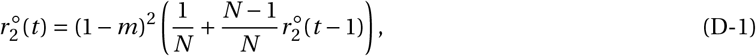

which can be understood as follows. To be IBD, two individuals must be philopatric, which occurs with probability (1 − *m*)^2^. Then with probability 1/*N* they have the same parent (and thus coalesce immediately). Otherwise, with probability (*N* −1)/*N*, they have different parents and therefore are related with probability 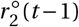.

Solving for the equilibrium 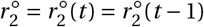 obtains

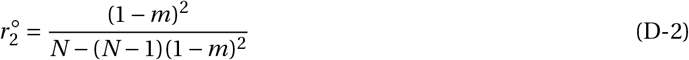

(Rousset, 2004). In turn, the probability that two individuals sampled with replacement from the same patch reads as

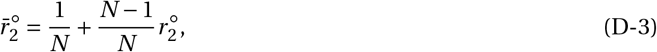

as with probability 1/*N*, the same individual is sampled twice, and with probability (*N* − 1)/*N*, two different individuals are sampled.

##### D.1.2 Threeway relatedness

Using a similar argument as above, the equilibrium probability that three individuals randomly sampled without replacement form the same patch are IBD satisfies,

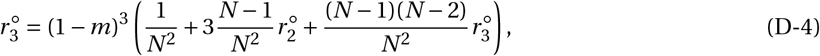

which yields,

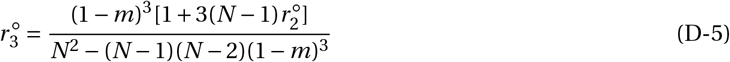

(e.g. Ohtsuki, 2010, eq. 9-10). Meanwhile,

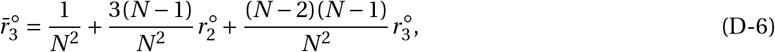

gives the probability that three individuals sampled with replacement at the same generation from the same patch are IBD.

#### D.2 Inter-generational relatedness

Inter-generational relatedness is computed the same way as intra-generational (Lehmann, 2007).

##### D.2.1 Pairwise inter-generational relatedness

The probability that two individuals sampled *h* > 0 generations from the same patch are IBD is

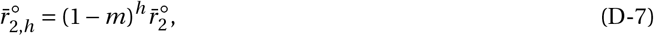

(eq. 6 in Lehmann, 2007). To understand this, consider the lineage back in time of a randomly sampled individual. With probability (1−*m*)^*h*^, this lineage remains in the focal patch for *h* generations and in turn coalesces with the lineage of another individual from that generation with probability 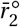.

##### D.2.2 Threeway inter-generational relatedness

Computing disruptive and correlational selection requires three further inter-generational relatedness coefficients that involve three individuals. These coefficients have so far not been characterized in the literature.

###### (i) IBD between one individual at generation *t* and two patch neighbours at generation *t* −*h*

We first compute 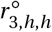 (eq. C-30),

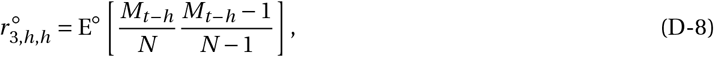

which is the probability that one individual at some random generation *t* and two patch neighbours at generation *t* − *h*, all randomly sampled from the same patch are IBD. This probability is given by,

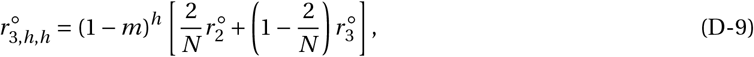

which can be understood as follows. Consider the lineage of the individual from generation *t* back in time. This lineage coalesces with an individual *h* generations ago in the same patch with probability (1 − *m*)^*h*^. Then, with probability 2/*N*, this direct ancestor is one of the two (distinct) individuals sampled at generation *t* − *h*, in which case all three individuals are IBD with probability 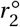. With complementary probability 1 − 2/*N*, this direct ancestor is not one of the two sampled individuals, in which case all three individuals are IBD with probability 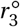.

###### (ii) IBD between two patch neighbours at generation *t* and one individual at generation *t* − *h*

Next, let us calculate 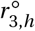 (eq. C-39),

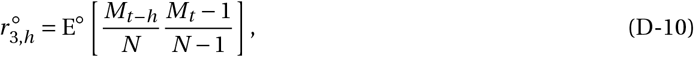

which is the probability that two patch neighbours at generation *t* and a third individual randomly sampled at generation *t* − *h* are all IBD. We need to consider two cases: either (1) the lineages of the two individuals sampled at *t* remain in the focal patch without coalescing within *h* generations; or (2) their lineages coalesce into one common ancestor within *h* generations and their common lineage remains in the focal patch (Fig. D-1). To reflect these two pathways, we decompose 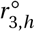 as

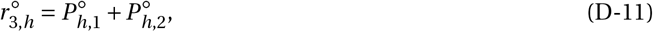

where 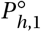 is the probability of coalescence in case (1) and 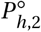 in case (2) (see Fig. D-1 for a graphical representation).

**Figure D-1:**
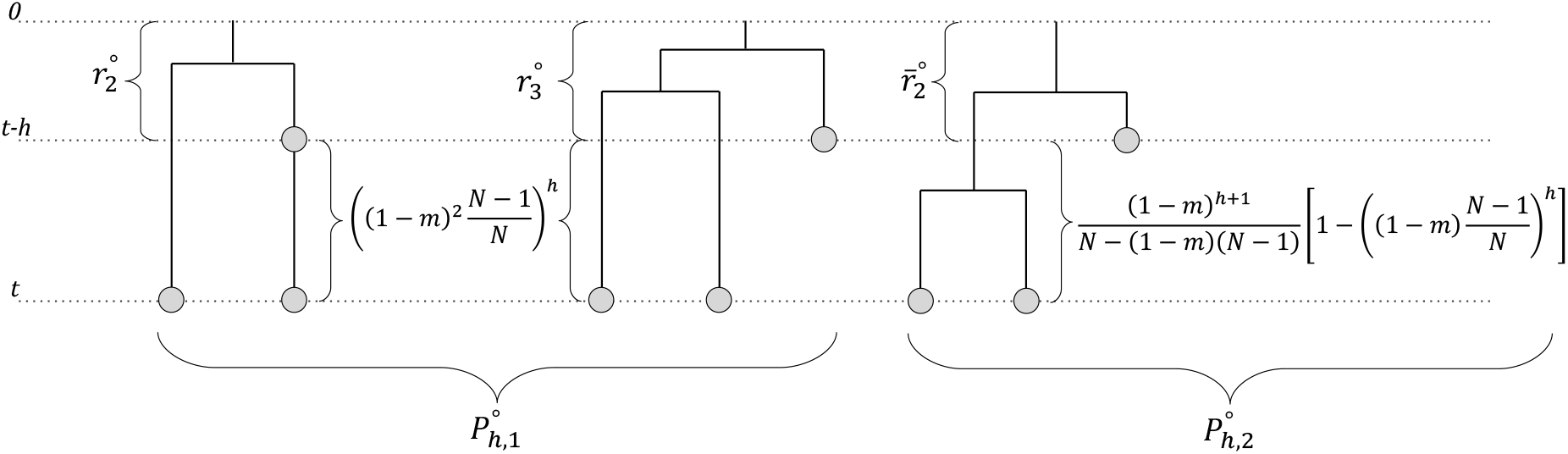
Decomposition of 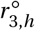. On the left, 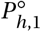 corresponds to scenario (1) (eq. D-12), where the two lineages of the individuals sampled at generation *t* do not coalesce within *h* generations while staying in the focal patch. On the right, 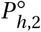 corresponds to scenario (2) (eq. D-14), where the two lineages of the individuals sampled at generation *t* coalesce within *h* generations.

To characterise 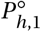, consider the two individuals from generation *t* and their lineages back in time (Fig. D-1, left and middle). With probability (1 − *m*)^2^(*N* − 1)/*N*, these lineages remain in the same patch but do not coalesce within one generation. So with probability [(1 − *m*)^2^(*N* − 1)/*N*]^*h*^, the two lineages remain in the focal patch without coalescing over *h* generations. Assuming this is the case, a randomly sampled individual from the focal patch at generation *t* − *h* generations is a direct ancestor to either individuals with probability 2/*N*, in which case all three individuals are IBD with probability 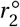 (Fig. D-1, left). With complementary probability 1 − 2/*N*, however, a randomly sampled individual from the focal patch at generation *t* − *h* generations is not a direct ancestor to either individuals, in which case they are all IBD with probability 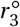 (Fig. D-1, middle). These arguments lead us to

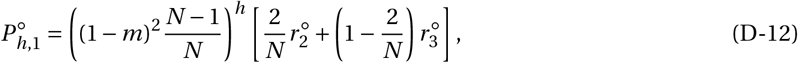

for the first term of 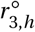.

For 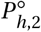, consider again the two individuals from generation *t* and their lineages back in time (Fig. D-1, right hand side). The probability that their lineages coalesce at generation *t* − *i* (1 ≤ *i* ≤ *h*) and further remain in the focal until generation *t* − *h* is given by

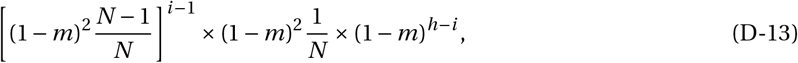

as with probability [(1 − *m*)^2^(*N* − 1)/*N*]^*i* −1^ the lineages remain in the focal patch without coalescing for *i* − 1 generations, then coalesce in one generation with probability (1 − *m*)^2^/*N*, and their common lineage remains in the focal patch until generation *t* − *h* with probability (1 − *m*)^*h*−*i*^. In turn, a randomly sampled individual from the patch at generation *t* − *h* belongs to that same lineage with probability 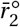. Summing over all possible 1 ≤ *i* ≤ *h*, we thus obtain,

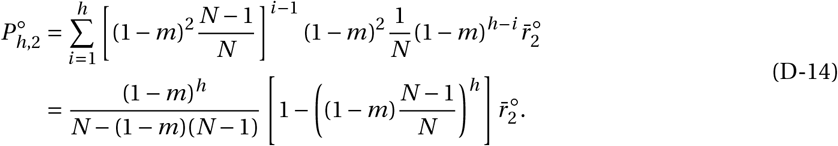

Plugging eqs. (D-12) and (D-14) into eq. (D-11) finally leads us to

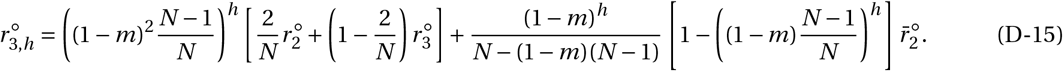

###### (iii) IBD between individuals at generations *t, t* − *h* and *t* − *h*^′^

Finally, we calculate 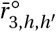 (eq. C-23),

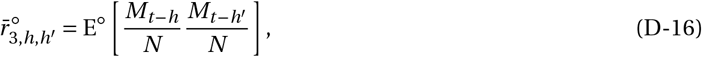

which is the probability that individuals randomly at generations *t, t* − *h* and *t* − *h*^′^ (*h* > 0 and *h*^′^ > 0) from the same patch are all IBD. It is useful to distinguish two cases,

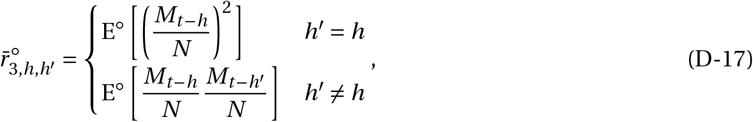

which we consider separately. The case where *h*^′^ = *h* reads as

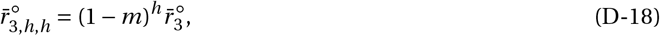

as with probability (1 − *m*)^*h*^ an individual living in the focal patch at generation *t* has a direct ancestor in that same patch at generation *t* −*h*. In turn, this direct ancestor is IBD with two individuals sampled randomly with replacement at generation *t* − *h* with probability 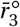. For the case where *h*^′^ ≠ *h*, let us assume without loss of generality that *h*^′^ > *h*. We then have

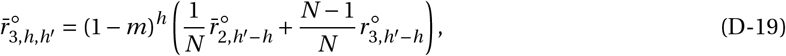

where 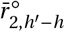 is given by eq. (D-7) and 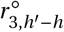 by eq. (D-15). To understand eq. (D-19), consider the lineage of an individual living in the focal patch at generation *t* back in time. With probability (1 − *m*)^*h*^, this individual has a direct ancestor in that same patch at generation *t* −*h*. In turn, with probability 1/*N* this direct ancestor is the randomly sampled individual of generation *t* − *h*, in which case these are IBD with a third from generation *t* −*h*^′^ (a further *h*^′^ −*h* generations ago) with probability 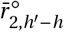. Otherwise, with probability (*N* −1)/*N* the direct ancestor is not the randomly sampled individual of generation *t* − *h*, in which case this randomly sampled individual, the direct ancestor, plus another individual from generation *t* − *h*^′^ are all IBD with probability 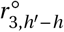.

We thus have altogether

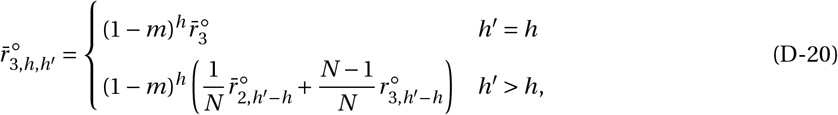

where 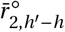 is given by eq. (D-7) and 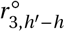 by eq. (D-15).

### E Effect of selection on relatedness

In this appendix, we characterize the effect of selection on intra- and inter-generational relatedness, *∂r*_2_(*ζ, z*)/*∂ζ* (which appears in eq. C-16) and 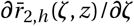 (which appears in eq. C-44), and necessary for the rows 9-10 of Table 1 of the main text. For the ease of presentation, we do so when the phenotype consists of a single trait. The extension to multiple traits is straightforward.

#### E.1 Intra-generational relatedness

Intra-generational relatedness under selection has been derived in previous papers in the absence of ecological inheritance (Ajar, 2003; Wakano and Lehmann, 2014; Mullon et al., 2016). Here, we extend these derivations to include such inheritance. As a starting point, we substitute eq. (A-3) into eq. (C-15) to obtain

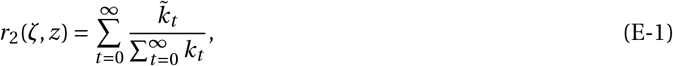

where

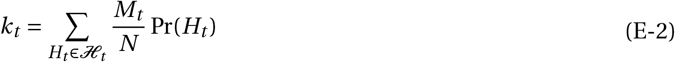

is the probability that an individual randomly sampled from a mutant patch at generation *t* is mutant, and

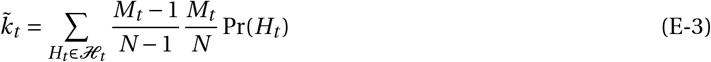

is the probability that two individuals randomly sampled without replacement from a mutant patch at generation *t* are both mutants. Both *k*_*t*_ and 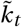 depend on mutant and resident trait values, *ζ* and *z*, but we do not write such dependency explicitly to avoid notational clutter. Taking the derivative of eq. (E-1) with respect to *ζ* and evaluating at *z*, we get

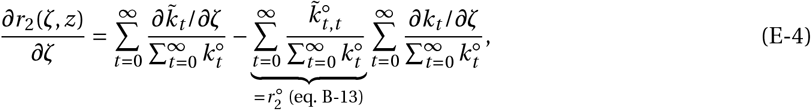

using the quotient rule.

We can unpack 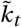 (eq. E-3) using conditional probability as

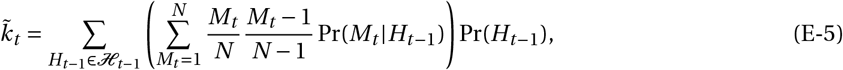

where Pr(*M*_*t*_ |*H*_*t* −1_) is the probability that there are *M*_*t*_ mutants in the focal patch at generation *t* given the history *H*_*t* −1_ of this patch up to generation *t* − 1. Expanding eq. (E-5) then yields,

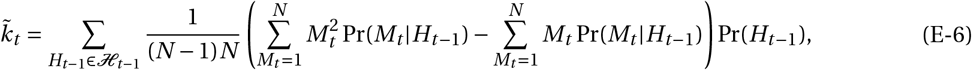

where 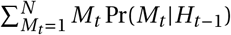 and 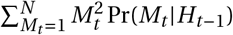 are the first and second moments of the distribution for the number of mutants at generation *t* given *H*_*t* −1_, respectively. To characterise these moments, let us first decompose individual fitness of a mutant at generation *t* − 1 (from eq. A-5) as

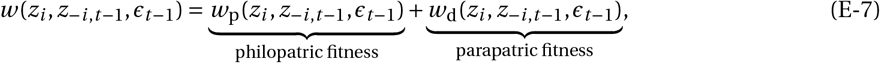

where *w*_p_(*z*_*i*_, *z*_−*i, t* −1_, *ϵ*_*t* −1_) is the number of offspring that remain in their natal patch, and *w*_d_(*z*_*i*_, *z*_−*i, t* −1_, *ϵ*_*t* −1_) those that establish in other patches (and where also *z*_*i*_ = *ζ* and *z*_−*i, t* −1_ is a *N* − 1 vector with *M*_*t* −1_ − 1 entries consisting of *ζ* and *N* − *M*_*t* −1_ of *z*, see eq. A-4). Note that under neutrality (i.e. *ζ* = *z*), these fitness components reduce to

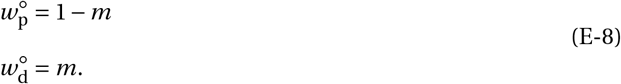

We can then use the fact the life-cycle is Wright-Fisher, so that the number of mutants at generation *t* in a mutant patch follows a binomial distribution with parameters

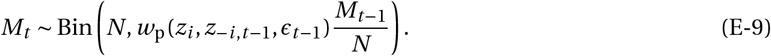

Hence, the two first moments of the mutant distribution are given by

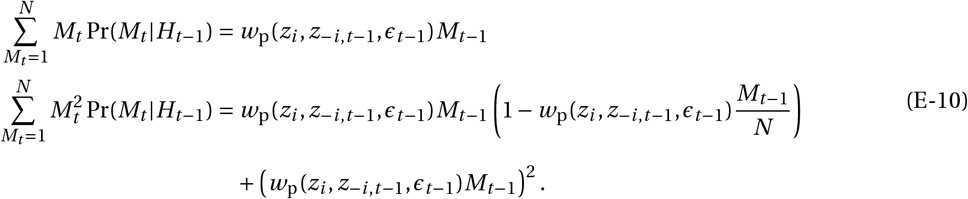

Substituting eq. (E-10) into eq. (E-6), we thus obtain

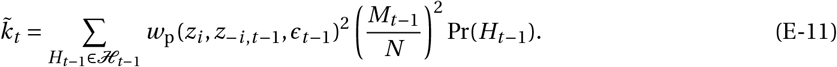

Taking the derivative of eq. (E-11) and using eq. (E-8), we obtain

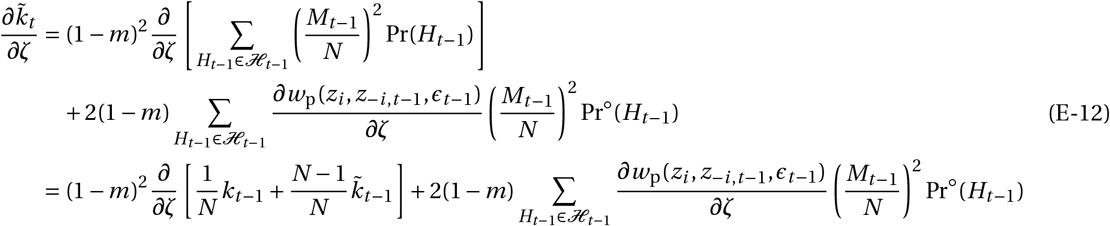

using *k*_*t* −1_ and 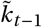 defined in eq. (E-2) and eq. (E-3).

Before substituting eq. (E-12) into eq. (E-4), note first that the first term of eq. (E-4) can be expanded as

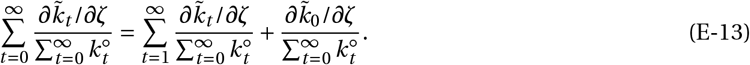

However since the mutation arises as a single copy at generation *t* = 0, the second term of eq. (E-13) vanishes (i.e. 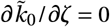), leaving us with

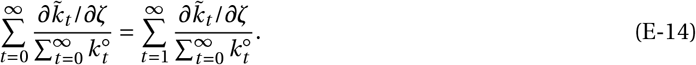

Substituting eq. (E-12) into the right hand eq. (E-14), we obtain

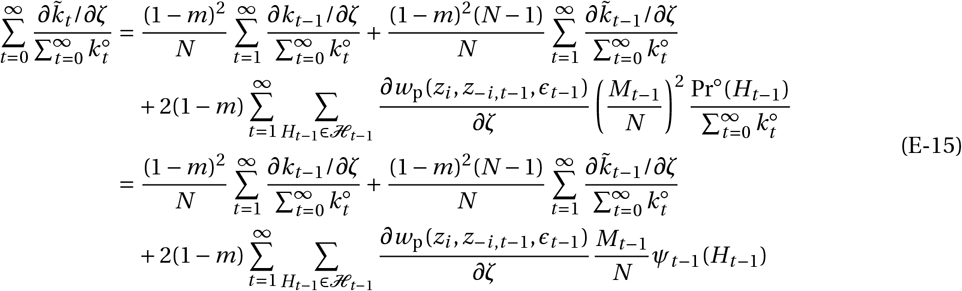

using eq. (A-3). Changing the dummy variable *t* on the right hand side of eq. (E-15) and using eq. (B-4) lead us to

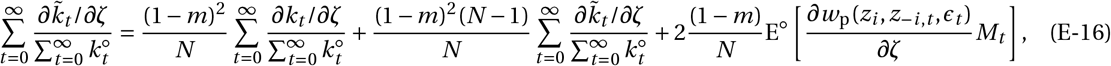

with E^◦^[*f* (*H*_*t*_)] defined in eq. (B-4). Re-arranging eq. (E-16) and using eq. (D-2), we finally obtain

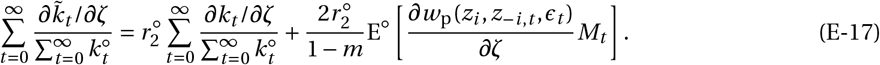

Plugging eq. (E-17) into eq. (E-4) gives

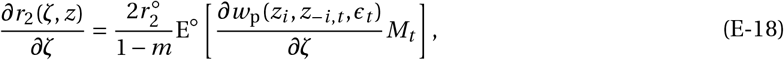

into which we substitute

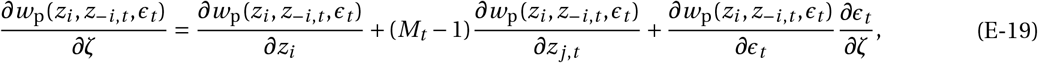

leaving us with

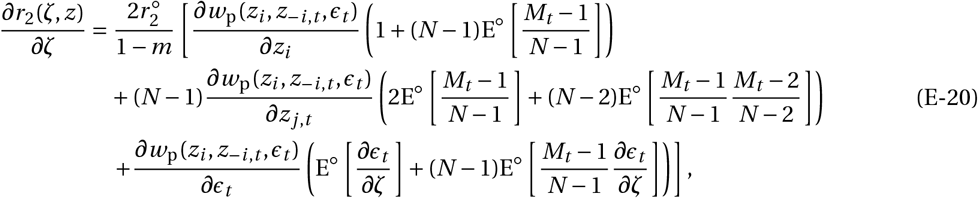

after some straightforward re-arrangements. Using eqs. (B-13) and (C-14), this eq. (E-20) simplifies to

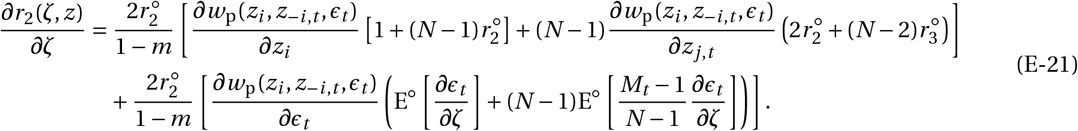

The first line of eq. (E-21) corresponds to the effect of selection on intra-generational relatedness in the absence of ecological inheritance (eq. B.46 in Wakano and Lehmann, 2014) while the second is due to such inheritance.

We can readily extend eq. (E-21) to the multi-dimensional case as required by eq. (15) of the main text. Specifically, we use to eq. (E-21) to get the effect of trait *p* on relatedness,

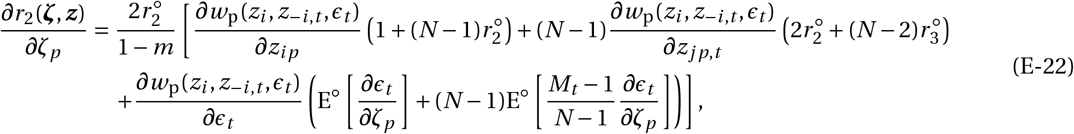

where E^◦^ [*∂ϵ*_*t*_ /*∂ζ*_*p*_] and E^◦^ [(*M*_*t*_ − 1)/(*N* − 1) × *∂ϵ*_*t*_ /*∂ζ*_*p*_] are given in eq. (B-20) and (C-38), respectively. Substituting those into eq. (E-22) gives row 9 of Table 1.

#### E.2 Inter-generational relatedness

We now turn to the effects of selection on inter-generational relatedness, 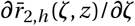, which appears in eq. (C-44) in Appendix C.3.3 (and in eq. 22 of the main text as 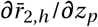 for the effect of trait *p* on relatedness). Our derivation follows broadly the same argument as the one used in Appendix E.1 but is more involved.

As a starting point, we plug eq. (A-3) into eq. (C-44) to write

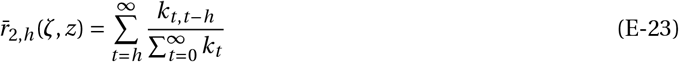

where *k*_*t*_ is defined in eq. (E-2) and

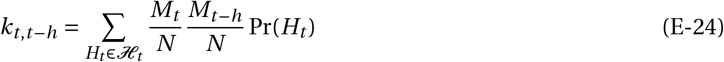

is the probability that two individuals randomly sampled from a mutant patch, one at generation *t* and the other at generation *t* − *h*, are both mutants. Like *k*_*t*_, *k*_*t, t* −*h*_ is a function of *ζ* and *z*. Taking the derivative of eq. (E-23) with respect to *ζ* and evaluating it at *z* obtains,

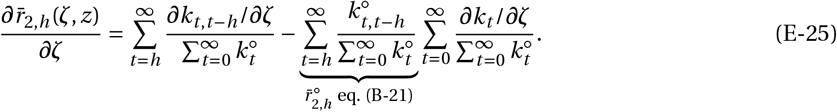

It will be convenient to define

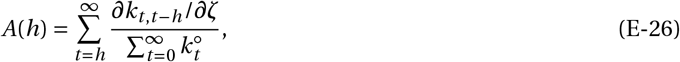

so as to write

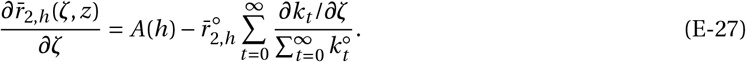

To specify *A*(*h*), we first use conditional probabilities to re-write eq. (E-24) as

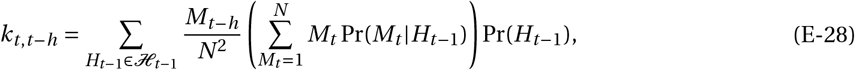

into which we substitute eq. (E-10) to get

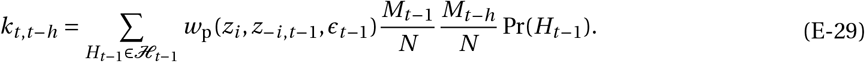

Taking the derivative of eq. (E-29) and using eqs. (E-8) and (E-24) then gives

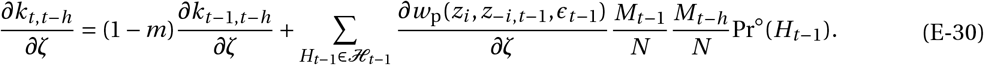

Substituting eq. (E-30) into eq. (E-26) we obtain

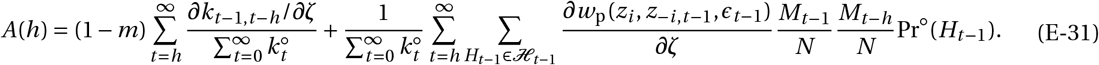

A change of dummy variable *t* allows us to write eq. (E-31) as

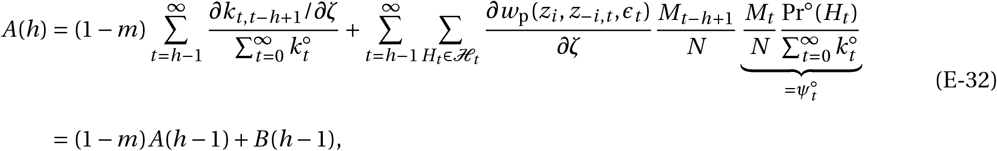

where we used eq. (E-26) and defined

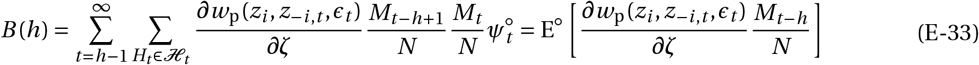

(using eq. B-4).

Eq. (E-32) is a linear (non-homogeneous) recurrence for *A*(*h*) in *h*. Solving the recurrence using standard methods yields

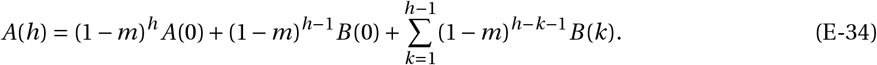

We now specify in turn *A*(0) and *B* (*k*) (for *k* = 0 and *k* > 0). For *A*(0), we first note from eq. (E-26) that

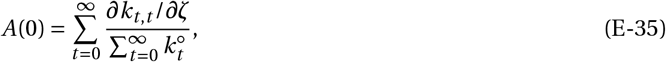

where by definition (eq. E-24), *k*_*t, t*_ is the probability that two individuals randomly sampled with replacement in a mutant patch at generation *t* are both mutants. This probability can be decomposed as

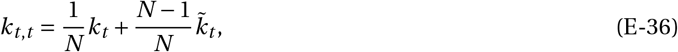

using eq. (E-2) and eq. (E-3). Thus eq. (E-35) becomes,

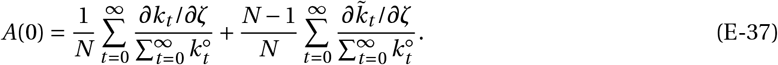

Substituting eq. (E-17) into eq. (E-37), we get

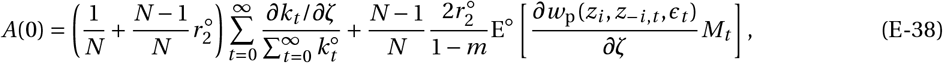

which can be re-written as

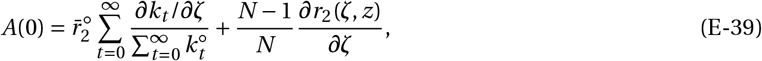

using eqs. (D-3) and (E-18).

For *B* (0), we see from eq. (E-33) that it is

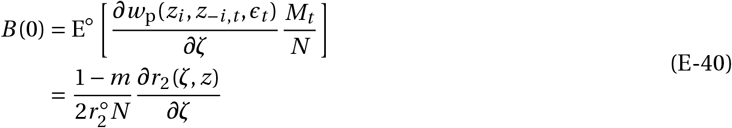

(using eq. E-18).

For *B* (*k*) (with *k* ≥ 1), we first substitute eq. (E-19) into eq. (E-33) to obtain

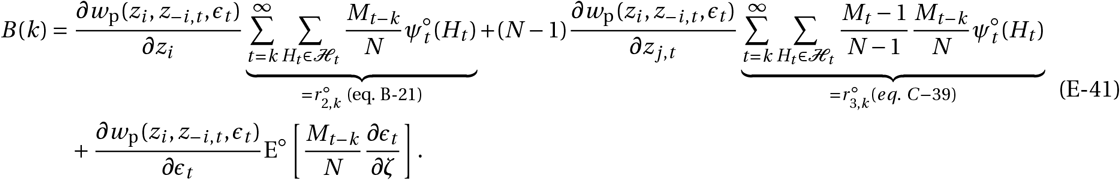

The last term of eq. (E-33) can be computed using eqs. (B-4) and (B-18):

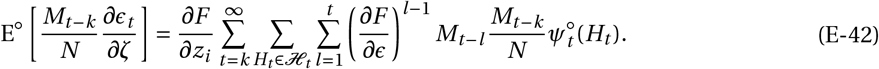

Since the mutant appeared at *t* = 0, we have *M*_*t* −*l*_ *M*_*t* −*k*_ = 0 for any *t* < *l* or *t* < *k*. We can thus re-arrange the order of the summation in eq. (E-42) to obtain

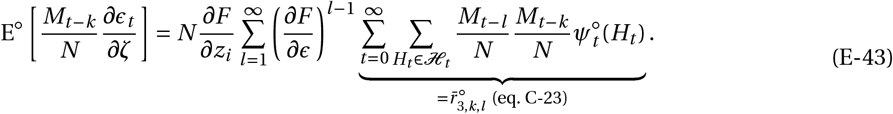

Substituting eq. (E-43) into eq. (E-41) we thus get

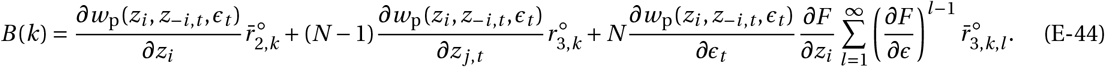

We can then substitute eqs. (E-39), (E-40) and (E-44) into eq. (E-26), yielding

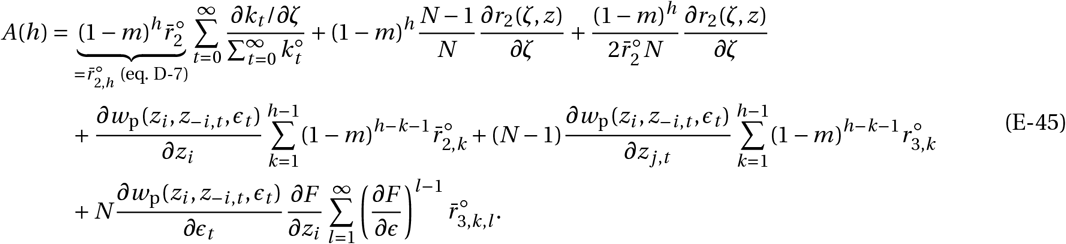

Finally, plugging eq. (E-45) into eq. (E-27), we obtain after some straightforward re-arrangements

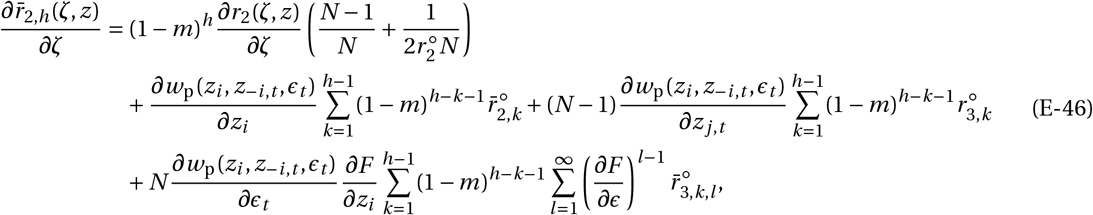

as required to compute eq. (C-44). Eq. (E-46) is readily extended to the multi-trait case (i.e. for eq. (22) of the main text) and to give the effect of a trait *p* on inter-generational relatedness,

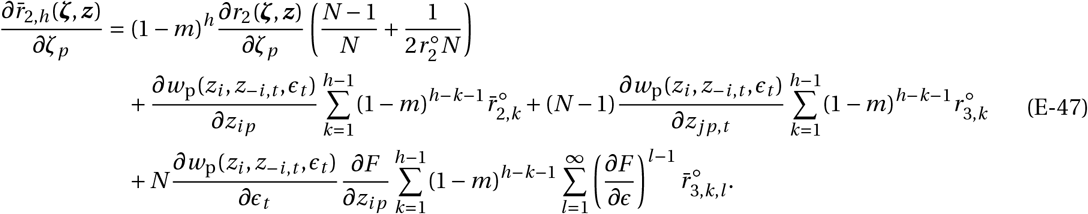

For computational purposes (and be able to use 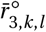 as calculated in eq. D-19), we re-arrange the last term of eq. (E-47) such that 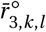 has either *k* = *l* or *l* > *k*:

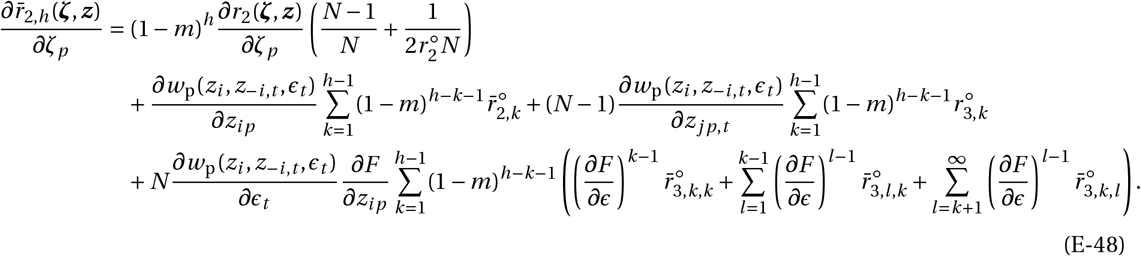

Substituting for the relevant inter-generational relatedness coefficients found in Appendix D into eq. (E-48), which is in turn substituted into eq. (22) of the main text (or equivalently eq. C-48 in Appendix C.3.3), we obtain row 10 of Table 1 in the main text.

### F Coevolution of dispersal and attack rate on a local resource

In this appendix, we describe our analysis behind the results presented in section 4 of the main text. Our computations can be followed from the accompanying Mathematica notebook (found in Supplementary Files).

#### F.1 Ecological dynamics

We first characterise the equilibrium 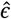 of the resource system when the consumer population is monomorphic. Substituting eq. (29) into eq. (2), we find a single non-zero equilibrium

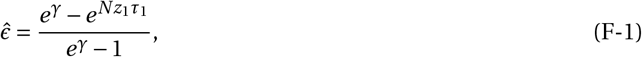

which is stable (i.e. satisfies eq. 3) when

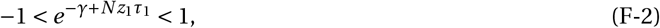

or equivalently when

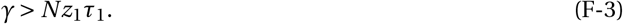

#### F.2 Evolutionary dynamics under directional selection

First, we investigate directional selection on attack rate and dispersal.

##### F.2.1 Directional selection on attack rate

Substituting eqs. (29)-(33) into eq. (10), we find that we can express the selection gradient on the attack rate as

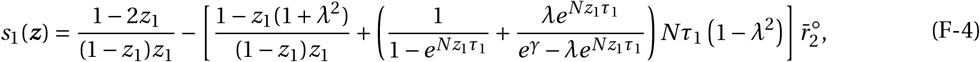

where *λ* = 1 − *m*(***z***) (with *m*(***z***) given eq. 34). Re-arranging the condition that *s*_1_(***z***^∗^) = 0 for singular strategy 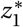 from eq. (F-4) leads to eq. (36) of the main text. This equation cannot be solved analytically. Note however that: (i) the selection gradient decreases monotonically with *z*_1_:

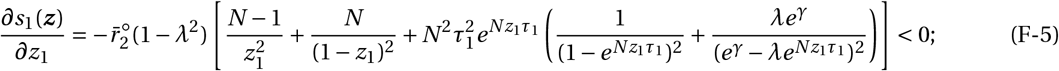

and (ii) that at its boundaries, 0 < *z*_1_ < *z*_max_ = max(1,(*γ* − ln(*λ*))/(*Nτ*_1_)) (from eq. F-3), the selection gradient is positive and negative, respectively:

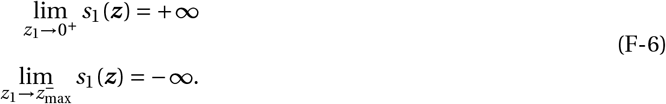

Using the intermediate value theorem, we can deduce from (i) and (ii) that there exists a unique singular strategy 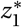 and that this strategy is convergence stable (when the attack rate evolves in isolation).

##### F.2.2 Directional selection on dispersal

The selection gradient on dispersal is found to be as in previous works that have studied it for a Wright-Fisher life-cycle under the island model of dispersal (e.g. Ajar, 2003), namely,

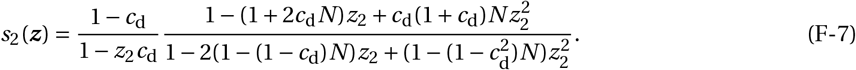

This gradient vanishes at the singular strategy,

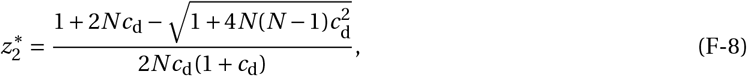

which is such that

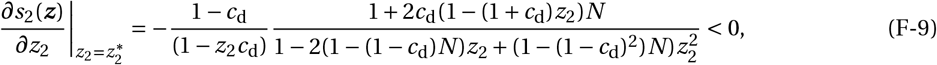

meaning that 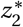 is convergence stable (when dispersal evolves in isolation).

##### F.2.3 Convergence stability when attack rate and dispersal coevolve

The above shows that when either the attack rate or dispersal evolve independently, the population converges to a singular strategy for either trait. When they coevolve, whether the joint singular strategy 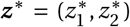 (such that ***s***(***z***^∗^) = **0**) is convergence stable depends on the leading eigenvalue of the Jacobian matrix (eq. 8). However, since we have from eq. (F-7) that:

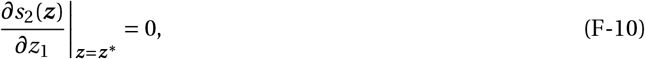

i.e. that one of the off-diagonal entry of *J* (***z***^∗^) is zero, the eigenvalues of *J* (***z***^∗^) are in fact equal to its diagonal entries. From section F.2.1 and F.2.2, we know that these diagonal elements are both negative. Hence the joint singular strategy ***z***^∗^ given by eq. (36) of the main text and eq. (F-8) is also convergence stable when both traits coevolve.

#### F.3 Disruptive selection and polymorphism

Under directional selection, the population gradually converges to 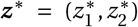. Whether it remains monomorphic for ***z***^∗^ or becomes polymorphic depends on the Hessian matrix **H**(***z***^∗^) (eq. 12). With two traits coevolving, this Hessian is

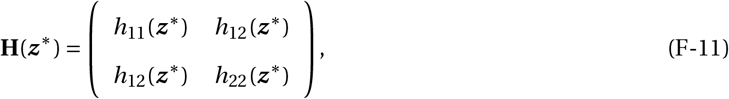

where *h*_11_(***z***^∗^) is the disruptive selection coefficient on the attack rate, *h*_22_(***z***^∗^) the disruptive selection coefficient on dispersal and *h*_12_(***z***^∗^) the correlational selection coefficient on both traits. We compute the elements of the Hessian matrix by substituting eqs. (29)-(33) into eqs. (14)-(21) and using Table 1 in the accompanying Mathematica Notebook (see Supplementary Files), summarising our findings below.

##### F.3.1 Stabilising selection on dispersal

In line with previous studies (Ajar, 2003), we find that the disruptive selection coefficient on dispersal can be expressed as

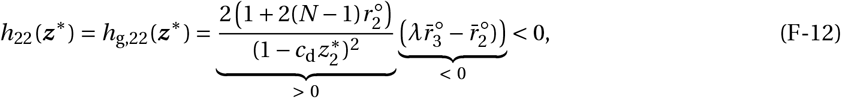

which is always negative. Selection on dispersal alone is thus always stabilising in this model.

##### F.3.2 Stabilising and disruptive selection on attack rate

Because we cannot solve for the singular attack rate 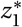 analytically (given by eq. (36) of the main text), we performed a numerical analysis of the remaining entries of the Hessian matrix, *h*_11_(***z***^∗^) and *h*_12_(***z***^∗^) over a large number of sets of parameters (280’000 sets, Supplementary Mathematica Notebook for analysis). This analysis reveals that typically *h*_11_(***z***^∗^) < 0, i.e. selection on the attack rate is typically stabilising (white and light grey region in Fig. 2B), *h*_11_(***z***^∗^), except for values of dispersal cost *c*_d_ and renewal *γ* close to their threshold values below which the resource goes extinct (dark grey region in Fig. 2B).

##### F.3.3 Positive correlational selection on the attack rate and dispersal

For all sets of parameters we tested, we found that *h*_12_(***z***^∗^) > 0, i.e. correlational selection correlational selection on the attack rate and dispersal is positive, thus favouring a positive association between these two traits. Our numerical analysis also reveals that the different components of correlational selection (decomposed as eqs. 14 and 16 of the main text) have the following signs,

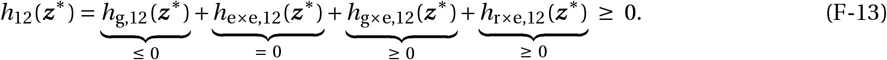

Additionally, as *h*_r×e,12_(***z***^∗^) is typically much larger than *h*_g,12_(***z***^∗^) + *h*_g×e,12_(***z***^∗^), our analysis shows that the main driver for a positive correlational selection between dispersal and attack rate is biased ecological inheritance. This contrasts with the social polymorphism reported in Mullon et al. (2018), which is driven by intra-generational effects only (i.e. by *h*_g,12_(***z***^∗^), which here in fact is negative).

From eqs. (21) and (22) of the main text, *h*_r×e,12_(***z***^∗^) for our model can be expressed as

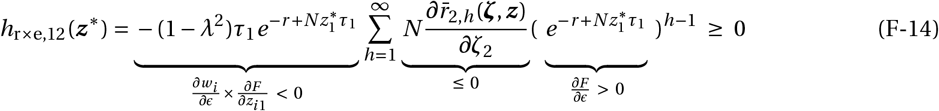

(see Mathematica Notebook for derivations of the signs of the different quantities). This shows that correlational selection is positive owing to the negative effect of the attack rate on the local resource and thus on fitness of individuals in the patch in the future (*∂w*_*i*_ /*∂ϵ* × *∂F* /*∂z*_*i* 1_ < 0), and the negative effect of dispersal on relatedness with such individuals 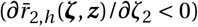. In other words, selection favours a positive association among attack rate and dispersal as it ensures that individuals that consume more do not do so at the expense of future relatives.

Computing the eigenvalue of the Hessian matrix **H**(***z***^∗^) numerically reveals that the range of parameters under which this eigenvalue is positive is greater than the range under which *h*_11_(***z***^∗^) > 0 (e.g. compare total gray region with dark gray region in Fig. 2B). This shows that correlational selection *h*_12_(***z***^∗^) favours polymorphism in this example.

#### F.4 Individual based simulations

To check our mathematical analysis about the emergence of polymorphism and see the long term outcome of this polymorphism, we ran individual-based simulations using C++ (version 11, Supplementary Files for code). Unlike our invasion analysis which assumes an infinite number of patches and only two alleles segregating, we track evolution in a population with a large but finite number *N*_p_ of patches, and where mutations are rare but not so rare that only one mutant can segregate at a time. Such simulations have been shown to give reproducible results that agree very well with analytical results from local analyses in group-structured populations (e.g. Wakano and Lehmann, 2014; Mullon et al., 2018; Mullon and Lehmann, 2019, 2022; Schmid et al., 2022).

Our algorithm follows exactly the life-cycle described in main text section 4. The population is subdivided among *N*_p_ patches, each carrying *N* individuals (see figure legends for parameter values). Each individual *i* ∈ {1,…, *N* } in each patch *j* ∈ {1,…, *N*_p_} is characterized by a vector of two quantitative traits: (*z*_1,*i j*_, *z*_2,*i j*_), with attack rate *z*_1,*i j*_ and dispersal probability *z*_2,*i j*_ ; and each patch by its resource abundance *ϵ*_*j*_. The parental population thus consists in an ordered collection of trait values of size 2 × *N* × *N*_p_, which we can collect into 𝒞 = {(*z*_1,*i j*_, *z*_2,*i j*_)_*i, j*_ } and in a 1 × *N*_p_ vector of patch states: (*ϵ*_*j*_)_*j*_. We update these two objects in four steps at each generation:

i. At the beginning of each generation, we calculate the fecundity *f*_*i j*_ of each individual *i* in each patch *j* according eq. (31), i.e. as

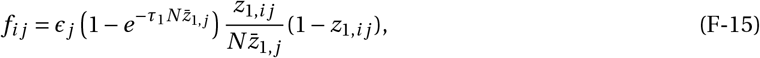

where 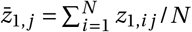 is the average attack rate in patch *j*.
ii. We form the offspring generation. We fill each patch *k* ∈ {1,…, *N*_p_} by sampling *N* individuals from the parental generation 𝒞 using multinomial sampling (i.e. with replacement). Each parent is weighted according to fecundity and according to the patch they live in relative to the one which is being filled to model dispersal. Specifically, when filing patch *k* ∈ {1,…, *N*_p_}, each (*i, j*) element of 𝒞 is weighted by

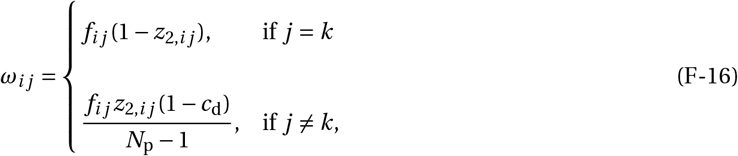

where the factor 1/(*N*_p_ − 1) captures the probability that a dispersing offspring lands in a specific patch.
iii. Mutation occurs. Independently from one another, each offspring mutates with probability *μ* = 0.01, in which case its phenotype consists of its parental phenotype to which we add a perturbation sampled from a bivariate Normal distribution with mean (0, 0) and covariance matrix

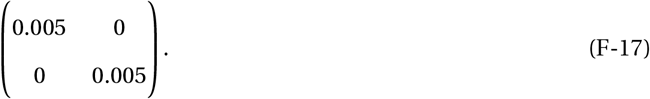

The resulting phenotypic values are truncated such that the attack rate remains positive and the dispersal probability remains between 0 and 1.
iv. We update the resource abundance *ϵ*_*j*_ in each patch *j* according to eq. (29), i.e. as

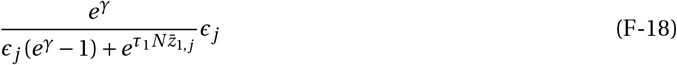

where 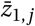 is the average attack rate in patch *j* in the parental generation (i.e. from 𝒞) and recall that *γ* = *γ*_0_(*T* − *τ*_1_) is per-generation renewal rate.

We iterate these four steps for 100^′^000 generations.

## Notes

### Competing Interest Statement

The authors have declared no competing interest.

### Summary of Updates

Added results where demography is stochastic (Fig S6).

